# Scaling of information in large neural populations reveals signatures of information-limiting correlations

**DOI:** 10.1101/2020.01.10.902171

**Authors:** MohammadMehdi Kafashan, Anna Jaffe, Selmaan N. Chettih, Ramon Nogueira, Iñigo Arandia-Romero, Christopher D. Harvey, Rubén Moreno-Bote, Jan Drugowitsch

## Abstract

How is information distributed across large neuronal populations within a given brain area? One possibility is that information is distributed roughly evenly across neurons, so that total information scales linearly with the number of recorded neurons. Alternatively, the neural code might be highly redundant, meaning that total information saturates. Here we investigated how information about the direction of a moving visual stimulus is distributed across hundreds of simultaneously recorded neurons in mouse primary visual cortex (V1). We found that information scales sublinearly, due to the presence of correlated noise in these populations. Using recent theoretical advances, we compartmentalized noise correlations into information-limiting and nonlimiting components, and then extrapolated to predict how information grows when neural populations are even larger. We predict that tens of thousands of neurons are required to encode 95% of the information about visual stimulus direction, a number much smaller than the number of neurons in V1. Overall, these findings suggest that the brain uses a widely distributed, but nonetheless redundant code that supports recovering most information from smaller subpopulations.

## Introduction

Our brains encode information about sensory features and other variables of interest in the activity of large neural populations. The amount of encoded information provides an upper bound on behavioral performance, and so exposes the efficiency and structure of the computations implemented by the brain. The format of this encoding reveals how downstream brain areas ought to access the encoded information for further processing. For example, the amount of information in visual cortex about the drift direction of a moving visual stimulus determines how well one could in principle discriminate different drift directions if the brain operates at maximum efficiency, and its format tells us how downstream motion-processing areas ought to “read out” this information. Therefore, knowing how the brain encodes information about the world is necessary if we are to understand the computations it performs. Unfortunately, we still know little about how information is distributed across neuronal populations even within a single brain area. Is information spread evenly and largely independently across neurons, or in a way that introduces significant redundancy? In the first scenario, one would need to record from the whole neuronal population to get access to all available information, whereas in the second scenario only a fraction of neurons would be needed.

The amount of information about a stimulus feature that can be extracted from neural population activity depends on how this activity changes with a change in the stimulus feature. For information that can be extracted by a linear decoder, which is the information we focus on in this work, it depends on the neurons’ tuning curves, as well as how their activity varies across repetitions of the same stimulus (i.e., “noise”) (Averbeck, Latham, & Pouget, 2006; Kohn, Coen-cagli, Kanitscheider, & Pouget, 2016; Nogueira et al., 2019; Shamir, 2014). Due to the variability in neural responses to repetitions of the same stimulus, each neuron’s response provides limited information about the stimulus feature (Carandini, 2004; Faisal, Selen, & Wolpert, 2008; Shadlen & Newsome, 1998; Softky & Koch, 1993; Tolhurst, Movshon, & Dean, 1983). If the noise is independent across neurons, it can be averaged out by pooling across neurons (Zohary, Shadlen, & Newsome, 1994), and total information would on average increase by the same amount with every neuron added to this pool (**Fig. 1a**, red). This corresponds to the first scenario in which information is spread evenly across neurons. If, however, the trial-to-trial variations in spiking are shared across neurons – what are referred to as “noise correlations” – the situation is different. In general, depending on their structure, noise correlations can either improve or limit the amount of information (**Fig. 1b**), such that the presence of correlated noise alone does not predict its impact. In a theoretical population with translation-invariant tuning curves (i.e., the individual neurons’ tuning curves are shifted copies of each other) and noise correlations that are larger for neurons with similar tuning, information might quickly saturate with population size (Abbott & Dayan, 1999; Zohary et al., 1994), corresponding to the second scenario (**Fig. 1a**, black). Even though such a correlation structure has been observed across multiple brain areas (Adibi, McDonald, Clifford, & Arabzadeh, 2013; Averbeck & Lee, 2003; Gu et al., 2011; Maynard et al., 1999; Zohary et al., 1994), neural tuning is commonly more heterogeneous than assumed by Zohary et al. (1994). A consequence of this heterogeneity is that information might grow without bound even with noise correlations of the aforementioned structure (Ecker, Berens, Tolias, & Bethge, 2011). Overall, it remains an open question if information saturates in large neural populations of human and animal brains (Kohn et al., 2016).

**Figure 1.**
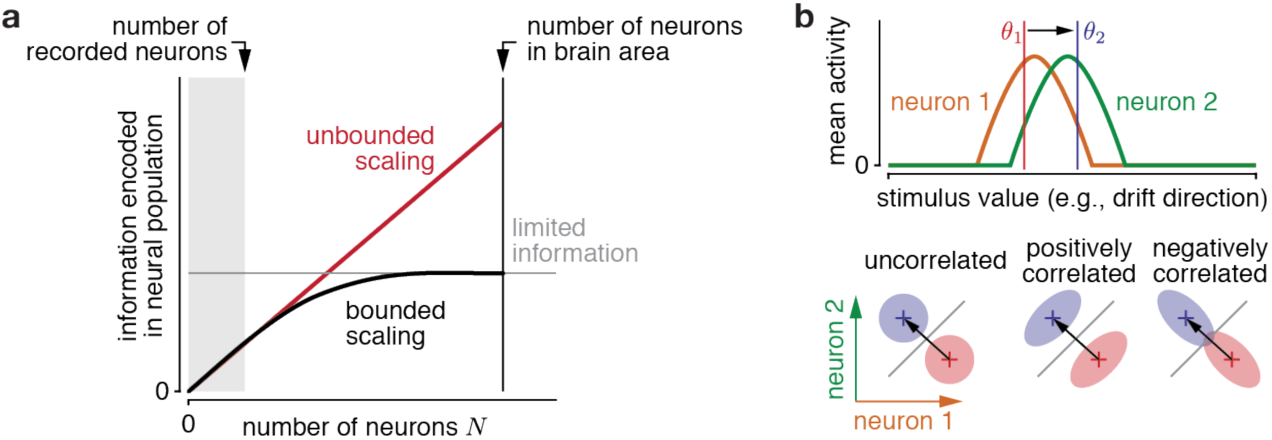
Information scaling in large neural populations, and the impact of noise correlations on information. **a.** The information that a population of neurons can encode about some stimulus value is always a non-decreasing function of the population size. Information might on average increase with every added neuron (unbounded scaling; red) if the information is evenly distributed across all neurons. In contrast, information can rapidly saturate if information is redundant, and thus it is not strictly limited by population size, but by other factors. In general, it has only been possible to record from a very small subset of neurons of a particular area (grey shaded), from which it is hard to tell the difference between the two scenarios if the sampled population size is too small. **b.** The encoded information is modulated by noise correlations. This is illustrated using two neurons with different tunings to the stimulus value (top). The amount of information to discriminate between two stimulus values (*θ*_1_/red and *θ*_2_/blue) depends on the difference in mean population activity (crosses) between stimuli, and the noise correlations (shaded ellipsoids) for either stimulus (bottom, showing joint neural activity of both neurons). The information is largest when the noise is smallest in the direction of the mean population activity difference (black arrow), which leads to the largest separation across the optimal discrimination boundary (grey line). In this example, positive correlations boost information (middle), whereas negative correlations lower it (right), when compared to uncorrelated neurons (left). In general, the impact of noise correlations depends on how they interact with the population’s tuning curves.

If information saturates in such populations, then, by the theory of information-limiting correlations (TILC) (Moreno-Bote et al., 2014), information in large populations is limited exclusively by one specific component of the noise correlations. This component introduces noise in the direction of the change of the mean population activity with stimulus value (e.g., drift direction; black arrow in **Fig. 1b** bottom), thus limiting information about this value. Measuring this noise correlation component directly in neural population recordings is difficult, as noise correlations are, in general, difficult to estimate well (Cohen & Kohn, 2011), and the information-limiting component is usually swamped by other types of correlation that do not limit information (Kanitscheider, Coen-Cagli, & Pouget, 2015; Moreno-Bote et al., 2014). Fortunately, however, TILC also predicts *how* information scales with population size if information-limiting correlations are present. We thus exploited this theory to detect the presence of information-limited correlations indirectly by examining how information scales with population size.

To search for the presence of information-limiting correlations, we simultaneously recorded the activity of hundreds of neurons in V1 of awake mice in response to drifting gratings, with hundreds of repeats of each stimulus. We asked how these neurons encoded information about the direction of the moving visual stimulus. We found that noise correlations reduce information even within the limited neural populations we could record. Applying TILC to compartmentalize information-limiting correlations from nonlimiting correlations, and to extrapolate the growth of information to larger neural populations, we found that on the order of tens of thousands of neurons would be required to encode 95% of the information about the direction of the moving stimulus. Given that there are hundreds of thousands of neurons in this brain region, this means that only a small fraction of the total population is needed to encode this information. This is not because only a small fraction of neurons contains information about the stimulus; rather, we found that most neurons contain information about the stimulus, but because information is represented redundantly, only a small fraction of these neurons is actually needed. Notably, the size of the required neural population depends only weakly on stimulus contrast; thus, increasing the amount of information in this brain area does not substantially increase the number of neurons required to encode 95% of the information about the stimulus. Finally, we found that the low-dimensional neural subspace that captures a large fraction of the noise correlations does not encode a comparably large fraction of information. Overall, our results suggest that information in mouse V1 is both highly distributed and highly redundant, which is true regardless of the total amount of information encoded.

## Results

To measure how information scales with population size, we used two-photon calcium imaging to record neural population activity from layer 2/3 of V1 in awake mice observing a low-contrast drifting grating (10% contrast). The drift direction varied across trials, with each trial drawn pseudorandomly from eight possible directions, spaced evenly around the circle (**Fig. 2a**). We simultaneously recorded 273-386 neurons (329 on average) across 4 mice and a total of 16 sessions (**Fig. 2b**), and analyzed temporally deconvolved calcium activity, summed up over the stimulus presentation period as a proxy for their spike counts within that period. The tuning curves of individual neurons (**Fig. 2c**) revealed that, on average, only a small fraction of neurons (5%-45% across mice/sessions, 18% average) were tuned to the grating’s drift direction, while a larger fraction of neurons (38%-60% across mice/sessions, 48% average) were sensitive to the grating’s orientation, but not its direction of drift. The remaining neurons had no appreciable tuning (14%-52% across mice/sessions, 34% average), but were nonetheless included in the analysis, as they can contribute to the information that the population encodes through noise correlations (Leavitt, Pieper, Sachs, & Martinez-Trujillo, 2017; Pruszynski & Zylberberg, 2019).

**Figure 2.**
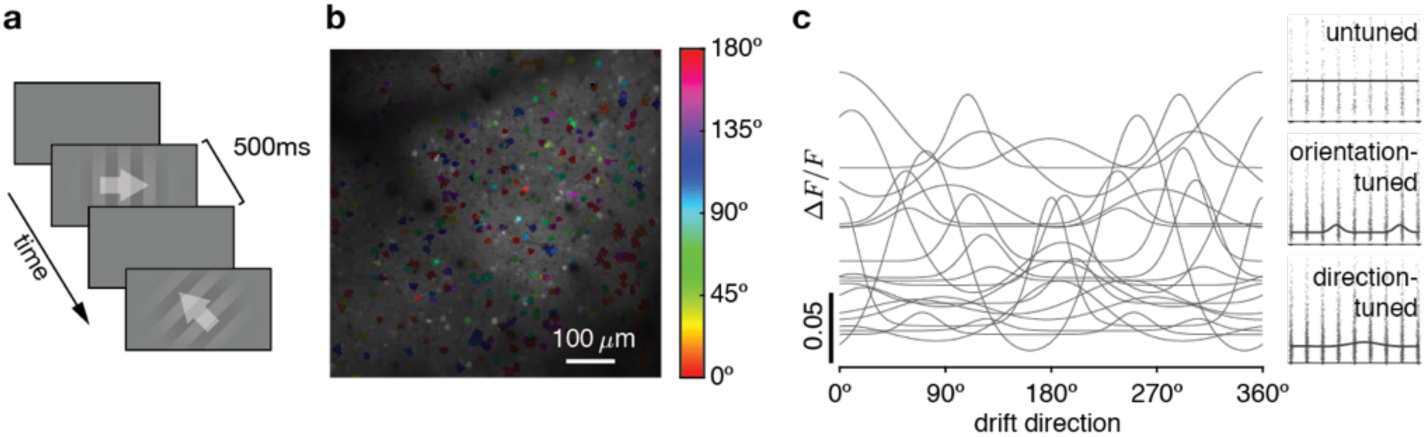
Experimental design, population recordings, and neural tuning. **a.** Mice passively observed sequences of drifting gratings (white arrows overlaid for illustration only), interleaved with blank screens. **b.** Example field-of-view with significantly tuned neurons color coded by their preferred orientation tuning. **c.** Left: example fitted tuning curves of 20 significantly tuned neurons. Right: example tuning curves (line) fitted to per-trial neural responses (dots, horizontally jittered) for an untuned (top), orientation-tuned (middle) and direction-tuned (bottom) neuron.

### Noise correlations limit information

To quantify stimulus information encoded in the response of neural populations, we asked how well a linear decoder of the recorded population activity (i.e., information decodable by a single neural network layer) would allow us to discriminate between a pair of drift directions (**Fig. 3a**). We measured the decoder’s performance by generalizing linear Fisher information, usually restricted to fine discriminations, to coarse discrimination (**Fig. 3b**). This generalization is closely related to the sensitivity index *d*′ from signal detection theory (Green & Swets, 1966; Nogueira et al., 2019), and has a set of appealing properties (see Methods). In particular, combining the activity of two uncorrelated neural populations causes their associated Fisher information to add, so that it does not trivially saturate like other measures of discrimination performance (**Fig. 3c**, inset).

**Figure 3.**
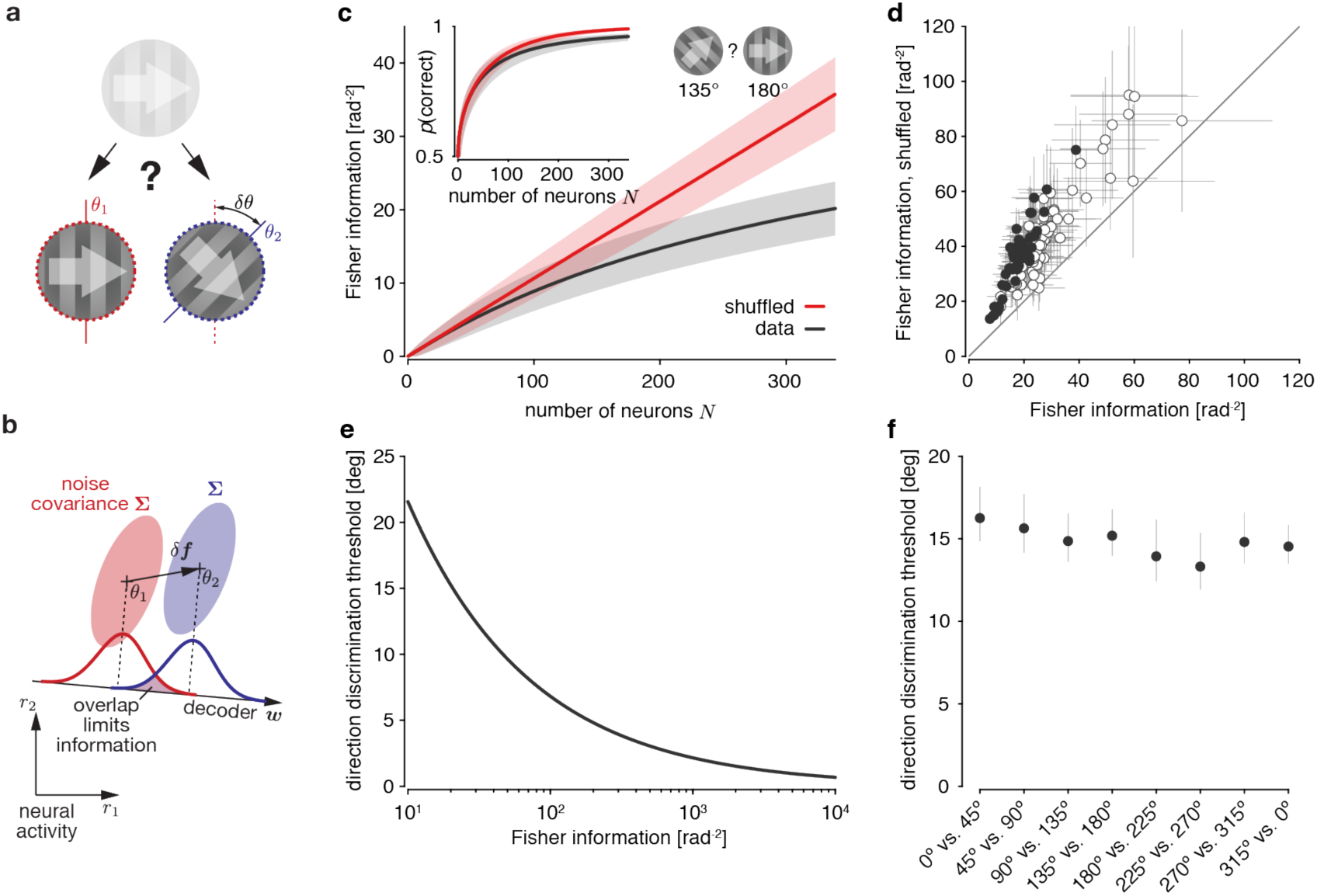
Noise correlations limit information across all drift directions. **a.** A drift direction discrimination task, in which a hypothetical observer needs to judge which of two template drift directions (*θ*_1_ or *θ*_2_; indicated by white arrows) an observed low-contrast drifting grating corresponds to. **b.** Mean activity ***f*** (crosses) and noise covariance Σ (shaded area ≈ 2SDs) of a pair of neurons across repeated presentation of the same two drift directions, *θ*_1_ (red) and *θ*_2_ (blue). Linear information about drift direction is limited by the projection of the noise onto the optimal linear decoder ***w***. This decoder depends on how mean activity changes with drift direction (*δ****f*** = ***f***(*θ*_2_) − ***f***(*θ*_1_)) and the noise covariances Σ. **c.** The information associated with discriminating between drift directions 135**°** and 180**°** scales sublinearly with population size (black; mean +/- 1SD across random orderings of neurons within the population). If we remove noise correlations by shuffling trials across neurons, the information scales linearly (red). This linear growth would not be apparent from the probability of correctly identifying the stimulus’ drift direction (inset), which is monotonically, but non-linearly related to Fisher information, and saturates in both cases. **d.** Information in the recorded population was consistently larger for trial-shuffled data across different discriminations, sessions, and mice. Each dot (mean +/- 1SD of information estimate; filled = significant increase, bootstrap, *p* < 0.05) shows the information estimated for one discrimination with *δθ* = 45°. **e.** The drift direction discrimination threshold (corresponding to 80% correct discriminations) we would expect to see in a virtual discrimination experiment drops with the amount of information that V1 encodes about drift directions. **f.** The inferred drift direction discrimination threshold for the same session as in panel **c** is comparable across the different drift direction pairs with *δθ* = 45° used to estimate Fisher information with the recorded population.

We used generalized Fisher information to measure how information about drift direction scales with the number of neurons in the recorded population. Because this scaling depends on the order in which we add particular neurons to the population (individual neurons might contribute different amounts of additional information to a population), we measured average scaling by averaging across a large number of different random orderings (see Methods). **Figure 3c** shows this average scaling for one example session for discriminating between drift directions of 135° and 180° (arbitrary choice; as shown below, other drift direction combinations resulted in comparable information scaling). Information increases with population size, but, on average, additional neurons contribute less additional information to larger populations than to smaller ones. The resulting sublinear scaling is expected if noise correlations limit information. Indeed, trial-shuffling the data to remove pairwise correlations resulted in information that scaled linearly, with average information exceeding that of the non-shuffled data for all population sizes (except, trivially, for single neurons), and a significantly higher total information within the recorded population (bootstrap, *p* ≈ 0.0062). Such linear scaling was not apparent if we measured discrimination performance by the fraction of correct discriminations (**Fig. 3c**, inset), illustrating the point that Fisher information is indeed a better measure to analyze information scaling. Removing noise correlations resulted in a significant information increase in all our datasets (**Fig. 3d**; paired *t*_63_ = −17.93, two-sided *p* ≈ 1.96 × 10^−26^; statistics computed across all sessions and mice, but only across non-overlapping *δθ* = 45°discriminations to avoid duplicate use of individual drift direction trials; see **Table S1** for avg. per-neuron information for all sessions/mice), confirming that noise correlations indeed limit information in our recorded populations.

To aid interpretation of the estimated amounts of Fisher information, we translated them into quantities that are more frequently measured in experiments. Specifically, we assumed that the recorded neural population was used to discriminate between two close-by drift directions in a virtual fine discrimination task (similar to **Fig. 3a**). For a given estimate of Fisher information, we could then determine the expected discrimination threshold at which the ideal observer could correctly discriminate between two drift directions in 80% of the trials based solely on neuronal responses (**Fig. 3e**). This resulted in a discrimination threshold of ∼15.2° for the Fisher information estimated from a 135° vs. 180° discrimination (**Fig. 3f**). Previous work has shown that attending to a stimulus boosts the information encoded about this stimulus (Ni, Ruff, Alberts, Symmonds, & Cohen, 2018; Otazu, Tai, Yang, & Zador, 2009). As our animals were passive observers that were not actively engaged in any task, the estimated threshold is most likely a significant overestimate (i.e., an underestimate of Fisher information). Nonetheless, it provides a reasonable interpretation of the information encoded in the recorded population. Computing the discrimination threshold for all drift direction pairs with *δθ* = 45° resulted in comparable thresholds that did not differ significantly (bootstrap, two-sided *p* ≈ 0.50 for session shown in **Fig. 3f**, two-sided *p* > 0.49 for all sessions/mice). We found comparable information across all drift directions, confirming that we recorded from populations that were homogeneously tuned across all drift directions.

### Neural signatures of limited asymptotic information

To identify neural signatures of limited encoded information, we relied on the theory of information-limiting correlations (TILC) that showed that noise correlations in large populations can be compartmentalized into information-limiting and non-limiting components (Moreno-Bote et al., 2014). The limiting component is scaled by the inverse of the asymptotic information *I*_∞_, which is where information asymptotes in the limit of a large number of neurons (Kanitscheider, Coen-Cagli, & Pouget, 2015; Moreno-Bote et al., 2014). This compartmentalization allowed us to split the information *I*_*N*_ in a population of *N* neurons into the contribution of limiting and non-limiting components (see Methods), resulting in

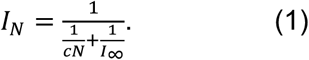

This expression assumes that the non-limiting component contributes *c* information per neuron on average, irrespective of the current population size. Model comparison to alternative non-limiting component scaling models confirmed that this assumption best fits our data (**Fig. S1b**).

Increasing the population size *N* in Eq. (1) reveals how information ought to scale in small populations if it is limited in large populations (**Fig. 1**). Information would initially grow linearly, closely following *cN*. However, for sufficiently large *N*, it would start to level off and slowly approach the asymptotic information *I*_∞_. If we were to record from a small number of neurons, we might only observe the initial linear growth and would wrongly conclude that no information limit exists (**Fig. 1**). Therefore, simultaneously recording from sufficiently large populations is important to identify limited asymptotic information.

To distinguish between a population in which information does not saturate from one in which it does, we fitted two models to the measured information scaling. The first assumed that, within the recorded population, information scales linearly and without bound. We might observe this information scaling if, on average, each neuron contributes the same amount of information. The second model corresponds to Eq. (1), and assumes that information asymptotes at *I*_∞_. Our fits relied on a large number of repetitions (at least as many as the number of recorded neurons) of the same drift direction within each experimental session to ensure reliable, bias-corrected information estimates (Kanitscheider, Coen-Cagli, Kohn, & Pouget, 2015). These estimates are correlated across different population sizes, as estimates for larger populations share data with estimates for smaller populations. Unlike previous work that estimated how information scales with population size (Cotton et al., 2018; Mendels & Shamir, 2018), we accounted for these correlations by fitting how information increases with each additional neuron, rather than fitting the total information for each population size. This information increase turns out to be statistically independent across population sizes (see Methods), making the fits statistically sound.

**Figure 4a** illustrates the fit of the limited-information model to the data of a single session. We fitted the average information increase with each added neuron (**Fig. 4a**, top), and from this predicted the total information for each population size (**Fig. 4a**, bottom). Bayesian model comparison to a model that assumed unbounded information scaling confirmed that a model with limited asymptotic information was better able to explain the measured information scaling (Watanabe-Akaike Information Criterion WAIC_unlim_=-529.25 vs. WAIC_lim_=-531.59; smaller is better). This was the case for almost all discriminations with *δθ* = 45° across sessions and mice (**Fig. S1a**). Furthermore, the same procedure applied to the shuffled data resulted in better model fits for the unbounded information model, confirming that our model comparison was not *a priori* biased towards the limited-information model (**Fig. S1a**). Two sets of simulations with idealized and realistic neural models further confirmed that this model comparison was able to recover the correct underlying information scaling (**Fig. S2**). Therefore, information about drift direction is limited in the neural population responses within our dataset.

**Figure 4.**
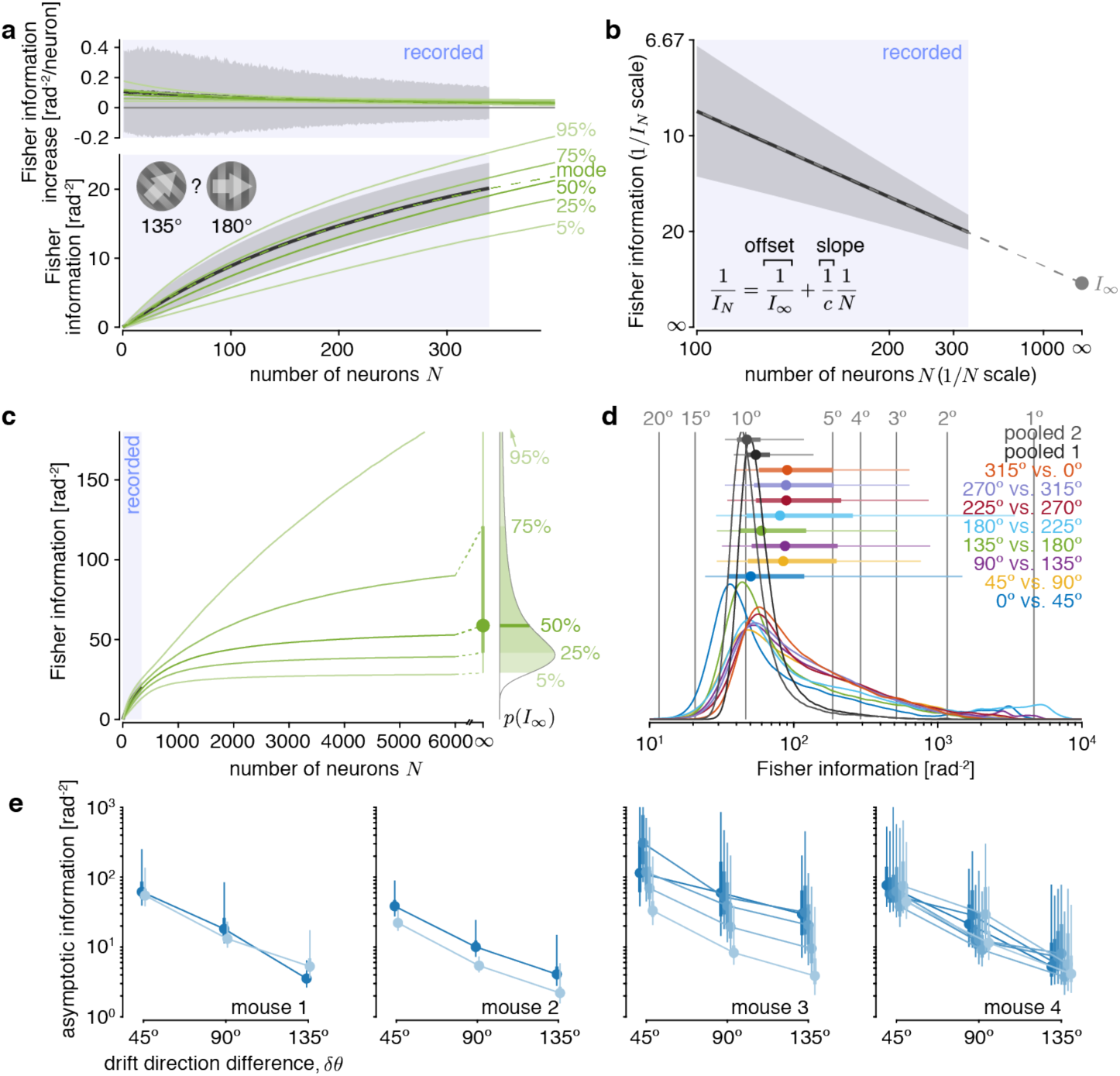
Information about drift direction is estimated to asymptote in large neural populations. **a.** Example information scaling fit, showing data (black; mean estimate ± 1SD; computed from 135**°** vs. 180**°** drift direction trials, as in **Fig. 3c**) and posterior predictive density for Bayesian fit (green; solid = percentiles, dashed = mode) for the Fisher information increase (top) and Fisher information (bottom) across different population sizes *N*. The model is fitted to the Fisher information increase estimates (top), as these are statistically independent across different population sizes. **b.** Plotting the inverse Fisher information 1/*I*_*N*_ over the inverse population size 1/*N* (mean estimate ± 1SD; same data as in **a**) shows an almost perfect linear scaling, as predicted by our theory. Fitting a linear model (grey dashed line) reveals a non-zero asymptotic information *I*∞ (grey dot) with *N* → ∞. The horizontal axis is reversed, such that population size grows towards the right, where 1/*N* becomes zero. **c.** The fitted model supports extrapolating the posterior predictive density beyond recorded population sizes (blue shaded area in **a, b**, and **c**) up to *N* → ∞. This results in a Bayesian posterior estimate over the asymptotic information *I*∞ (right), that we summarize by its median (dot), and its 50% (thick line) and 90% (thin line; truncated at top) credible intervals. **d.** Estimates of asymptotic information resulting from different drift direction pairs (colors; *δθ* = 45° for all pairs) results in comparable posterior densities (colored lines; associated density summaries above densities as in **c**) across different pairs. Therefore, we pooled the data across all non-overlapping pairs with the same *δθ* to achieve a more precise estimate. The pooled estimates were comparable across two different sets of non-overlapping pairs (grey). The vertical grey lines and numbers indicate the drift direction discrimination thresholds corresponding to different Fisher information estimates. **e.** The asymptotic Fisher information estimate (density summaries as in **c**; lines connect posterior medians) is comparable across sessions (different colors; horizontally shifted to ease comparison) and mice.

This result of limited drift direction information was corroborated by a second analysis. We start by observing that Eq. (1) can be rewritten as 1/*I*_*N*_ = *a*(1/*N*) + 1/*I*_∞_, which is linear in the inverse population size 1/*N* with slope *a* = 1/*c*. Increasing the population size, *N* → ∞, causes the inverse information to approach the asymptotic information, 1/*I*_*N*_ → 1/*I*_∞_. Therefore, we can distinguish between limited asymptotic information and unbounded information scaling (i.e., *I*_∞_ → ∞) by plotting 1/*I*_*N*_ against 1/*N*, and estimating its intercept at 1/*N* → 0. A non-zero intercept confirms limited asymptotic information, whereas a zero intercept would suggest information to scale without apparent bounds. When we analyzed the previous single-session data, we found that the inverse information indeed tightly scales linearly with the information population size (linear regression, adjusted *R*^2^ ≈ 1), as predicted by the model (**Fig. 4b**). Furthermore, the intercept at 1/*N* → 0 was significantly above zero (linear regression, *β*_0_ ≈ 0.023, two-sided *p* < 10^−6^), suggesting that information saturates with *N*. We found comparably good linear fits for all sessions/mice across all *δθ* = 45°discriminations (average adjusted *R*^2^ ≈ 0.999; **Fig. S3a**), and intercepts that were all significantly above zero (⟨*β*_0_⟩ ≈ 0.023, *t*_63_ = 17.95, two-sided *p* < 10^−10^ across non-overlapping discriminations; **Fig. S3b**), confirming the results of our model comparison.

In addition to supporting the distinction between information-limited and unbounded information scaling, TILC also allowed us to estimate the magnitude at which information would asymptote if we increased the population size beyond that of our recorded population. This is a theoretical measure that would be reached only for infinitely large virtual populations that have the same statistical structure as the recorded neurons. Despite this limitation, it gives insight into the order of magnitude of the information that we could expect to be encoded in the large populations of neurons present in mammalian cortices. To quantify the uncertainty associated with extrapolations beyond observed population sizes, we relied on Bayesian model fits that provide posterior distributions over our estimates of *I*_∞_, as illustrated in **Fig. 4c**. These posteriors were comparable across the discrimination of different drift direction pairs (**Fig. 4d**). Comparable information estimates across different drift direction pairs were essential to make these estimates meaningful, as different estimates would have implied that these estimates are driven by neural subsets within a heterogeneous population rather than being a statistical property of the whole population, as desired. Furthermore, it allowed us to reduce our uncertainty in the *I*_∞_ estimates by pooling the fits across different, non-overlapping drift direction pairs (**Fig. 4d**; grey). Indeed, Bayesian model comparison that accounts for the larger number of parameters of multiple individual per-discrimination fits confirmed that those were outperformed by pooled fits for all but two experimental sessions across all tested drift direction differences (**Fig. S4**). This provided further evidence that, for a fixed drift direction difference, the measured information scaling was statistically indistinguishable across different discriminations within each session.

Comparing these pooled estimates across sessions and mice revealed these estimates to be similar (**Fig. 4e**). These estimates dropped with an increase in the angular difference *δθ* in the compared drift directions, as is to be expected from a linear decoder used to discriminate between circular quantities (**Fig. S5**). Together, these observations strongly suggest that the recorded populations were part of a larger population that encoded limited information about the drift direction of the presented stimuli.

### No neural subpopulation encodes a disproportionate amount of information across all stimulus drift directions

The recorded population might contain neurons that are not only untuned to drift direction, but also do not contribute information through being correlated with other neurons in the population (Leavitt et al., 2017; Pruszynski & Zylberberg, 2019). As our information scaling measures are averaged across different orderings of how neurons are added to the population, uninformative neurons would contribute at different population sizes across different orderings. As a result, they make information scaling curves appear shallower than for populations that exclude uninformative neurons. These shallower scaling curves could in turn impact our estimates of asymptotic information (**Fig. 4**).

To ensure that uninformative neurons did not significantly affect our estimates, we asked if we could identify neural subpopulations within the set of recorded neurons that encode most of the information. Previous work identified such subpopulations in auditory cortex (Ince, Panzeri, & Kayser, 2013) and lateral prefrontal cortex (Leavitt et al., 2017) of monkeys, but we are not aware of any work that has shown this for V1. To identify highly informative subpopulations, we ordered the neurons within the recorded population by incrementally adding the neuron that resulted in the largest overall information increase (Ince et al., 2013; Leavitt et al., 2017). With this ordering, 90% of the information in the recorded population for a particular discrimination could be recovered from only about 30% of the recorded neurons (**Fig. 5a**). However, the same subpopulation did not perform well once we changed the pair of drift directions that were to be discriminated (**Fig. 5a**; left vs. right), for which the same subset of neurons only recovered about 55% of the information. Even a population ordering that boosted the average information across all drift direction pairs did not reveal a highly informative subpopulation within the recorded set of neurons (**Fig. 5a**; green). To determine whether there is any advantage to a particular ordering, we estimated the population size required to capture 90% of information of the recorded population if we ordered the neurons according to this objective. Across sessions/mice and discriminations, the required population size turns out to not differ significantly compared with a random ordering of the population (**Fig. 5b**; *t*_63_ = −0.215, two-sided *p* ≈ 0.83; across non-overlapping *δθ* = 45° discriminations). If a significant fraction of neurons is uninformative across all drift direction pairs, we would expect these sizes to differ. Therefore, it is unlikely that our asymptotic information estimates were significantly influenced by the presence of uninformative neurons in the recorded populations.

**Figure 5.**
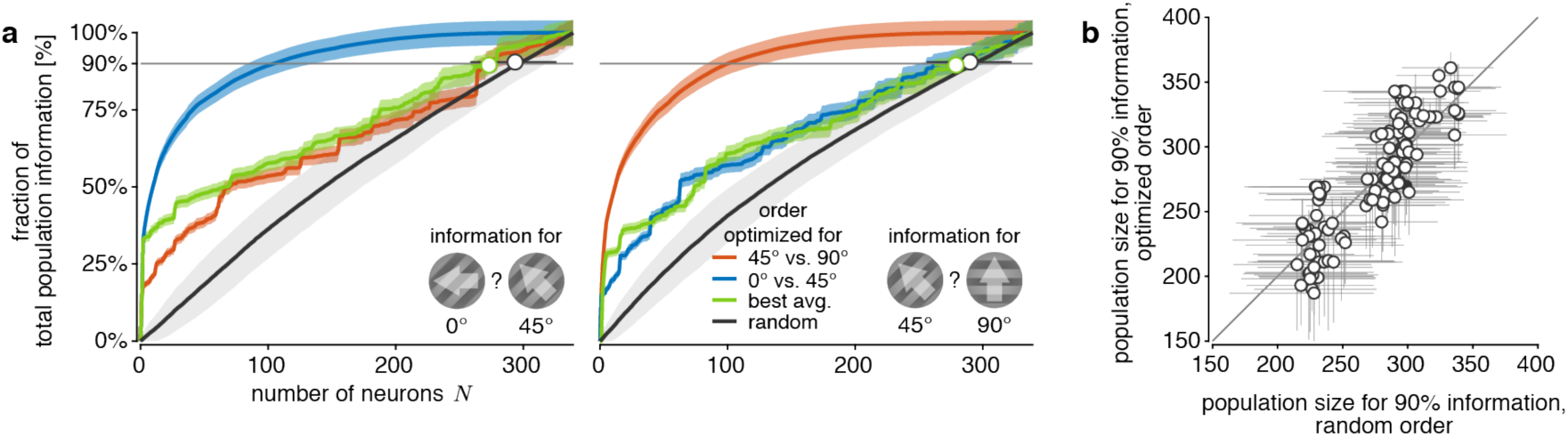
No single neural subpopulation appears to encode a disproportionate amount of information across all stimulus drift directions. **a.** Both panels show that the information increase in the recorded population depends on the order with which neurons are added to the population (colors). The panels differ in the considered drift direction discrimination (left: 0° vs. 45°; right: 45° vs 90°). The neuron order was optimized by incrementally adding the neuron that resulted in the largest information increase for a 0° vs. 45° (blue) or 45° vs 90° (orange) drift direction discrimination, or largest average increase across all discriminations with *δθ* = 45° (green). The optimal ordering for the 0° vs. 45° was also applied to the 45° vs 90° discrimination (blue line in right panel) and vice versa (orange line in left panel). The average information increase across random orders (black) is shown as baseline reference. Shaded error regions illustrate the uncertainty (mean +/- 1SD) due to limited numbers of trials (all curves), and variability across random orderings (black only). The black and green open circle (bootstrapped median +/- 95% CI) show the population sizes required to capture 90% of the information in the recorded population for the associated orderings. **b.** Plotting population sizes required to capture 90% of the information in the recorded population (bootstrapped median +/- 95% CI) for random ordering vs. orderings optimized to maximize average information across all discriminations revealed no significant difference between the two orderings. Each dot reflects one discrimination for one session.

### A finite-population information change impacts asymptotic information

If estimated asymptotic information mirrors the total information encoded by the animals’ brains, it should increase if we increase the amount of information provided by the stimulus in retinal photoreceptor activity. As has been shown previously, higher contrast stimuli result in higher decoding performance from recorded population responses (e.g., Busse et al., 2011). However, we might observe an information increase in recorded populations even when the asymptotic information remains unchanged (**Fig. 6c**, right). To determine if increasing the stimulus contrast results in an increase of asymptotic information, we performed a separate set of experiments in which two mice observed the same drift directions as before, but with a grating contrast that pseudo-randomly varied between 10% and 25% across trials. We hypothesized that the 25% contrast stimuli provide more information about the drift direction, and expected a corresponding increase in asymptotic information.

**Figure 6.**
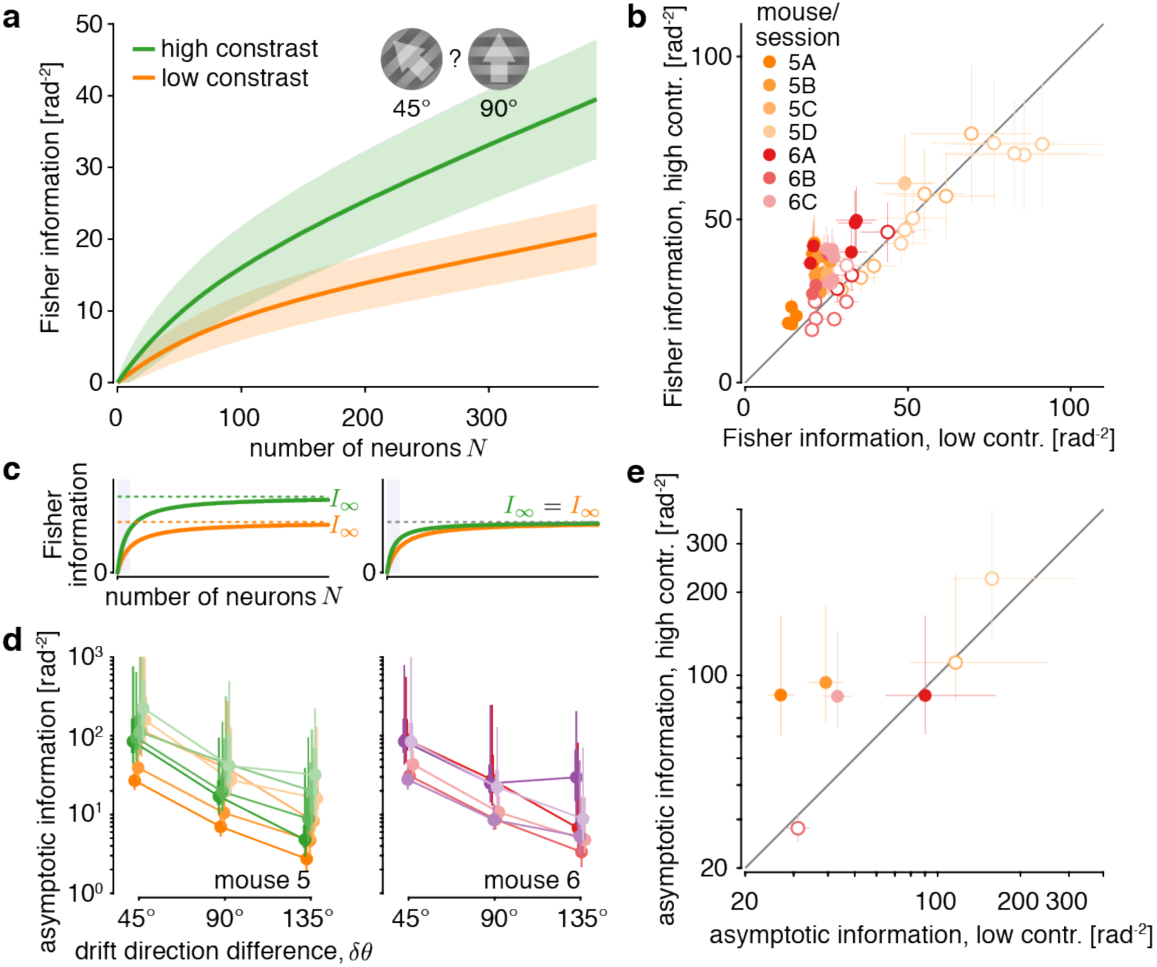
Increasing stimulus contrast boosts asymptotic information in V1. **a.** The information increases more rapidly with population size for high-contrast stimuli (green; mean +/- 1SD) than for low-contrast stimuli (orange; mean +/- 1SD), here shown for the discrimination between 45**°** vs. 90**°** drift direction trials of one session of mouse 5. An increase in stimulus contrast significantly increases the total information in the recorded population (bootstrap, two-sided *p* < 10^−5^). **b.** The information in the recorded population was larger for high than low stimulus contrast for most *δθ* = 45° discriminations across mice and sessions (colors, XY = mouse X, session Y). Each dot shows the information for one discrimination between different drift directions (8 dots per session; error bars = +/- 1SD of the information estimation uncertainty). Filled dots indicate a significant information increase (bootstrap, two-sided *p* ≥ 0.05). **c.** Observing an information increase in the recorded population (blue shaded area) does not necessarily imply an increase in asymptotic information (left vs. right). **d.** The estimated asymptotic information was generally higher for high-contrast stimuli (green/magenta; shades = sessions) than low-contrast stimuli (orange/red; shades = session; colors as in panel **b**), across different drift direction differences *δθ* (pooled estimates across different drift direction pairs; posterior density summaries as in **Fig. 4c**). **e.** To compare the pooled asymptotic information estimates for *δθ* = 45° for low-contrast trials to those for high-contrast trials, we plot them against each other (one dot per session, colors as in panel **b**, error bars = 50% posterior credible intervals). The four filled dots indicate sessions for which the information in the recorded population (panel **b**) is significantly larger for higher contrast stimuli for the majority of discriminations.

Information encoded in the recorded populations significantly increased for higher stimulus contrasts (**Fig. 6a** for single discrimination and session; **Fig. 6b** for all sessions/mice, non-overlapping discriminations with *δθ* = 45°: paired *t*_27_ = 2.78, two-sided *p* ≈ 0.0098). We in turn applied the same procedure as before (see **Fig. 4e**) to estimate asymptotic information, but did so separately for the two contrasts (**Fig. 6d**). We then compared these estimates for *δθ* = 45° within each session between low- and high-contrast trials (**Fig. 6d**). In principle, increasing contrast could increase asymptotic information, or it could leave asymptotic information unchanged (**Fig. 5c**). For three out of the four sessions in which information in the recorded population increased with contrasts for a majority of discriminations (as shown in **Fig. 6b**), we also observed an increase in asymptotic information with contrast (**Fig. 6e**, filled dots). This suggests that a more informative stimulus not only increased information in the recorded neural populations, but also in the larger (unrecorded) neural population.

### Tens of thousands of neurons are required to decode most of the information about stimulus drift direction

Information in the brain must saturate, as noisy sensors fundamentally limit the sensory information it receives. However, it remains unclear whether information saturates within the population size of V1 (**Fig. 1**). In our information scaling model, Eq. (1), saturation by definition only occurs in the limit of infinite neurons. We can nonetheless use the model to estimate saturating population sizes by asking how large these populations need to be to encode a large fraction of the asymptotic information (**Fig. 7a**). We will here focus on population sizes *N*_95_ that achieve 95% of asymptotic information, which can be found by setting *I*_*N*_ = 0.95 *I*_∞_ in Eq. (1) and solving for *N*. The required population sizes for other fractions of asymptotic information are easily found by a rescaling of *N*_95_ (**Fig. S6**).

**Figure 7.**
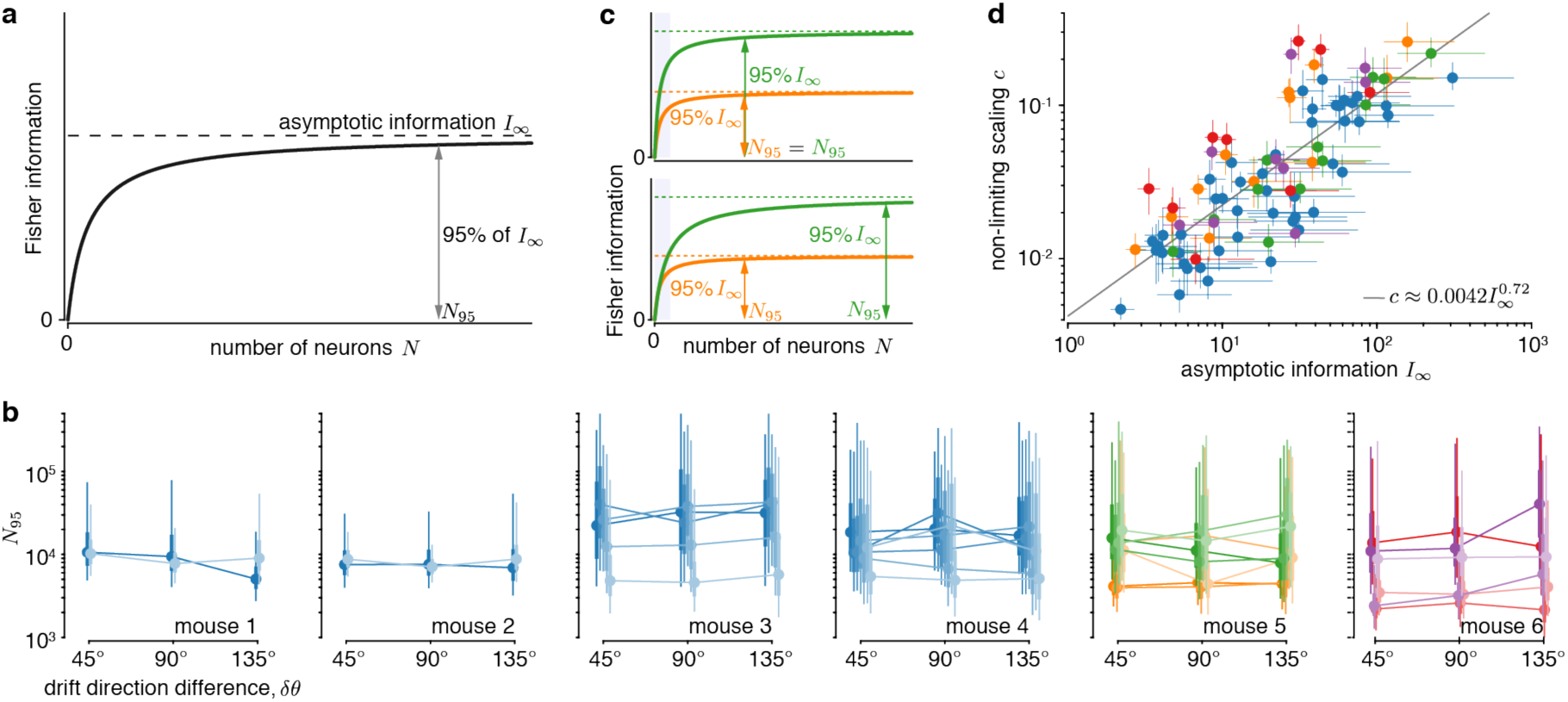
Tens of thousands of neurons are required to capture most of the information about stimulus drift direction. **a.** The TILC predicts how information grows with population size and allows us to estimate the population size *N*_95_ required to capture 95% of the asymptotic information *I*_∞_. **b.** Applied to our data, we find *N*_95_ in the order of tens of thousands of neurons, consistently across mice (panels) and sessions (colors; blue = uniform contrasts; orange/red = low contrast; green/magenta = high contrast; lines connect individual sessions; horizontally shifted to ease comparison; posterior densities as is **Fig. 4c**). **c.** An increase in information (orange to green) could be achieved by increasing the average information per neuron (top) or by leaving the average information per neuron roughly unchanged while recruiting more neurons (bottom). We would expect *N*_95_ to grow in the second, but not the first case. These two cases are hard to distinguish from the observed information scaling in smaller populations (shaded blue). **d.** Plotting estimated non-limiting scaling *c* over asymptotic information *I*_∞_ (dot = median, lines = 50% credible interval; colors as in panel **b**) for all animals, sessions, drift direction differences, and contrasts from **b** reveals that *c* grows sub-linearly with *I*_∞_ (grey line = linear regression of median estimates in log-log plot), which indicates that the estimated population size *N*_95_ increases weakly with the asymptotic information.

To estimate *N*_95_, we again relied on the information scaling fits pooled across non-overlapping pairs of drift directions. The recovered population sizes were all on the order of tens of thousands of neurons (**Fig. 7b**). Our previous analysis (**Fig. 5**) makes it unlikely that uninformative neurons within the recorded population strongly impact our estimated population sizes. Interestingly, increasing the drift direction difference *δθ* did not strongly affect these estimates (mice 1-4 in **Fig. 7b**), even though it modulated asymptotic information (**Fig. 4d**). Increasing stimulus contrast appeared to increase the estimated population sizes (mice 5-6 in **Fig. 7b**, orange vs. green), but not consistently so. Thus, it was unclear if a change in information resulted in a global re-scaling of the information scaling curve without changing its shape (**Fig. 7c**, top), or in the need for more neurons to encode this information (**Fig. 7c**, bottom).

To clarify the relationship between the asymptotic information *I*_∞_ and required population size *N*_95_, we did not directly relate these two quantities, as *N*_95_ is derived from the estimate of *I*_∞_. Instead, we relied on the property that *N*_95_ is proportional to *I*_∞_ /*c*, where *c* is the scaling factor associated with the non-limiting covariance component (see Eq. (1); Methods). Therefore, if *N*_95_ remains constant across different estimates of *I*_∞_ and *c*, these two quantities need to vary in proportion to each other. In a log-log plot, this implies that the slope describing their relationship would be one. However, we found a slope of *β*_1_ ≈ 0.72, which is slightly, but significantly below one (**Fig. 7d**; F-test, *F*_1_ = 21.49, *p* ≈ 1.2 × 10^−5^). Substituting the measured relationship between *c* and *I*_∞_ into the expression for *N*_95_ results in *N*_95_ ≈ 4523.8 *I*_∞_ ^0.28^. This implies that the population size required to encode 95% of the asymptotic information increases with *I*_∞_, but does so only weakly. To illustrate this weak increase, let us consider sessions in which the estimated asymptotic information increased three-fold with an increase in stimulus contrast (**Fig. 6e**). In this case, a population of the size required to capture 95% of the asymptotic information for low-contrast trials could capture 93% of the asymptotic information for high-contrast trials (see Methods).

### Information is not well-aligned with principal noise dimensions

Previous work has observed that most neural population activity fluctuations are constrained to a low-dimensional linear subspace that is embedded in the high-dimensional space of neural activity (Engel & Steinmetz, 2019; Semedo, Zandvakili, Machens, Yu, & Kohn, 2019; Williamson et al., 2016). This might suggest that focusing on such a low-dimensional subspace is sufficient to understand brain function (Williamson et al., 2016). Thus, we asked if we can recover most of the information about visual drift direction from such subspaces, defined by the dimensions where population activity is most variable. The information encoded in each dimension grows with how well the signal, ***f***′, is aligned with this dimension, but shrinks with the magnitude of noise in this dimension (**Fig. 8a**; (Mendels & Shamir, 2018; Moreno-Bote et al., 2014)). This tradeoff makes it unclear whether the subspace where population activity is the most variable is indeed the subspace that encodes the most information.

**Figure 8.**
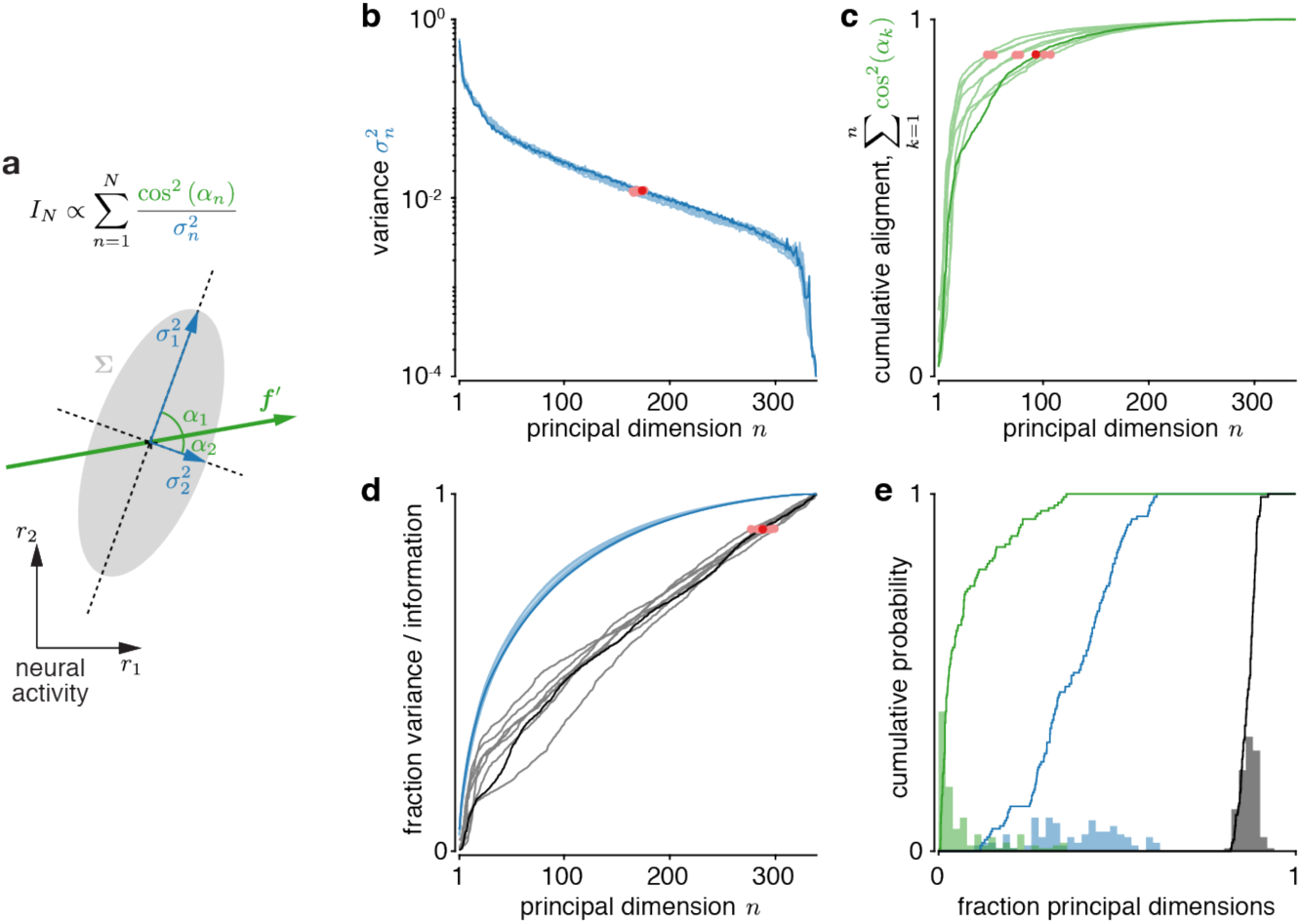
Information is not well-aligned with principal noise dimensions. **a.** The total information in a recorded population, *I*_*N*_, can be decomposed into how well the change in population tuning, ***f***′, is aligned to the different principal dimensions of the noise covariance (given by cos^2^(*α*_*n*_)), as well as the noise variances, 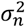, in these dimensions. Low-variance dimensions that are strongly aligned to ***f***′ contribute more information. **b.** The variance along principal dimensions drops rapidly with dimension (red dots: >90% of total variance). **c.** The cumulative alignment of ***f***′ to the principal dimensions of the noise covariance rises rapidly with principal dimension (red dots: >90% of total alignment), indicating that ***f***′ is most strongly aligned to the first few principal dimensions. **d.** The fraction of total information (black; red dots: >90% information) rises more slowly with additional principal dimensions than the fraction of total noise variance (blue). The principal dimensions in panels **b**-**d** are ordered in decreasing order of variance, and show data for the same session as in **Fig. 3c** (dark = same discrimination as in **Fig. 3c**; light = other *δθ* = 45° discriminations for that session). **e.** The histograms (bar plots) and cumulative probabilities (lines) of the fractions of the total number of principal dimensions at which the cumulative ***f***′ alignment (green), cumulative variance (blue), and cumulative information (black) exceed 90% of their respective totals. These fractions are shown for all *δθ* = 45° discriminations, sessions and mice. All estimates are cross-validated, and averaged across ten train/test splits (see Methods).

We found the principal dimensions of the noise covariance matrix and asked how much information a subset of the most variable dimensions is able to encode. In our data, 90% of the total variance was captured by approximately 37.6% ± 12.4pp (mean % ± 1SD percentage points across all sessions/mice, *δθ* = 45°discriminations) of all available dimensions (**Fig. 8b/e**), confirming previous reports that relatively few dimensions are required to capture most noise variance. Furthermore, ***f***′ was most strongly aligned to the first few of these principal dimensions (Mendels & Shamir, 2018) (**Fig. 8c**). Using cosine similarity to measure this alignment, we found that 90% of the cumulative alignment was reached by approximately 7.4% ± 9.1pp of all available dimensions (**Fig. 8c/e**). Finally, we asked how many dimensions were required to capture 90% of the information encoded in the recorded population. Even though later dimensions were not well-aligned with ***f***′ (see the shallow cumulative alignment increase in **Fig. 8c**), they were also less noisy (**Fig. 8b**) and so could contribute significantly to the encoded information. As evident by the continual information growth in **Fig. 8d**, this resulted in information which was fairly evenly spread across all dimensions, such that, on average, approximately 86.7% ± 2.2pp of all principal noise dimensions were required to encode 90% of all of the recorded information. This is significantly higher than the fraction required to capture 90% of all variance (difference = 48.7 ± 1.5pp, mean ± 1SEM, paired *t*_63_ = 32.53, two-sided *p* < 10^−6^ across non-overlapping discriminations). In fact, if we restricted ourselves to the subspace that captures 90% of all noise variance, we could only decode 58.9% ± 5.6pp of information. Therefore, in our data, relying only on information encoded in the subspace of most variable principal dimensions would result in significant information loss.

## Discussion

We asked how information about the drift direction of a visual stimulus is distributed in large neural populations, and addressed this question by analyzing how information scales with population size. We observed that, in recorded populations, information scaled sublinearly with population size, indicating that noise correlations limited this information. The information scaled in line with TILC if information is indeed limited in larger populations. Based on this theory, we found that we require on the order of tens of thousands of neurons to encode 95% of the asymptotic information. When varying input information by changing stimulus contrast, the required population size appeared to change. Indeed, we found that more information required larger populations, but this relationship was extremely weak. Overall, these findings suggest the presence of information-limiting correlations that cause information in mouse V1 to saturate with population size, indicating the use of a highly redundant, distributed neural code within mouse V1.

Previous attempts at measuring how information scales with population size have frequently found noise correlations to either be beneficial (Denman & Reid, 2019) or to not affect information scaling (Cotton et al., 2018; Mendels & Shamir, 2018). These studies focused on smaller populations (<200 neurons in Denman & Reid, 2019; <100 neurons in Mendels & Shamir, 2018) in which sublinear scaling might be hard to identify (**Fig. 1**), and in part included spike timing information (Denman & Reid, 2019) in addition to the spike counts used here. Recent recordings from ∼20,000 neurons in mouse V1 suggest information about visual stimuli does saturate (Stringer, Michaelos, & Pachitariu, 2019; Fig. S6), but it appears to do so above the population sizes we estimated. These recordings used a slower image scan rate (3 Hz vs. the 30 Hz used for this study), which introduces additional recording noise. This additional noise makes information saturate more slowly with population size (see SI), potentially explaining the larger required population sizes.

Even though information is highly distributed across neurons in a population, most variability is captured by a low-dimensional subspace, leading to suggestions that we might only need to consider the information encoded in this subspace (Williamson et al., 2016). As we have shown, this argument does not consider that information does not only depend on variability, but also on how the signal aligns with this variability (**Fig. 8a**). Once both are taken into account, the dimensions of largest variability become a poor proxy for the most informative dimensions (**Fig. 8d**). This is in line with recent work showing that the most variable subspace in macaque V1 is different from the one that most co-varies between V1 and V2 (Semedo et al., 2019), which presumably transmits information between these areas. Our work explicitly shows such misalignment, and does so in larger populations.

To compare our required population size estimates to the total number of neurons in mouse V1, we conservatively estimated the need for about 48,000 neurons (see Methods) to achieve drift direction discrimination performance that most likely exceeds that of the animals (Abdolrahmani, Lyamzin, Aoki, & Benucci, 2019; Glickfeld, Histed, & Maunsell, 2013). Our use of time-deconvolved calcium activity as a noisy proxy for spike counts (T.-W. Chen et al., 2013; Ledochowitsch et al., 2019) makes these estimates upper bounds on required population sizes (see SI). Nonetheless, they compare favorably to the number of neurons in mouse V1, whose estimates range from 283,000 to 655,500 (Herculano-Houzel, Watson, & Paxinos, 2013; Keller, Erö, & Markram, 2018). This confirms that mouse V1 has more neurons than required to encode most of the estimated asymptotic information about the direction of a moving visual stimulus.

If animals are required to perform tasks that rely on the encoded information we measured (e.g., to discriminate between different drift directions), each neuron in the population would ideally contribute to the animal’s choices. Quantified by choice correlations (Britten, Newsome, Shadlen, Celebrini, & Movshon, 1996; Haefner, Gerwinn, Macke, & Bethge, 2013), an optimal read-out requires the choice correlations of individual neurons to be the fraction of the population’s discrimination threshold over that of the neuron (Pitkow, Liu, Angelaki, DeAngelis, & Pouget, 2015). In contrast to previous work (e.g., Britten, Shadlen, Newsome, & Movshon, 1992) that found that individual neurons’ thresholds match that of the animal, the neurons’ average threshold in our data (see information for *N* = 1 in **Fig. 3c**) is exceedingly small when compared to that of the recorded population (**Fig. 3c** for full population), and even smaller when compared to estimated asymptotic information (**Fig. 4e**). This mismatch might arise from shorter stimulus presentation, not tailoring the stimuli to match the neuron’s tuning (as done in Britten et al. (1992)), recording from lower-level visual areas (V1 vs. V4 or MT) with smaller receptive fields, as well as increased recording noise with calcium imaging as compared to electrophysiological recordings. These lower discrimination thresholds predict increasingly small choice correlations, in line with recent reports from area V1 of monkeys, where fewer than 7% of V1 neurons were found to feature significant choice correlations (Jasper, Tanabe, & Kohn, 2019). In general, the estimated asymptotic information predicted direction discrimination thresholds compatible with previous behavioral reports in mice (Abdolrahmani et al., 2019; Glickfeld et al., 2013), but the use of different stimuli in these experiments precludes a direct quantitative comparison. A more detailed analysis of the relation between neural activity and choice would require training animals to report their percepts, and then relating these reports to population activity fluctuations.

A prediction of our findings is that neural information should continue to scale according to Eq. (1) in larger populations than those recorded in our experiments. Testing these predictions involves precise estimates of noise correlations, which require about the same number of trials in which the same stimulus (e.g., drift direction) is presented as there are neurons in the population (Kanitscheider, Coen-Cagli, & Pouget, 2015; Moreno-Bote et al., 2014). Therefore, even with more powerful recording techniques, information estimates might be limited by the number of trials that can be collected within individual sessions. The use of decoders to estimate information might sidestep these estimates (Kanitscheider, Coen-Cagli, Kohn, et al., 2015; Whiteway, Bartolo, Averbeck, & Butts, 2017), with the downside of potentially confounding decoder biases. A further challenge is to record from a population that homogeneously encodes the same amount of information about each stimulus. Such homogeneity ensures that the estimated asymptotic information and population sizes are not specific to particular stimulus values. The weak spatial organization of drift direction selectivity in mouse V1 (Ringach et al., 2016) supports this, but the same would be harder to achieve in monkeys due to the much stronger spatial correlations of orientation and direction selectivity in their visual cortices (Dow, 2002). Finally, even if Eq. (1) is confirmed to match the information in larger populations than used here, it does not allow us to guarantee that the cortex’s information is limited by sensory noise and suboptimal computations. Though unlikely, information might continue to grow linearly after an initial sublinear growth (Ecker et al., 2011). The only way to conclusively rule out this scenario is to record from all neurons in the information-encoding population, which, at least in mammals, will likely not be possible in the foreseeable future (Mott, Gordon, & Koroshetz, 2018).

Although all information entering the brain is limited by sensory noise (Faisal et al., 2008), such that it can never grow without bound, the information could be so plentiful as to not saturate within the population sizes of mammalian sensory areas. Our findings suggest this not to be the case. However, we suspect the main limiting factor not to be noisy sensors. Instead, most problems that the brain has to deal with require fundamentally intractable computations that need to be approximated, resulting in substantial information loss (Beck, Ma, Pitkow, Latham, & Pouget, 2012). Indeed, suboptimal computations can dominate overall information loss, and resulting behavioral variability (Acerbi, Vijayakumar, & Wolpert, 2014; Drugowitsch, Wyart, Devauchelle, & Koechlin, 2016), such that they might be the main contributor to the information limitations we observe in our experiments.

If the brain operates in a regime in which information in sensory areas is limited, all information the brain deals with is uncertain. This idea finds support in the large body of work showing that behavior is well-described by Bayesian decision theory (Doya, Ishii, Pouget, & Rao, 2006; Moreno-Bote, Knill, & Pouget, 2011; Pouget, Beck, Ma, & Latham, 2013), which makes effective use of uncertainty. This, in turn, implies that the brain encodes this uncertainty, but its exact neural representations remain unclear (Fiser, Berkes, Orbán, & Lengyel, 2010; Pouget et al., 2013). A further consequence of limited information is that theories that operate on trial averages (e.g., Gao & Ganguli, 2015; Gao et al., 2017; Kobak et al., 2016) or assume essentially unlimited information (e.g., Ecker et al., 2011) only provide an incomplete picture of the brain’s operation. Therefore, an important next step is to refine these theories to account for trial-by-trial variation in the encoded information to achieve a more complete picture of how the brain processes information in individual trials, rather than on average.

## Online Methods

All experimental procedures were conducted according to the IACUC.

### Animals and surgery

Male C57BL/6J mice were obtained from The Jackson Laboratory and used for imaging experiments between 4-7 months of age. Prior to imaging, mice underwent surgery to implant a chronic cranial window and headplate. Mice were injected intraperitoneally with dexamethasone (3 μg per g body weight) 3-6 hrs before surgery to reduce brain swelling. During surgery, mice were stably anesthetized with isoflurane (1-2% in air). A titanium headplate was attached to the skull using dental cement (C&B Metabond, Parkell). A ∼3.5-mm diameter craniotomy was made over left V1 (stereotaxic coordinates: 2.5 mm lateral, 3.4 mm posterior to bregma). AAV2/1-syn-GCaMP6s (Penn Vector Core) was diluted into phosphate-buffered saline at a final titer of ∼2.5E12 gc/ml and mixed 10:1 with 0.5% Fast Green FCF dye (Sigma-Aldrich) for visualization. Virus was injected in a 3×3 grid with 350 μm spacing near the center of the craniotomy at 250 μm below the dura, with ∼75 nl at each site. Injections were made slowly (over 2-5 min) and continuously using beveled glass pipettes and a custom air pressure injection system. The pipette was left in place for an additional 2-5 min after each injection. Following injections, the dura was removed. A glass plug consisting of two 3.5-mm coverslips and one 4.5-mm coverslip (#1 thickness, Warner Instruments) glued together with UV-curable transparent optical adhesive (Norland Optics, NOA 65) was inserted into the craniotomy and cemented in place with cyanoacrylate (Insta-Cure, Bob Smith Industries) and metabond mixed with carbon powder (Sigma-Aldrich) to prevent light contamination from the visual stimulus. An aluminum ring was then cemented on top of the headplate, which interfaced with the objective lens of the microscope through black rubber light shielding to provide additional light-proofing. Data from mouse 1 and 2 were collected as part of a previously published study (Chettih & Harvey, 2019), following a similar surgical protocol. Imaging datasets were collected at least 2 weeks post-surgery, and data collection was discontinued once baseline GCaMP levels and expression in nuclei appeared to be high.

### Visual stimuli

Visual stimuli were displayed on a gamma-corrected 27-inch IPS LCD gaming monitor (ASUS MG279Q). The monitor was positioned at an angle of 30° relative to the animal and such that the closest point to the mouse’s right eye was ∼24 cm away, with visual field coverage ∼103° in width and ∼71° in height. Visual stimuli were generated using PsychoPy (Peirce, 2007) or Psychtoolbox (for mice 1 and 2 only) and consisted of square-wave gratings presented on a grey background to match average luminance across stimuli. Gratings were gradually windowed with a Gaussian central aperture mask to prevent monitor edge artifacts. Grating drift directions were pseudo-randomly sampled from 45-360° in 45° increments at 10% or 25% contrast, spatial frequency of 0.035 cycles per degree, and temporal frequency of 2 Hz. Stimuli were presented for 500 ms, followed by a 500 ms grey stimulus during the inter-stimulus interval (1 Hz presentation). Digital triggers from the computer controlling visual stimuli were recorded simultaneously with the output of the ScanImage frame clock for offline alignment.

### Microscope design

Data were collected using a custom-built two-photon microscope. A Ti:Sapphire laser (Coherent Chameleon Vision II) was used to deliver 950 nm excitation light for calcium imaging through a Nikon 16 × 0.8 NA water immersion objective, with an average power of ∼60-70 mW at the sample. The scan head consisted of a resonant-galvonometric scanning mirror pair separated by a scan lens-based relay. Collection optics were housed in a light-tight aluminum box to prevent contamination from visual stimuli. Emitted light was filtered (525/50, Semrock) and collected by a GaAsP photomultiplier tube (Hamamatsu). Microscope hardware was controlled by ScanImage 2018 (Vidrio Technologies). Rotation of the spherical treadmill along three axes was monitored by a pair of optical sensors (ADNS-9800) embedded into the treadmill support communicating with a microcontroller (Teensy, 3.1). The treadmill was mounted on an XYZ translation stage (Dover Motion) to position the mouse under the objective.

### Experimental protocol

Before data acquisition, mice were habituated to handling, head-fixation on a spherical treadmill (Harvey, Coen, & Tank, 2012), and visual stimuli for 2-4 days. For each experiment, a field of view (FOV) was selected. Multiple experiments conducted in each animal were performed at different locations within V1 or different depths within layer 2/3 (120-180 μm below the brain surface). Before each experiment, the monitor position was adjusted such that a movable flashing stimulus or drifting grating in the center of the screen drove the strongest responses in the imaged FOV, as determined by online observation of neural activity. A single experiment consisted of three blocks of ∼45 min each. Once a FOV was chosen, a baseline image (∼680 × 680 μm) was stored and used throughout the entire experiment to compare with a live image of the current FOV and manually correct for axial and lateral drift (typically <3 μm between blocks and <10 μm over the full experiment) by adjusting the stage. Drift and image quality stability were verified post-hoc by examining 1,000x sped-up movies of the entire experiment after motion correction and temporal downsampling, and experiments that were unstable were discarded without further analysis. Data from mouse 1 and 2 were from previously published experiments (Chettih & Harvey, 2019), where a small fraction of neurons were photostimulated simultaneous to drifting gratings presentation. All photostimulated neurons were excluded from analysis for this paper.

### Data processing

Imaging data were processed as previously described (Chettih & Harvey, 2019). Briefly, imaging frames were first motion-corrected using custom MATLAB code (https://github.com/HarveyLab/Acquisition2P_class) on sub-frame, full-frame, and long (minutes to hours) timescales. Batches of 1,000 frames were corrected for rigid translation using subpixel image registration, after which frames were corrected for non-rigid warping on sub-frame timescales using a Lucas-Kanade method. Non-rigid deformation on long timescales was corrected by selecting a global alignment reference image (average of a 1,000-frame batch) and aligning other batches by fitting a rigid 2D translation, followed by an affine transform and then nonlinear warping. After motion correction, due to large dataset size (∼130 GB), imaging frames were temporally downsampled by a factor of 25 from 30 Hz to 1.2 Hz. Downsampled data was used to find spatial footprints, using a CNMF-based method (https://github.com/Selmaan/NMF-Source-Extraction). Full temporal-resolution fluorescence traces for each source (including cell bodies, processes, and unclassified sources) were then extracted and deconvolved using the constrained AR-1 OASIS method. Using a convolutional neural network trained on manually annotated labels, sources were identified as cell bodies, axial processes (bright spots), horizontal processes, or unclassified. Only data from cell bodies were used in this paper.

### Tuning curve fits

We used three nested models to fit tuning curves for each neuron. In the direction-tuned model, the average neural response of each neuron was fitted by a mixture of two Von Mises function given by

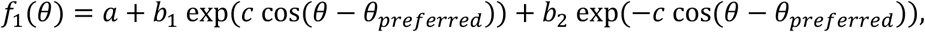

where *a, b*_1_, *b*_2_, *c*, and *θ*_*preferred*_ are model parameters, and *θ* is the stimulus’ drift direction. In the orientation-tuned model, the average neural response of each neuron was fitted using a single Von Mises function given by

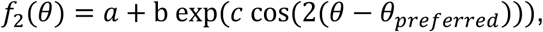

with parameters *a, b, c*, and *θ*_*preferred*_. The third and last model is a null model that assumes neurons are not significantly tuned to drift direction, and fits a constant value to neural responses, that is *f*_3_(*θ*) = *a*. We fitted all three models to the response of neuron across all trials by minimizing the sum of squared residuals between observed neural response and the tuning function across different stimulus drift direction. We then compared the nested models by an F-test (with Bonferroni correction for multiple comparisons) to test whether neurons are direction-tuned, orientation-tuned or untuned.

### Generalized Fisher information

Linear Fisher information (Ganguli & Simoncelli, 2014; Moreno-Bote et al., 2014; Seriès, Latham, & Pouget, 2004), which is the Fisher information that can be recovered by a linear decoder, can for stimulus *θ*_0_ be computed by *I*(*θ*_0_) = ***f***′(*θ*_0_)^*T*^Σ^−1^(*θ*_0_)***f***′(*θ*_0_). Here, ***f***′(*θ*_0_) is the vector of derivatives of each neuron’s average response with respect to *θ*, with the *i*th element given by *∂f*_*i*_(*θ*_0_)/*∂θ* = *∂* < *r*_*i*_|*θ*_0_ >/*∂θ*, and Σ(*θ*_0_) = cov(***r***|*θ*_0_) is the noise covariance of the population activity vector ***r***. Therefore, linear Fisher information is fully determined by the first two moments of the population activity, irrespective of the presence of higher-order moments. Furthermore, if 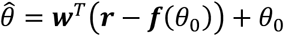 is the unbiased minimum-variance locally linear estimate of *θ*, its variance is given by 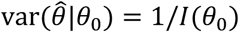 (Cover & Thomas, 2006). In practice, ***f***′(*θ*_0_) and Σ(*θ*_0_) are approximated by their empirical estimates, 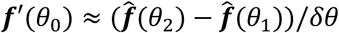, and Σ(*θ*_0_) ≈ (cov(***r***|*θ*_1_) + cov(***r***|*θ*_2_))/2, where *θ*_1,2_ = *θ*_0_ ∓ *δθ*/2. This naïve estimate is biased but a bias-corrected estimate can be used (Kanitscheider, Coen-Cagli, Kohn, et al., 2015).

By definition, Fisher information is a measure of fine discrimination performance around a specific reference *θ*_0_, requiring small *δθ*. As we show in the SI, the same measure with ***f***′(*θ*_0_) and Σ(*θ*_0_) replaced by their empirical estimate, can be used for coarse discrimination for which *δθ* is larger. Furthermore, this generalization corresponds to (*d*′/*δθ*)^2^, where *d*′ is the sensitivity index used in signal detection theory (Green & Swets, 1966), becomes equivalent to Fisher information in the *δθ* → 0 limit, and shares many properties with the original Fisher information estimate. In particular, the same bias correction leads to unbiased estimates. Kanitscheider et al. (2015) lack an estimate of the variance of the bias-corrected Fisher information estimate that can be computed from data, so we provide a derivation thereof in the SI.

To relate (generalized) Fisher information to discrimination thresholds, we observe that the variance of the stimulus estimate 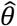 is 1/*I*(*θ*_0_). Assuming this estimate to be Gaussian across trials, the difference in estimates across two stimuli which differ by Δ*θ* is distributed as *N*(Δ*θ*, 2/*I*(*θ*_0_)). Therefore, the probability of correctly discriminating these stimuli is 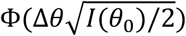 (Averbeck & Lee, 2006; Y. Chen, Geisler, & Seidemann, 2006; Nogueira et al., 2019), where Φ(·) is the cumulative function of a standard Gaussian. Setting the desired probability correct to 80% and solving for Δ*θ* results in the drift direction discrimination threshold 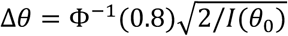.

### Estimating Fisher information from neural data

Our Fisher information estimates have two sources of uncertainty. First, they rely on empirical estimates of ***f***′(*θ*_0_) and Σ(*θ*_0_) from a limited number of trials that are thus noisy. Second, we assume that recorded neurons to be a small, random subsample of the full population. As we want to estimate the average Fisher information across such subsamples across different population sizes, observing only a single subsample introduces additional uncertainty.

We will first focus on the uncertainty due to a limited number of trials. We can find an unbiased estimate of *I*_*N*_ for a population of *N* neurons by a biased-corrected estimate *Î*_*N*_. Our aim is to fit models to how *Î*_*N*_ changes with *N*. We can estimate this change by computing *Î*_1_ for a single neuron, and then successively add neurons to the population to find *Î*_2_, *Î*_3_, … However, this procedure causes *Î*_*N*_ and *Î*_*N*+1_ to be correlated, as their estimates share the data of the previous *N* neurons. Therefore, we need to account for these correlations when fitting the information estimates across multiple *N*. Fortunately, the change in information across successive *N*, Δ*Î*_*N*_ = *Î*_*N*_ − *Î*_*N*−1_ is uncorrelated, that is cov(Δ*Î*_*N*_, Δ*Î*_*N*−1_) = 0 (see SI). The intuition underlying this independence is that the response of each neuron can be decomposed into a component that is collinear to the remaining population and one that is independent of it. Only the independent component contributes additional information, making the information increase due to adding this neuron independent of the information encoded in the remaining population. Overall, rather than fitting the information estimates, we will instead fit the information increases across different *N*.

To handle the uncertainty associated with subsampling larger populations, we assumed that the small recorded population is statistically representative of the full population. Then, our aim is to simulate random draws of the size of the recorded population from the full, much larger population. We achieved this simulation by randomly drawing neurons from the recorded population, without replacement, up to the full recorded population size, effectively resulting in a random order of adding recorded neurons to the population. For each such ordering, we estimated the information increase with each additional neuron. As the information in the total recorded population is the same, irrespective of this ordering, the information increases Δ*I*_*N*_ and Δ*I*_*M*_ for *N* ≠ *M* will on average be negatively correlated across different orderings. This is an artifact of re-using the same data to simulate samples from a larger population. As long as the full population is significantly larger than the one we recorded from, the probability of re-sampling the same pair of neurons from the full population is exceedingly small, such that we can ignore these correlations (see SI). Any negative correlations between information increases, however small, will reduce the variance of our Fisher information estimates. Therefore, by ignoring these correlations, we will estimate an upper bound of this variance, and thus overestimate the uncertainty. In summary, we estimated the uncertainty associated with subsampling larger populations by estimating the moments of the Fisher information increase by bootstrap estimates across different orderings with which neurons are added to the population. As shown in **Fig. S7a**, this procedure also captures the uncertainty associated with a limited number of trials, such that no extra correction is needed to account for this second source of uncertainty.

Overall, we estimated the moments of the Fisher information increase Δ*Î*_*N*_ for the discrimination of *θ*_1_ and *θ*_2_ as follows. First, we estimated the empirical moments 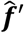 and 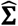 using the same number of trials for *θ*_1_ and *θ*_2_. Second, we chose a particular random order with which to add neurons to the population. Third, we used this order to estimate Δ*Î*_1_, Δ*Î*_2_, … by use of the biased-corrected Fisher information estimate applied to 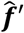 and 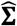. Fourth, we repeated this estimate across 10^4^ different neural ordering to get 10^4^ bootstrap estimates of the Fisher information increase sequence. Fifth, we used the bootstrap estimate to compute the moments µ_*N*_ = ⟨Δ*Î*_*N*_ ⟩ and 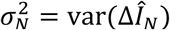 for each *N*, which we in turn use to fit the information scaling curves (see below). As the individual increases are independent across *N*, we used its moments to additionally estimate the moments of 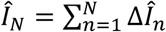, which are given by 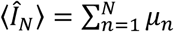 and 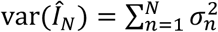. We used these moments to plot the Fisher information estimates in **Figs. 3a, 4b/d** and **5a**.

### Fisher information scaling with limited information

Moreno-Bote et al. (2014) have shown that for large populations encoding limited asymptotic information *I*_∞_, the noise covariance can be decomposed into 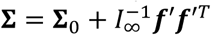, where only the ***f***′***f***′^*T*^ component, called *differential correlations*, limits information. Assuming a population size of *N* neurons, we can apply the Sherman-Morrison formula to the above noise covariance decomposition (Moreno-Bote et al., 2014; Pitkow et al., 2015) to find *I*_*N*_^−1^ = *I*_0,*N*_^−1^ + *I*_∞_^−1^, where 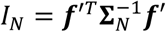 is the Fisher information in this population, and 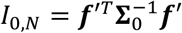 is the Fisher information associated with the non-limiting noise covariance component Σ_0_. Furthermore, assuming that this non-limiting component contributes average information *c* per neuron, that is *I*_0,*N*_ = *cN*, results in Eq. (1) in the main text. We also tested a model in which *I*_0,*N*_ initially scaled supralinearly in *N*. We found this model by integrating *c*(1 – *e*^−*N*/*τ*^) from zero to *N*, resulting in *I*_0,*N*_ = *c*(*N* + *τ*(*e*^−*N*/*τ*^ − 1)) with parameter *τ* that controls the extent of the initial supralinearity. The two models become equivalent with *τ* → 0. The above derivation relies on the traditional Fisher information definition for fine discrimination. The results remain unchanged when moving to Fisher information generalized to coarse discrimination.

### Fitting information scaling models

We compared three models for how Fisher information *I*_*N*_ scales with population size *N*. The first *unlim* model assumes linear scaling, *I*_*N*_ = *cN*, and has one parameter, *ϕ*_1_ = {*c*}. The second *lim* model, given by Eq. (1) in the main text, assumes asymptotic information *I*_∞_, and that the Fisher information associated with the non-limiting covariance component increased linearly, *I*_0,*N*_ = *cN*. This model thus has two parameters, *ϕ*_2_ = {*c, I*_∞_}. The third *lim-exp* model assumes an initial supralinear scaling of *I*_0,*N*_, as described above, and has three parameters, *ϕ*_3_ = {*c, I*_∞_, *τ*}. The *lim-exp* model fits the data consistently worse than the *lim* model (**Fig. S1b**), such we did not consider it in the main text.

As the Fisher information estimates in data are correlated across different population sizes, we did not directly fit these estimates. Instead, we fitted how they changed when adding additional neurons, as the estimated Fisher information increase is uncorrelated across different population sizes. That is, we used the likelihood function 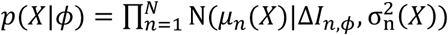, where *X* is the recorded data (that is, the recorded population activity in all trials with the drift directions that are being discriminated, yielding the desired moments *μ*_1_, …, *μ*_*N*_ and 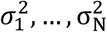), *ϕ* are the model parameters, Δ*I*_*n,ϕ*_ = *I*_*n,ϕ*_ − *I*_*n*−1,*ϕ*_ is the information increase predicted by that model, and *μ*_*n*_ and 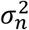 are the mean and variance of the estimated information increase in data *X* for a particular discrimination when moving from population size *n* − 1 to *n* (see further above).

We regularized the fits by weakly informative parameter priors. For *c* we used *p*(*c*) ∝ St_1_(⟨*μ*_*n*_⟩, 100(⟨*μ*_*n*_⟩ + 0.5)^2^), which is a Student’s t distribution with mean ⟨*μ*_*n*_⟩, variance 100(⟨*μ*_*n*_⟩ + 0.5)^2^ and one degree of freedom, and where ⟨*μ*_*n*_⟩ is the average estimated information increase in the recorded population. Thus, the prior is centered on the empirical estimate for *c* for the linear scaling model, but has a wide variance around this estimate. We furthermore limited *c* to the range *c* ∈ [0, ∞]. For *I*_∞_ we used *p*(*I*_∞_) ∝ St_1_(⟨*Î*_*N*_⟩, 100 *max*{1, ⟨*Î* _*N*_⟩}^2^) over *I*_∞_ ∈ [0, ∞], which is a weak prior centered on the empirical information estimate 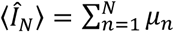 for the recorded population. For *τ* we used *p*(*τ*) ∝ St_1_(0, *N*^2^) over *τ* ∈ [0, ∞]. Technically, the data shouldn’t inform the priors, as it does here. However, this is not a concern for the extremely weak and uninformative priors used here.

We fitted the different models to data *X* of individual sessions/mice and discriminations by sampling the associated parameter posteriors, *p*(*ϕ*|*X*) ∝ *p*(*X*|*ϕ*)*p*(*ϕ*), by slice sampling (Neal, 2003). The slice sampling interval widths were set to (⟨*μ*_*n*_⟩ + 0.5)/2 for *c*, to *max*{1, ⟨*Î N*⟩}/5 for *I*_∞_, and to 10 for *τ*. The samplers were initiated by parameter values found by maximum-likelihood fits for the respective model. For each fit, we sampled four chains with 10^5^ posterior samples each, after discarding 100 burn-in samples, and keeping only each 10th sample. We used the Gelman-Rubin potential scale reduction factor (Gelman & Rubin, 1992) to assess MCMC convergence. To fit the same model to multiple discriminations simultaneously (i.e., our *pooled* fits), we sampled from the pooled posterior 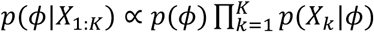, where *X*_*k*_ is the data associated with the *k* the discrimination.

We compared the fit quality of different models by the Watanabe-Akaike information criterion (WAIC; Watanabe, 2013). This criterion supports comparing models with different numbers of parameters, as it takes the associated change in model complexity into account. It is preferable to the Akaike information criterion or Bayesian information criterion, as it provides a better approximation to the cross-validated predictive density than other methods (Gelman, Hwang, & Vehtari, 2014).

We found posterior predictive densities by empirically marginalizing over the posterior parameter samples, *ϕ*^(1)^, …, *ϕ*^(*J*)^, pooled across all four chains. That is, we approximated the density of any function *f*(*ϕ*) of these parameters by 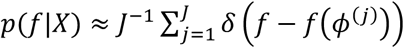, where *δ*(·) is the Dirac delta function. This approach was used to find the predictive density of the fitted information increase in **Fig. 4a** (top), as well as the information in **Fig. 4a** (bottom) and **Fig. 4c**. We also used it to estimate the posterior distribution of the required population size *N*_95_ to capture 95% of the asymptotic information.

### Additional data analysis and statistical tests

Except for **Figs. 6** and **7**, all statistical tests across sessions/mice were restricted to mice 1-4.

**Figure 3**. We removed noise correlations in the recorded data by, for each neuron, randomly permuting the trial order across all trials in which the same drift direction was presented. We then compared the total information in the recorded population with 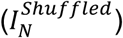 and without (*I*_*N*_) trial-shuffling by a bootstrap test (**Fig. 3d**). To do so, we estimated mean and variance of that total recorded information as described above, and then computed the probability of the null hypotheses 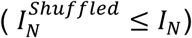 by 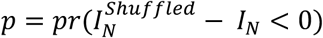, where we assumed Gaussian information estimates. We compared 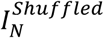 to *I*_*N*_ across sessions/mice by a paired t-test across all non-overlapping discriminations with *δθ* = 45° (**Fig. 3d**). We focused exclusively on discriminations that didn’t share any drift directions, to avoid comparing estimates that rely on the same underlying set of trials. Unless otherwise noted, all non-overlapping discriminations with *δθ* = 45° were performed on the 0° vs. 45°, 90° vs. 135°, 180° vs. 225°, and 270° vs. 315° discriminations. To test for significant differences in the drift direction discrimination thresholds (**Fig. 3f**) across multiple discriminations with the same difference in drift directions, *δθ*, we relied on the one-to-one mapping between information and discrimination threshold, and performed the test directly on the estimated information. For *K*discriminations (in our case *K* = 4 for non-overlapping discriminations), let *I*_*N,k*_, *k* = 1, …, *K* denote the information in the recorded population for discrimination 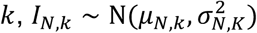. To test the null hypothesis that all *I*_*N,k*_ share the same mean, we drew 10^5^ bootstrap samples each from 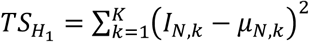 and 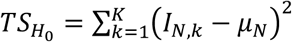 with 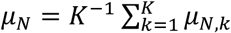, and then computed the probability that 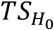 is larger than 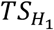 by 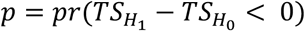.

**Figure 4**. To test how 1/*I*_*N*_ scales with 1/*N* (**Fig. 4b**), we found the moments of 1/*I*_*N*_ by ⟨1/*I*_*N*_⟩ ≈ 1/⟨*I*_*N*_⟩ and var(1/*I*_*N*_) ≈ *var*(*I*_*N*_)/⟨*I*_*N*_⟩^4^. To fit ⟨1/*I*_*N*_⟩ over 1/*N*, we performed weighted linear regression with weights 1/var(1/*I*_*N*_) for each *N*. The pooling across different discriminations in **Fig. 4d** was performed over 45° vs. 90°, 135° vs. 180°, 225° vs. 270°, and 0° vs. 315° for *pooled 1*, and 0° vs. 45°, 90° vs. 135°, 180° vs. 225°, and 270° vs. 315° for *pooled 2*. All other pooled estimates (**Figs. 4e, 6d** and **e**, and **7b**) were pooled across 45° vs. 90°, 135° vs. 180°, 225° vs. 270°, and 0° vs. 315° for *δθ* = 45°, across 45° vs. 135°, 90° vs. 180°, 225° vs. 315°, and 0° vs. 270° for *δθ* = 90°, and across 45° vs. 180°, 90° vs. 315°, and 0° vs. 225° for *δθ* = 135°. Note that the estimate *I*_*N*_’s are correlated across different *N*’s, and we did not correct for these correlations. Such a correction might lower the reported *R*^2^ values. Therefore, the Bayesian model comparison across different information scaling models, as reported in the main text, provides a statistically sounder confirmation of limited asymptotic information.

**Figure 5**. The shaded error regions in **Fig. 5a** relied on parametric bootstrap estimates. For information scaling for a fixed ordering, we computed the estimate and variance of *I*_1_, *I*_2_, … by the Fisher information and the variance of this estimator (see SI), and used these estimates to compute mean and variance of the information increase associated with adding individual neurons to the population. We then re-sampled these information increases from Gaussian distributions with the found moments, and summed the individual samples to find different samples for the whole information scaling curve. These samples were in turn used to estimate mean and variance of the information scaling for a fixed order with which neurons were added to the population. This procedure was chosen, as the increase in Fisher information is independent across added neurons, whereas the total Fisher information is not. A similar procedure was used to find the estimates for random orderings, for which we additionally shuffled the order of neurons across different samples of the information scaling curve. The above procedures yielded 10^3^ bootstrap samples for each information scaling curve, which we in turn used to find samples for the population sizes required to capture 90% of the total information (**Figs. 5a** and **b**). In neither case did we apply bias-correction of the Fisher information estimate. This bias correction would have been stronger for larger population sizes, which would have led to a seeming (but not real) drop of information with population size, resulting from a lower number of trials per neuron in the population, and an associated stronger bias correction.

**Figure 6**. To identify for individual discriminations if increasing the stimulus contrast increased information in the recorded population (**Figs. 6a** and **b**), we estimated information in the recorded population by the bias-corrected Fisher information estimate (Kanitscheider, Coen-Cagli, Kohn, et al., 2015), and its variance by our analytical expression for this estimate’s variance (see SI). We assumed the estimate for low and high contrast, *I*_*N*_^*LO*^ and *I*_*N*_^*HI*^, to be Gaussian, and found the probability of no information increase by *pr*(*I*_*N*_^*HI*^ ≤ *I*_*N*_^*LO*^), using the aforementioned moments. The paired t-test across sessions/mice (**Fig. 6b**) did not take into account the information estimates’ variance. For **Fig. 6e**, higher contrast was considered to significantly increase the information in the recorded population (filled dots in **Fig. 6e**), if it did so for at least five out of eight possible discriminations with *δθ* = 45°.

**Figure 7**. To test the relationship between *c* and *I*_∞_ in **Fig. 7d**, we performed the linear regression *log*_10_(*c*) = *β*_0_ + *β*_1_*log*_10_(*I*_∞_). The relationship between *N*_95_ and *I*_∞_ was found by substituting 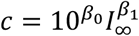 into the expression for *N*_95_, resulting in 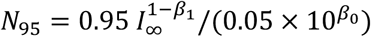. To find the information loss for using a smaller population size than required, we assumed 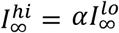 and computed the fraction 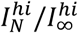 at 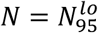, which is the population size that captures 95% of 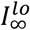. Substituting the found relationships between *I*_∞_, *c*, and *N*_95_ results in this fraction to be given by 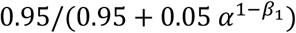, which, for *α* = 3, equals 0.93. Interestingly, this fraction depends only the relationship between 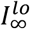 and 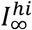, as quantified by *α*, but not on their individual values.

**Figure 8**. All estimates in **Fig. 8** are averages across 10 random splits of the recorded data. For each split, half of the trials were used to compute the principal dimensions, ***Q***_*train*_, using the spectral decomposition 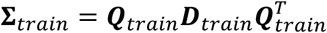, where ***D***_*train*_ is diagonal, ***Q***_*train*_ is the matrix of unit eigenvectors, and we denote the *n*th column vector of ***Q***_*train*_ by ***q***_*n,train*_. The second half of trials was used to find 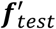 and Σ_*test*_, from which we computed the shown estimates as follows. The noise variance associated with the *n*th principal dimension was found by 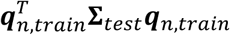. The ***f***′ alignment to the *n*th principal dimension was found by 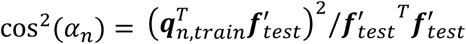. The information encoded in the first *n* principal dimensions was found by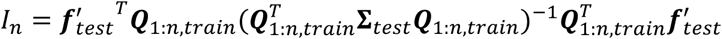, where ***Q***_1:*n,train*_ is the matrix formed by the first *n* columns of ***Q***_*train*_.

## Discussion

To compare the estimated population sizes to the number of neurons in V1, we asked for the number of neurons required to encode 95% of the asymptotic information associated with a direction discrimination threshold of 1°. This threshold most likely exceeds the behavioral performance that mice can reach even for high contrast stimuli (Abdolrahmani et al., 2019; Glickfeld et al., 2013) and thus provides an upper bound on the required population size. Achieving such a low threshold requires an asymptotic information of 4651 rad^-2^ (**Fig. 3e**), and approximately 48,000 neurons are necessary to encode 95% of this information (**Fig. 7d**). Current estimates of the neural density of mouse V1 range from 92,400 to 214,000 neurons per mm^3^ (Herculano-Houzel et al., 2013; Keller et al., 2018). For area V1 with an approximate size of 3.063mm^3^ (Herculano-Houzel et al., 2013), this amounts to 283,000 to 655,500 neurons (Keller et al., 2018). Therefore, our estimated population sizes are well within those available in V1 of mice.

## Acknowledgments

We would like to thank members of the HMS Neurobiology department for useful discussions and feedback on the work, and Rachel Wilson and Richard Born for comments on early versions of the manuscript. The work was supported by a scholar award from the James S. McDonnell Foundation (grant# 220020462, J.D.), grants from the NIH (R01MH115554, J.D.; R01MH107620, C.D.H.; R01NS089521, C.D.H.; R01NS108410, C.D.H.), the NSF’s NeuroNex program (DBI-1707398. R.N.), MINECO (Spain; BFU2017-85936-P, R.M.-B.), the Howard Hughes Medical Institute (HHMI, ref 55008742, R.M.-B.), the ICREA Academia (2016, R.M.-B.), the Government of Aragon (Spain; ISAAC lab, cod T33 17D, I.A.-R.), the Spanish Ministry of Economy and Competitiveness (TIN2016-80347-R, I.A.-R.), the Gatsby Charitable Foundation (R.N.), and an NSF Graduate Research Fellowship (A.J.).

## Author Contributions

All authors designed the research and wrote the paper; A.J. and S.N.C. performed the experiments; M.K., R.N., I.A.-R., R.M.-B, and J.D. developed the theory; M.K., A.J., S.N.C., and J.D. analyzed the data.

## References

Abbott, L. F., & Dayan, P. (1999). The effect of correlated variability on the accuracy of a population code. Neural Computation, 11(1), 91–101. https://doi.org/10.1162/089976699300016827

Abdolrahmani, M., Lyamzin, D. R., Aoki, R., & Benucci, A. (2019). Cognitive modulation of interacting corollary discharges in the visual cortex. BioRxiv, 615229. https://doi.org/10.1101/615229

Acerbi, L., Vijayakumar, S., & Wolpert, D. M. (2014). On the Origins of Suboptimality in Human Probabilistic Inference. PLoS Computational Biology, 10(6), e1003661. https://doi.org/10.1371/journal.pcbi.1003661

Adibi, M., McDonald, J. S., Clifford, C. W. G., & Arabzadeh, E. (2013). Adaptation Improves Neural Coding Efficiency Despite Increasing Correlations in Variability. Journal of Neuroscience, 33(5), 2108–2120. https://doi.org/10.1523/JNEUROSCI.3449-12.2013

Averbeck, B. B., Latham, P. E., & Pouget, A. (2006). Neural correlations, population coding and computation. Nature Reviews Neuroscience, 7(5), 358–366. https://doi.org/10.1038/nrn1888

Averbeck, B. B., & Lee, D. (2003). Neural Noise and Movement-Related Codes in the Macaque Supplementary Motor Area. The Journal of Neuroscience, 23(20), 7630–7641. https://doi.org/10.1523/JNEUROSCI.23-20-07630.2003

Averbeck, B. B., & Lee, D. (2006). Effects of Noise Correlations on Information Encoding and Decoding. Journal of Neurophysiology, 95(6), 3633–3644. https://doi.org/10.1152/jn.00919.2005

Beck, J. M., Ma, W. J., Pitkow, X., Latham, P. E., & Pouget, A. (2012). Not Noisy, Just Wrong: The Role of Suboptimal Inference in Behavioral Variability. Neuron, 74(1), 30–39. https://doi.org/10.1016/j.neuron.2012.03.016

Britten, K. H., Newsome, W. T., Shadlen, M. N., Celebrini, S., & Movshon, J. A. (1996). A relationship between behavioral choice and the visual responses of neurons in macaque MT. Visual Neuroscience, 13(01), 87–100. https://doi.org/10.1017/S095252380000715X

Britten, K. H., Shadlen, M. N., Newsome, W. T., & Movshon, J. a. (1992). The analysis of visual motion: a comparison of neuronal and psychophysical performance. The Journal of Neuroscience, 12(12), 4745–4765. https://doi.org/10.1.1.123.9899

Busse, L., Ayaz, A., Dhruv, N. T., Katzner, S., Saleem, A. B., Scholvinck, M. L., … Carandini, M. (2011). The Detection of Visual Contrast in the Behaving Mouse. Journal of Neuroscience, 31(31), 11351–11361. https://doi.org/10.1523/JNEUROSCI.6689-10.2011

Carandini, M. (2004). Amplification of Trial-to-Trial Response Variability by Neurons in Visual Cortex. PLoS Biology, 2(9), e264. https://doi.org/10.1371/journal.pbio.0020264

Chen, T.-W., Wardill, T. J., Sun, Y., Pulver, S. R., Renninger, S. L., Baohan, A., … Kim, D. S. (2013). Ultrasensitive fluorescent proteins for imaging neuronal activity. Nature, 499(7458), 295–300. https://doi.org/10.1038/nature12354

Chen, Y., Geisler, W. S., & Seidemann, E. (2006). Optimal decoding of correlated neural population responses in the primate visual cortex. Nature Neuroscience, 9(11), 1412–1420. https://doi.org/10.1038/nn1792

Chettih, S. N., & Harvey, C. D. (2019). Single-neuron perturbations reveal feature-specific competition in V1. Nature, 567(7748), 334–340. https://doi.org/10.1038/s41586-019-0997-6

Cohen, M. R., & Kohn, A. (2011). Measuring and interpreting neuronal correlations. Nature Neuroscience, 14(7), 811–819. https://doi.org/10.1038/nn.2842

Cotton, R. J., Ecker, A. S., Froudarakis, E., Berens, P., Bethge, M., Saggau, P., & Tolias, A. S. (2018). Accuracy of sensory information does not saturate for large neuronal populations. 2018 Neuroscience Meeting Planner, 219.02/BB10. San Diego, CA: Society for Neuroscience.

Cover, T. M., & Thomas, J. A. (2006). Elements of Information Theory (2nd Editio). Wiley.

Denman, D. J., & Reid, R. C. (2019). Synergistic population encoding and precise coordinated variability across interlaminar ensembles in the early visual system. BioRxiv, 1–24. https://doi.org/10.1101/812859

Dow, B. M. (2002). Orientation and Color Columns in Monkey Visual Cortex. Cerebral Cortex, 12(10), 1005–1015. https://doi.org/10.1093/cercor/12.10.1005

Doya, K., Ishii, S., Pouget, A., & Rao, R. P. N. (2006). Bayesian Brain: Probabilistic Approaches to Neural Coding. MIT Press.

Drugowitsch, J., Wyart, V., Devauchelle, A.-D., & Koechlin, E. (2016). Computational Precision of Mental Inference as Critical Source of Human Choice Suboptimality. Neuron, 92(6), 1–14. https://doi.org/10.1016/j.neuron.2016.11.005

Ecker, A. S., Berens, P., Tolias, A. S., & Bethge, M. (2011). The Effect of Noise Correlations in Populations of Diversely Tuned Neurons. Journal of Neuroscience, 31(40), 14272–14283. https://doi.org/10.1523/JNEUROSCI.2539-11.2011

Engel, T. A., & Steinmetz, N. A. (2019). New perspectives on dimensionality and variability from large-scale cortical dynamics. Current Opinion in Neurobiology, 58, 181–190. https://doi.org/10.1016/j.conb.2019.09.003

Faisal, A. A., Selen, L. P. J., & Wolpert, D. M. (2008). Noise in the nervous system. Nature Reviews Neuroscience, 9(4), 292–303. https://doi.org/10.1038/nrn2258

Fiser, J., Berkes, P., Orbán, G., & Lengyel, M. (2010). Statistically optimal perception and learning: from behavior to neural representations. Trends in Cognitive Sciences, 14(3), 119–130. https://doi.org/10.1016/j.tics.2010.01.003

Ganguli, D., & Simoncelli, E. P. (2014). Efficient Sensory Encoding and Bayesian Inference with Heterogeneous Neural Populations. Neural Computation, 26(10), 2103–2134. https://doi.org/10.1162/NECO_a_00638

Gao, P., & Ganguli, S. (2015). On simplicity and complexity in the brave new world of large-scale neuroscience. Current Opinion in Neurobiology, 32, 148–155. https://doi.org/10.1016/j.conb.2015.04.003

Gao, P., Trautmann, E., Yu, B. M., Santhanam, G., Ryu, S., Shenoy, K., & Ganguli, S. (2017). A theory of multineuronal dimensionality, dynamics and measurement. BioRxiv, 214262. https://doi.org/10.1101/214262

Gelman, A., Hwang, J., & Vehtari, A. (2014). Understanding predictive information criteria for Bayesian models. Statistics and Computing, 24(6), 997–1016. https://doi.org/10.1007/s11222-013-9416-2

Gelman, A., & Rubin, D. B. (1992). Inference from Iterative Simulation Using Multiple Sequences. Statistical Science, 7(4), 457–472. https://doi.org/10.1214/ss/1177011136

Glickfeld, L. L., Histed, M. H., & Maunsell, J. H. R. (2013). Mouse Primary Visual Cortex Is Used to Detect Both Orientation and Contrast Changes. Journal of Neuroscience, 33(50), 19416–19422. https://doi.org/10.1523/JNEUROSCI.3560-13.2013

Green, D. M., & Swets, J. A. (1966). Signal Detection Theory and Psychophysics. New York: Whiley.

Gu, Y., Liu, S., Fetsch, C. R., Yang, Y., Fok, S., Sunkara, A., … Angelaki, D. E. (2011). Perceptual learning reduces interneuronal correlations in macaque visual cortex. Neuron, 71(4), 750–761. https://doi.org/10.1016/j.neuron.2011.06.015

Haefner, R. M., Gerwinn, S., Macke, J. H., & Bethge, M. (2013). Inferring decoding strategies from choice probabilities in the presence of correlated variability. Nature Neuroscience, 16(2), 235–242. https://doi.org/10.1038/nn.3309

Harvey, C. D., Coen, P., & Tank, D. W. (2012). Choice-specific sequences in parietal cortex during a virtual-navigation decision task. Nature, 484(7392), 62–68. https://doi.org/10.1038/nature10918

Herculano-Houzel, S., Watson, C., & Paxinos, G. (2013). Distribution of neurons in functional areas of the mouse cerebral cortex reveals quantitatively different cortical zones. Frontiers in Neuroanatomy, 7(October), 1–14. https://doi.org/10.3389/fnana.2013.00035

Ince, R. A. A., Panzeri, S., & Kayser, C. (2013). Neural Codes Formed by Small and Temporally Precise Populations in Auditory Cortex. Journal of Neuroscience, 33(46), 18277–18287. https://doi.org/10.1523/JNEUROSCI.2631-13.2013

Jasper, A. I., Tanabe, S., & Kohn, A. (2019). Predicting Perceptual Decisions Using Visual Cortical Population Responses and Choice History. The Journal of Neuroscience, 39(34), 6714–6727. https://doi.org/10.1523/JNEUROSCI.0035-19.2019

Kanitscheider, I., Coen-Cagli, R., Kohn, A., & Pouget, A. (2015). Measuring Fisher Information Accurately in Correlated Neural Populations. PLoS Computational Biology, 11(6), 1–27. https://doi.org/10.1371/journal.pcbi.1004218

Kanitscheider, I., Coen-Cagli, R., & Pouget, A. (2015). Origin of information-limiting noise correlations. Proceedings of the National Academy of Sciences of the United States of America, 112(50), E6973–82. https://doi.org/10.1073/pnas.1508738112

Keller, D., Erö, C., & Markram, H. (2018). Cell Densities in the Mouse Brain: A Systematic Review. Frontiers in Neuroanatomy, 12(October). https://doi.org/10.3389/fnana.2018.00083

Kobak, D., Brendel, W., Constantinidis, C., Feierstein, C. E., Kepecs, A., Mainen, Z. F., … Machens, C. K. (2016). Demixed principal component analysis of neural population data. ELife, 5(APRIL2016), 1–36. https://doi.org/10.7554/eLife.10989

Kohn, A., Coen-cagli, R., Kanitscheider, I., & Pouget, A. (2016). Correlations and Neuronal Population Information. Annual Review of Neuroscience, 39(April), 237–256. https://doi.org/10.1146/annurev-neuro-070815-013851

Leavitt, M. L., Pieper, F., Sachs, A. J., & Martinez-Trujillo, J. C. (2017). Correlated variability modifies working memory fidelity in primate prefrontal neuronal ensembles. Proceedings of the National Academy of Sciences, 114(12), E2494–E2503. https://doi.org/10.1073/pnas.1619949114

Ledochowitsch, P., Huang, L., Knoblich, U., Oliver, M., Reid, C., Li, L., … Buice, M. A. (2019). On the correspondence of electrical and optical physiology in in vivo population-scale two-photon calcium imaging. BioRxiv, 800102. https://doi.org/10.1101/800102

Maynard, E. M., Hatsopoulos, N. G., Ojakangas, C. L., Acuna, B. D., Sanes, J. N., Normann, R. A., & Donoghue, J. P. (1999). Neuronal Interactions Improve Cortical Population Coding of Movement Direction. The Journal of Neuroscience, 19(18), 8083–8093. https://doi.org/10.1523/JNEUROSCI.19-18-08083.1999

Mendels, O. P., & Shamir, M. (2018). Relating the Structure of Noise Correlations in Macaque Primary Visual Cortex to Decoder Performance. Frontiers in Computational Neuroscience, 12(March). https://doi.org/10.3389/fncom.2018.00012

Moreno-Bote, R., Beck, J., Kanitscheider, I., Pitkow, X., Latham, P., & Pouget, A. (2014). Information-limiting correlations. In Nature Neuroscience (Vol. 17). https://doi.org/10.1038/nn.3807

Moreno-Bote, R., Knill, D. C., & Pouget, A. (2011). Bayesian sampling in visual perception. Proc Natl Acad Sci U S A, 108(30), 12491–12496. https://doi.org/10.1073/pnas.1101430108

Mott, M. C., Gordon, J. A., & Koroshetz, W. J. (2018). The NIH BRAIN Initiative: Advancing neurotechnologies, integrating disciplines. PLOS Biology, 16(11), e3000066. https://doi.org/10.1371/journal.pbio.3000066

Ni, A. M., Ruff, D. A., Alberts, J. J., Symmonds, J., & Cohen, M. R. (2018). Learning and attention reveal a general relationship between population activity and behavior. Science, 359(6374), 463–465. https://doi.org/10.1126/science.aao0284

Nogueira, R., Peltier, N. E., Anzai, A., DeAngelis, G. C., Martínez-Trujillo, J., & Moreno-Bote, R. (2019). The effects of population tuning and trial-by-trial variability on information encoding and behavior. The Journal of Neuroscience, 0859–19. https://doi.org/10.1523/JNEUROSCI.0859-19.2019

Otazu, G. H., Tai, L.-H., Yang, Y., & Zador, A. M. (2009). Engaging in an auditory task suppresses responses in auditory cortex. Nature Neuroscience, 12(5), 646–654. https://doi.org/10.1038/nn.2306

Peirce, J. W. (2007). PsychoPy—Psychophysics software in Python. Journal of Neuroscience Methods, 162(1–2), 8–13. https://doi.org/10.1016/j.jneumeth.2006.11.017

Pitkow, X., Liu, S., Angelaki, D. E., DeAngelis, G. C., & Pouget, A. (2015). How Can Single Sensory Neurons Predict Behavior? Neuron, 87(2), 411–424. https://doi.org/10.1016/j.neuron.2015.06.033

Pouget, A., Beck, J. M., Ma, W. J., & Latham, P. E. (2013). Probabilistic brains: knowns and unknowns. Nature Neuroscience, 16(9), 1170–1178. https://doi.org/10.1038/nn.3495

Pruszynski, J. A., & Zylberberg, J. (2019). The language of the brain: real-world neural population codes. Current Opinion in Neurobiology, 58, 30–36. https://doi.org/10.1016/j.conb.2019.06.005

Ringach, D. L., Mineault, P. J., Tring, E., Olivas, N. D., Garcia-Junco-Clemente, P., & Trachtenberg, J. T. (2016). Spatial clustering of tuning in mouse primary visual cortex. Nature Communications, 7(1), 12270. https://doi.org/10.1038/ncomms12270

Semedo, J. D., Zandvakili, A., Machens, C. K., Yu, B. M., & Kohn, A. (2019). Cortical Areas Interact through a Communication Subspace. Neuron, 102, 1–11. https://doi.org/10.1016/j.neuron.2019.01.026

Seriès, P., Latham, P. E., & Pouget, A. (2004). Tuning curve sharpening for orientation selectivity: coding efficiency and the impact of correlations. Nature Neuroscience, 7(10), 1129–1135. https://doi.org/10.1038/nn1321

Shadlen, M. N., & Newsome, W. T. (1998). The variable discharge of cortical neurons: implications for connectivity, computation, and information coding. The Journal of Neuroscience: The Official Journal of the Society for Neuroscience, 18(10), 3870–3896. https://doi.org/0270-6474/98/183870-27$05.00/0

Shamir, M. (2014). Emerging principles of population coding: in search for the neural code. Current Opinion in Neurobiology, 25, 140–148. https://doi.org/10.1016/j.conb.2014.01.002

Softky, W., & Koch, C. (1993). The highly irregular firing of cortical cells is inconsistent with temporal integration of random EPSPs. The Journal of Neuroscience, 13(1), 334–350. https://doi.org/10.1523/JNEUROSCI.13-01-00334.1993

Stringer, C., Michaelos, M., & Pachitariu, M. (2019). High precision coding in mouse visual cortex. BioRxiv, 679324. https://doi.org/10.1101/679324

Tolhurst, D. J., Movshon, J. A., & Dean, A. F. (1983). The statistical reliability of signals in single neurons in cat and monkey visual cortex. Vision Research, 23(8), 775–785. https://doi.org/10.1016/0042-6989(83)90200-6

Watanabe, S. (2013). A widely applicable bayesian information criterion. Journal of Machine Learning Research, 14(1), 867–897.

Whiteway, M. R., Bartolo, R., Averbeck, B. B., & Butts, D. A. (2017). Unsupervised nonlinear dimensionality reduction of large-scale neural recordings in prefrontal cortex. 2017 Neuroscience Meeting Planner, 249.07/RR3. Washington, DC: The Society of Neuroscience.

Williamson, R. C., Cowley, B. R., Litwin-Kumar, A., Doiron, B., Kohn, A., Smith, M. A., & Yu, B. M. (2016). Scaling Properties of Dimensionality Reduction for Neural Populations and Network Models. PLOS Computational Biology, 12(12), e1005141. https://doi.org/10.1371/journal.pcbi.1005141

Zohary, E., Shadlen, M. N., & Newsome, W. T. (1994). Correlated neuronal discharge rate and its implications for psychophysical performance. Nature, 370(6485), 140–143. https://doi.org/10.1038/370140a0

