## Supplementary Information for "Scaling of information in large neural populations reveals signatures of information-limiting correlations"

**5 Supplementary Tables** **19**

**6 Supplementary Figures** **20**

### 1 Generalized Fisher information

Fisher information quantifies how much neural activity $\boldsymbol{r}$ tells us about a stimulus $\theta$ around a particular reference $\theta_0$. As such, it is a measure of fine discrimination performance. Here, we show how *linear* Fisher information relates to Fisher information in general, show how it can be generalized beyond fine discrimination, and describe some properties of this generalization.

#### 1.1 Definition and properties of linear Fisher information

We can derive *linear* Fisher information in two ways [Moreno-Bote et al., 2014, Ganguli and Simoncelli, 2014]. The first is to assume that $p(\boldsymbol{r}|\theta)$ is a member of the exponential family with linear sufficient statistics. The second is to show that it is the Fisher information that can be extracted with a minimum-variance unbiased linear decoder. We will provide both derivations in turn.

Let us first assume that neural activity $\boldsymbol{r}$ in response to a stimulus $\theta$ follows an exponential family distribution with linear sufficient statistics,

$$p(\boldsymbol{r}|\theta) = g(\theta)\Phi(\boldsymbol{r})\exp(\boldsymbol{h}(\theta)^T\boldsymbol{r}), \tag{1}$$

where

$$g(\theta) = \frac{1}{\int \Phi(\boldsymbol{r})\exp(\boldsymbol{h}(\theta)^T\boldsymbol{r})d\boldsymbol{r}}, \tag{2}$$

in which $g(\theta)$, $\Phi(\boldsymbol{r})$, and $\boldsymbol{h}(\theta)$ are known functions.

The partial derivative with respect to $\theta$ of the log-likelihood function, $\frac{\partial}{\partial\theta}\log p(\boldsymbol{r}|\theta)$, is called the "score" which is given by

$$\frac{\partial}{\partial\theta}\log p(\boldsymbol{r}|\theta) = \boldsymbol{h}'^T(\theta)(\boldsymbol{r}(\theta) - \boldsymbol{f}(\theta)), \tag{3}$$

where $\boldsymbol{f}(\theta) = \mathbb{E}\left(\boldsymbol{r}(\theta)\right)$ is the population activity vector. Note that the first moment of the score function is zero.

The Fisher information can be derived using the variance of the score function [Cover and Thomas, 2006] which can be written as follows:

$$I(\theta) = \mathbb{E}\left[\left(\frac{\partial}{\partial\theta}\log p(\boldsymbol{r}|\theta)\right)^2\right] = \boldsymbol{h}'(\theta)^T\boldsymbol{\Sigma}(\theta)\boldsymbol{h}'(\theta), \tag{4}$$

where $\boldsymbol{\Sigma}(\theta) = \mathbb{E}\left[\left(\boldsymbol{r}(\theta) - \boldsymbol{f}(\theta)\right)\left(\boldsymbol{r}(\theta) - \boldsymbol{f}(\theta)\right)^T\right]$ is the noise covariance matrix. To express the Fisher information in terms of $\boldsymbol{f}(\theta)$, we note that

$$\boldsymbol{f}'(\theta) = \frac{\mathrm{d}}{\mathrm{d}\theta}\int \boldsymbol{r}p(\boldsymbol{r}|\theta)d\boldsymbol{r} = \boldsymbol{\Sigma}(\theta)\boldsymbol{h}'(\theta). \tag{5}$$

Thus, we have $\boldsymbol{h}'(\theta) = \boldsymbol{\Sigma}^{-1}(\theta)\boldsymbol{f}'(\theta)$ [Ma et al., 2006]. Taking this expression to substitute both instances of $\boldsymbol{h}'(\theta)$ in the Fisher information results in

$$I(\theta) = \boldsymbol{f}'(\theta)^T\boldsymbol{\Sigma}^{-1}(\theta)\boldsymbol{f}'(\theta). \tag{6}$$

To show that linear Fisher information is the information extractable by a minimum-variance unbiased linear decoder, assume that the decoder linearly combines neural activity of neurons with a projection vector $\boldsymbol{w}$. For fine discrimination task with two close-by stimuli $\theta_1 = \theta_0 - \delta\theta$ and $\theta_2 = \theta_0 + \delta\theta$ with small $\delta\theta$, the unbiased locally linear estimator for $\hat{\theta}$ is given by

$$\hat{\theta} - \theta_0 = \boldsymbol{w}^T(\boldsymbol{r} - \boldsymbol{f}(\theta_0)). \tag{7}$$

The expectation of the right-hand side around $\theta_0$ is $\boldsymbol{w}^T\left(\langle\boldsymbol{r}\rangle - \boldsymbol{f}(\theta_0)\right) = 0$, demonstrating that the estimator is unbiased. Our aim is to find a $w$ that yields a locally unbiased estimate, that is

$$\frac{\mathrm{d}\mathbb{E}_\theta(\hat{\theta})}{\mathrm{d}\theta} = 1, \tag{8}$$

imposing the constraint

$$\boldsymbol{w}^T\boldsymbol{f}'(\theta) = 1. \tag{9}$$

To find the minimum variance estimator satisfying this constraint, note that its variance is given by var $\left(\hat{\theta}\right) = \boldsymbol{w}^T \boldsymbol{\Sigma} \boldsymbol{w}$, where $\boldsymbol{\Sigma}$ is the noise covariance matrix around $\theta_0$. Therefore, we aim to find

$$\min_{\boldsymbol{w}} \boldsymbol{w}^T \boldsymbol{\Sigma} \boldsymbol{w}, \qquad \text{s.t. } \boldsymbol{w}^T \boldsymbol{f}'(\theta) = 1. \tag{10}$$

Using a Lagrange multiplier to solve the constraint optimization for $\boldsymbol{w}$ results in

$$\boldsymbol{w}^* = \frac{\boldsymbol{\Sigma}^{-1} \boldsymbol{f'}}{\boldsymbol{f'}^T \boldsymbol{\Sigma}^{-1} \boldsymbol{f'}}, \tag{11}$$

with associated estimator variance

$$\text{var}\left(\hat{\theta}\right) = \frac{1}{\boldsymbol{f'}^T \boldsymbol{\Sigma}^{-1} \boldsymbol{f'}}. \tag{12}$$

By the Cramér-Rao bound [Cover and Thomas, 2006], the Fisher information is the inverse of this variance, resulting in

$$I(\theta) = \frac{1}{\text{var}\left(\hat{\theta}\right)} = \boldsymbol{f'}^T \boldsymbol{\Sigma}^{-1} \boldsymbol{f'}, \tag{13}$$

which matches the previously derived expression for the linear Fisher information. This demonstrates that linear Fisher information can be interpreted in multiple ways: it is either the Fisher information when restricting the distribution of neural activity to the exponential family with linear sufficient statistics (which contains independent-Poisson populations with dense tuning curves, as well as other distributions [Ma et al., 2006], or the Fisher information that can be extracted with a linear decoder.

#### 1.2 Generalizing Fisher information beyond fine discrimination

Let us generalize the above to coarse discrimination. To do so, assume two classes, $C_1$ and $C_2$, which represent a pair of stimulus orientations at $\theta_1$ and $\theta_2$ in the experiment. As before, we will derive generalized Fisher information in two ways. First, we will derive it by making particular distributional assumptions on $p(\boldsymbol{r}|\theta_1)$ and $p(\boldsymbol{r}|\theta_2)$. Then, we will derive it from the perspective of optimal linear discrimination.

For the first approach, assume that $p(\boldsymbol{r}|\theta_j)$ for both $j \in \{1, 2\}$ follows a Gaussian distribution,

$$\begin{aligned} C_1 &: \; p(\boldsymbol{r}|\theta_1) = \mathcal{N}(\boldsymbol{r}|\boldsymbol{f}_1, \boldsymbol{\Sigma}) \\ C_2 &: \; p(\boldsymbol{r}|\theta_2) = \mathcal{N}(\boldsymbol{r}|\boldsymbol{f}_2, \boldsymbol{\Sigma}), \end{aligned} \tag{14}$$

which have different means, but the same covariance matrix. Under the assumption that $\theta$ is a random variable (which takes two values, $\theta \in \{\theta_1, \theta_2\}$, in coarse discrimination tasks), it is easy to find a decision rule that minimize the expected Bayes risk [Berger, 1993]. We will denote $L_{ij}$ as the loss of choosing $C_j$ when $C_i$ is correct. Furthermore, we assume a symmetric decision problem with symmetric loss, that is $L_{12} = L_{21}$ and $L_{11} = L_{22}$, a uniform prior $p(C_1) = p(C_2) = 1/2$, and a preference for making correct choices, that is $L_{11} < L_{12}$. In this case, the expected Bayesian risk, $\sum_{i \in \{1,2\}} L_{i\mathcal{D}(\boldsymbol{r})} p(C_i|\boldsymbol{r})$, associated with decision rule $\mathcal{D}(\boldsymbol{r}) \in \{1, 2\}$ is minimized by

$$\mathcal{D}\left(\boldsymbol{r}\right) = \begin{cases} 2 & \text{if } \Lambda(\boldsymbol{r}) = \log \frac{p(\boldsymbol{r}|\theta_2)}{p(\boldsymbol{r}|\theta_1)} > 0, \\ 1 & \text{otherwise }, \end{cases} \tag{15}$$

where $\Lambda(\boldsymbol{r})$ is the log-likelihood ratio. For the assumed Gaussian likelihoods, this log-likelihood ratio is given by

$$\Lambda(\boldsymbol{r}) = (\boldsymbol{f}_2 - \boldsymbol{f}_1)^T \boldsymbol{\Sigma}^{-1} (\boldsymbol{r} - \boldsymbol{f}_0), \tag{16}$$

where $\boldsymbol{f}_0 = \frac{1}{2}(\boldsymbol{f}_1 + \boldsymbol{f}_2)$, and $\boldsymbol{f}_j = \mathbb{E}_{\boldsymbol{r}|\theta_j}\left(\boldsymbol{r}\right)$ for $j \in \{1, 2\}$. Letting $\boldsymbol{w} = \boldsymbol{\Sigma}^{-1} \delta \boldsymbol{f}$ with $\delta \boldsymbol{f} = \boldsymbol{f}_2 - \boldsymbol{f}_1$, we can rewrite $\Lambda(\boldsymbol{r})$ as

$$\Lambda(\boldsymbol{r}) = \boldsymbol{w}^T (\boldsymbol{r} - \boldsymbol{f}_0). \tag{17}$$

In order to identify how likely this decision rule makes the correct choice, observe that $\Lambda(\boldsymbol{r})$ follows the following distributions under $C_1$ and $C_2$,

$$\Lambda(\boldsymbol{r})|C_1 \sim \mathcal{N}\left(-\frac{1}{2}\boldsymbol{w}^T\boldsymbol{\Sigma}\boldsymbol{w}, \boldsymbol{w}^T\boldsymbol{\Sigma}\boldsymbol{w}\right), \qquad \Lambda(\boldsymbol{r})|C_2 \sim \mathcal{N}\left(\frac{1}{2}\boldsymbol{w}^T\boldsymbol{\Sigma}\boldsymbol{w}, \boldsymbol{w}^T\boldsymbol{\Sigma}\boldsymbol{w}\right). \tag{18}$$

Therefore, we can find the probability of making a correct choice under $\mathcal{D}(\boldsymbol{r})$ by

$$p(\text{correct}) = \frac{1}{2}p\left(\Lambda(\boldsymbol{r}) \leq 0|C_1\right) + \frac{1}{2}p\left(\Lambda(\boldsymbol{r}) > 0|C_2\right) = \Phi\left(\frac{1}{2}\sqrt{\boldsymbol{w}^T\boldsymbol{\Sigma}\boldsymbol{w}}\right), \tag{19}$$

where $\Phi(\cdot)$ is the cumulative function of the standard normal distribution. After replacing both instances of $\boldsymbol{w}$ by its definition, $\boldsymbol{w} = \boldsymbol{\Sigma}^{-1}\delta\boldsymbol{f}$, $p(\text{correct})$ becomes

$$p(\text{correct}) = \Phi\left(\frac{1}{2}\sqrt{\delta\boldsymbol{f}^T\boldsymbol{\Sigma}^{-1}\delta\boldsymbol{f}}\right). \tag{20}$$

Comparing this expression to Eq. (6) reveals a close similarity which we can utilize to define the generalized linear Fisher information for coarse discrimination tasks by

$$I_g(\theta) = \frac{\delta\boldsymbol{f}^T\boldsymbol{\Sigma}^{-1}\delta\boldsymbol{f}}{\delta\theta^2}, \tag{21}$$

where $\delta\theta = \theta_2 - \theta_1$ is the stimulus difference. It is easy to see that, for small $\delta\theta$, generalized linear Fisher information converges to linear Fisher information,

$$\lim_{\delta\theta\to 0} I_g(\theta) = \lim_{\delta\theta\to 0}\frac{\delta\boldsymbol{f}^T\boldsymbol{\Sigma}^{-1}\delta\boldsymbol{f}}{\delta\theta^2} = \boldsymbol{f}'^T\boldsymbol{\Sigma}^{-1}\boldsymbol{f}' = I(\theta) \tag{22}$$

As the sensitivity index $d'$ [Green and Swets, 1989] in our case is given by $d' = \sqrt{\delta\boldsymbol{f}^T\boldsymbol{\Sigma}^{-1}\delta\boldsymbol{f}}$ [Averbeck and Lee, 2006, Chen et al., 2006, Nogueira et al., 2019], the generalized linear Fisher information can be re-expressed in terms of $d'$ by

$$I_g(\theta) = \frac{d'^2}{\delta\theta^2}. \tag{23}$$

This relationship furthermore results in

$$p(\text{correct}) = \Phi\left(\frac{\delta\theta}{2}\sqrt{I_g(\theta)}\right) = \Phi\left(\frac{d'}{2}\right) \tag{24}$$

illustrating the close relationship between $p(\text{correct})$, $d'$, and $I_g(\theta)$.

An alternative derivation for generalized linear Fisher information is through an optimal linear discriminator with less stringent assumptions on the class-conditional distribution. In this second approach, we assume a linear decoder projecting the neural activity to a one-dimensional readout using

$$\hat{\theta} = \boldsymbol{w}^T\boldsymbol{r}. \tag{25}$$

To assign an observed neural activity to a class, we just need to place a threshold on the readout $\hat{\theta}$. To do so, we optimize $\boldsymbol{w}$ to maximize the class separation following Fisher's linear discriminant analysis [Berger, 2011], which minimizes the within-class variance while maximizing the between-class variance of $\boldsymbol{r}$. As before, let $\boldsymbol{f}_j$ and $\boldsymbol{\Sigma}_j$ be mean and noise covariance of neural activity in class $C_j$, but without making any further assumptions about the class-conditional densities $p(\boldsymbol{r}|C_j)$. We aim to find the $\boldsymbol{w}$ that maximizes the ratio of the between-class variance to the within-class variance, which is formulated as

$$\max_{\boldsymbol{w}}\frac{\boldsymbol{w}^T\delta\boldsymbol{f}\delta\boldsymbol{f}^T\boldsymbol{w}}{\boldsymbol{w}^T\boldsymbol{\Sigma}\boldsymbol{w}}, \qquad \text{s.t. } \|\boldsymbol{w}\|^2 = 1, \tag{26}$$

where $\delta \boldsymbol{f} \delta \boldsymbol{f}^T$ is the between-class covariance matrix and $\boldsymbol{\Sigma}$ is the average within-class covariance matrix given by

$$\boldsymbol{\Sigma} = \frac{\boldsymbol{\Sigma}_1 + \boldsymbol{\Sigma}_2}{2}. \tag{27}$$

Here, we fix $\|\boldsymbol{w}\|^2 = 1$, as we are interested in the direction of $\boldsymbol{w}$ but not its length. Using a Lagrange multiplier to solve the constraint optimization for $\boldsymbol{w}$ results in

$$\boldsymbol{w} = \frac{\boldsymbol{\Sigma}^{-1}\boldsymbol{\delta f}}{\boldsymbol{\delta f}^T\boldsymbol{\Sigma}^{-1}\boldsymbol{\delta f}}. \tag{28}$$

This yields the direction, $\boldsymbol{w}$, to best project the neural activity into one dimension.

To find the associated $p(\text{correct})$, note that $\hat{\theta}$ is the sum of a (potentially large) set of random variables. These random variables are correlated, such that the central limit theorem does not directly apply. Nonetheless, we assume this sum to be approximately Gaussian for both $\hat{\theta}|C_1$ and $\hat{\theta}|C_2$, and given by

$$\hat{\theta}|C_1 \sim \mathcal{N}\left(\boldsymbol{w}^T\boldsymbol{f_1}, \boldsymbol{w}^T\boldsymbol{\Sigma}_1\boldsymbol{w}\right), \qquad \hat{\theta}|C_2 \sim \mathcal{N}\left(\boldsymbol{w}^T\boldsymbol{f_2}, \boldsymbol{w}^T\boldsymbol{\Sigma}_2\boldsymbol{w}\right). \tag{29}$$

This results in the sensitivity index, $d'$, to be given by

$$d' = \frac{\boldsymbol{w}^T\boldsymbol{f_2} - \boldsymbol{w}^T\boldsymbol{f_1}}{\sqrt{\frac{1}{2}(\boldsymbol{w}^T\boldsymbol{\Sigma}_1\boldsymbol{w} + \boldsymbol{w}^T\boldsymbol{\Sigma}_2\boldsymbol{w})}} = \frac{\boldsymbol{w}^T\boldsymbol{\delta f}}{\sqrt{\boldsymbol{w}^T\boldsymbol{\Sigma}\boldsymbol{w}}} = \sqrt{\delta\boldsymbol{f}^T\boldsymbol{\Sigma}^{-1}\delta\boldsymbol{f}}, \tag{30}$$

yielding the same expression as before. This makes it straightforward to derive the generalized Fisher information as before.

#### 1.3 Bias-corrected generalized Fisher information

Evaluating the generalized Fisher information, Eq. (21), by replacing $\delta\boldsymbol{f}$ and $\boldsymbol{\Sigma}$ by its empirical moments estimated from neural data with a limited number of trials leads to biased estimates [Kanitscheider et al., 2015a]. In Kanitscheider et al. [2015a], they provide a bias correction for standard Fisher information, but it is unclear if this bias correction also applies to our generalization of Fisher information. In this section, we will derive such a bias correction for our generalization. This correction turns out to be the same as that provided by Kanitscheider et al. [2015a]. This is unsurprising in hindsight, as Kanitscheider et al. [2015a] do not restrict the size of $\delta\theta$ in their derivation, such that it applies to both fine and coarse discrimination.

We assume neural activity $\boldsymbol{r}_j^t,\ j = 1, 2$ in response to stimulus $\theta_j$ in trials $t = 1, ..., T$ to follow a multivariate Gaussian distribution given by

$$\boldsymbol{r}_j^t \sim \mathcal{N}\left(\boldsymbol{f}_j, \boldsymbol{\Sigma}\right), \qquad j = 1, 2, \tag{31}$$

where we assume the same covariance matrix for neural activity in response to $\theta_1$ and $\theta_2$. This is not a restriction, as our above derivation from the perspective of a linear discriminator has shown that, if these covariances differ, we can replace them by their average (which is what we do in practice, see below). Under this assumption, the empirical mean and covariance over $T$ trials for each stimulus is distributed as [Johnson and Wichern, 2007]

$$\boldsymbol{\mu}_j = \frac{1}{T}\sum_{t=1}^{T}\boldsymbol{r}_j^t \sim \mathcal{N}\left(\boldsymbol{f}_j, \frac{\boldsymbol{\Sigma}}{T}\right), \qquad \boldsymbol{S}_j = \frac{1}{T-1}\sum_{t=1}^{T}(\boldsymbol{r}_j^t - \boldsymbol{\mu}_j)(\boldsymbol{r}_j^t - \boldsymbol{\mu}_j)^T \sim \mathcal{W}\left(\frac{\boldsymbol{\Sigma}}{T-1}, T-1\right), \tag{32}$$

where $\mathcal{W}(V_{p\times p}, n)$ is the $p$-dimensional Wishart distribution with $n$ degrees of freedom.

The naïve estimation of generalized Fisher information, Eq. (21), is obtained by replacing $\delta\boldsymbol{f}$ and $\boldsymbol{\Sigma}$ with their unbiased estimates, $\delta\boldsymbol{\mu}$ and $\boldsymbol{S}$, given by

$$\delta\boldsymbol{\mu} = \boldsymbol{\mu}_1 - \boldsymbol{\mu}_2 \sim \mathcal{N}\left(\delta\boldsymbol{f}, \frac{2\boldsymbol{\Sigma}}{T}\right), \qquad \boldsymbol{S} = \frac{1}{2}(\boldsymbol{S}_1 + \boldsymbol{S}_2) \sim \mathcal{W}\left(\frac{\boldsymbol{\Sigma}}{2(T-1)}, 2(T-1)\right), \tag{33}$$

where $\mathbb{E}(\delta\boldsymbol{\mu}) = \delta\boldsymbol{f}$ and $\mathbb{E}(\boldsymbol{S}) = \boldsymbol{\Sigma}$. Furthermore, the inverse of sample covariance, $\boldsymbol{S}^{-1}$, follows an inverse Wishart distribution [Johnson and Wichern, 2007] given by

$$\boldsymbol{S}^{-1} \sim \mathcal{W}^{-1}\left(2(T-1)\boldsymbol{\Sigma}^{-1}, 2(T-1)\right), \tag{34}$$

which has mean

$$\mathbb{E}(\boldsymbol{S}^{-1}) = \frac{2(T-1)}{2T-N-3}\boldsymbol{\Sigma}^{-1} \tag{35}$$

Replacing $\delta\boldsymbol{f}$ and $\boldsymbol{\Sigma}$ with $\delta\boldsymbol{\mu}$ and $\boldsymbol{S}$ in Eq. (21) results in the following naive estimator of the generalized Fisher information to be given by

$$\hat{I}_{g,nv}(\theta) = \frac{\delta\boldsymbol{\mu}^T\boldsymbol{S}^{-1}\delta\boldsymbol{\mu}}{\delta\theta^2}. \tag{36}$$

To evaluate the bias of $I_{g,nv}$, we utilize the fact that the sample mean and sample covariance of Gaussian distributions are independent [Johnson and Wichern, 2007], such that we can express the first moment of $I_{g,nv}$ by

$$\mathbb{E}\left(\hat{I}_{g,nv}\right) = \frac{\mathbb{E}_{\delta\boldsymbol{\mu},\boldsymbol{S}}\left(\delta\boldsymbol{\mu}^T\boldsymbol{S}^{-1}\delta\boldsymbol{\mu}\right)}{\delta\theta^2}, \tag{37}$$

where

$$\begin{aligned}
\mathbb{E}_{\delta\boldsymbol{\mu},\boldsymbol{S}}\left(\delta\boldsymbol{\mu}^T\boldsymbol{S}^{-1}\delta\boldsymbol{\mu}\right) &= \mathbb{E}_{\delta\boldsymbol{\mu},\boldsymbol{S}}\left(\mathrm{Tr}\left(\delta\boldsymbol{\mu}\delta\boldsymbol{\mu}^T\boldsymbol{S}^{-1}\right)\right)\\
&= \mathrm{Tr}\left(\mathbb{E}_{\delta\boldsymbol{\mu},\boldsymbol{S}}\left(\delta\boldsymbol{\mu}\delta\boldsymbol{\mu}^T\boldsymbol{S}^{-1}\right)\right)\\
&= \mathrm{Tr}\left(\mathbb{E}_{\delta\boldsymbol{\mu}}\left(\delta\boldsymbol{\mu}\delta\boldsymbol{\mu}^T\right)\mathbb{E}_{\boldsymbol{S}}\left(\boldsymbol{S}^{-1}\right)\right)\\
&= \mathrm{Tr}\left(\left(\delta\boldsymbol{f}\delta\boldsymbol{f}^T + \frac{2\boldsymbol{\Sigma}}{T}\right)\left(\frac{2(T-1)}{2T-N-3}\boldsymbol{\Sigma}^{-1}\right)\right)\\
&= \frac{2(T-1)}{2T-N-3}\left(\mathrm{Tr}\left(\delta\boldsymbol{f}\delta\boldsymbol{f}^T\boldsymbol{\Sigma}^{-1}\right) + \frac{2N}{T}\right)\\
&= \frac{2(T-1)}{2T-N-3}\left(\delta\boldsymbol{f}^T\boldsymbol{\Sigma}^{-1}\delta\boldsymbol{f} + \frac{2N}{T}\right)\\
&= \frac{2(T-1)}{2T-N-3}\left(I_g\delta\theta^2 + \frac{2N}{T}\right).
\end{aligned} \tag{38}$$

Having the first moment of $\hat{I}_{g,nv}$, we can obtain the expression for the bias-corrected generalized Fisher information, $\hat{I}_{g,bc}$, given by

$$\hat{I}_{g,bc} = \frac{2T-N-3}{2(T-1)}\frac{\delta\boldsymbol{\mu}^T\boldsymbol{S}^{-1}\delta\boldsymbol{\mu}}{\delta\theta^2} - \frac{2N}{T\delta\theta^2}. \tag{39}$$

This estimate is the same as provided by Kanitscheider et al. [2015a], and will, in expectation, equal the true Fisher information, that is, $\mathbb{E}\left(\hat{I}_{g,bc}\right) = I_g$.

#### 1.4 Variance of bias-corrected generalized Fisher information

Let us now consider the variance of the bias-corrected generalized Fisher information across different draws of $T$ trial/samples from the same neural population. This variance has already been computed by Kanitscheider et al. [2015a], but only as a function of the true information, $I_g$, which is an unknown quantity. Here, we re-derive this expression for completeness, and additionally derive an unbiased estimated thereof as a function of $\hat{I}_{g,bc}$, which can be computed from experimental data.

The variance of $\hat{I}_{g,bc}$ is given by

$$\mathrm{var}\left(\hat{I}_{g,bc}\right) = \frac{(2T-N-3)^2}{4(T-1)^2\delta\theta^4}\mathrm{var}(\delta\boldsymbol{\mu}^T\boldsymbol{S}^{-1}\delta\boldsymbol{\mu}), \tag{40}$$

where var $\left(\delta\boldsymbol{\mu}^T\boldsymbol{S}^{-1}\delta\boldsymbol{\mu}\right)$ can be decomposed into

$$\mathrm{var}(\delta\boldsymbol{\mu}^T\boldsymbol{S}^{-1}\delta\boldsymbol{\mu}) = \mathbb{E}\left((\delta\boldsymbol{\mu}^T\boldsymbol{S}^{-1}\delta\boldsymbol{\mu})^2\right) - \mathbb{E}\left(\delta\boldsymbol{\mu}^T\boldsymbol{S}^{-1}\delta\boldsymbol{\mu}\right)^2. \tag{41}$$

The first term in Eq. (41) can be expressed as

$$\begin{aligned}\mathbb{E}\left((\delta\boldsymbol{\mu}^T\boldsymbol{S}^{-1}\delta\boldsymbol{\mu})^2\right) &= \mathbb{E}\left(\delta\boldsymbol{\mu}^T\boldsymbol{S}^{-1}\delta\boldsymbol{\mu}\delta\boldsymbol{\mu}^T\boldsymbol{S}^{-1}\delta\boldsymbol{\mu}\right)\\ &= \mathbb{E}_{\delta\boldsymbol{\mu}}\left(\delta\boldsymbol{\mu}^T\mathbb{E}_{\boldsymbol{S}}\left(\boldsymbol{S}^{-1}\delta\boldsymbol{\mu}\delta\boldsymbol{\mu}^T\boldsymbol{S}^{-1}\right)\delta\boldsymbol{\mu}\right)\\ &= \frac{4(T-1)^2}{(2T-N-3)(2T-N-5)}\mathbb{E}_{\delta\boldsymbol{\mu}}\left(\delta\boldsymbol{\mu}^T\boldsymbol{\Sigma}^{-1}\delta\boldsymbol{\mu}\delta\boldsymbol{\mu}^T\boldsymbol{\Sigma}^{-1}\delta\boldsymbol{\mu}\right)\\ &= \frac{4(T-1)^2}{(2T-N-3)(2T-N-5)}\mathbb{E}_{\delta\boldsymbol{\mu}}\left((\delta\boldsymbol{\mu}^T\boldsymbol{\Sigma}^{-1}\delta\boldsymbol{\mu})^2\right)\\ &= \frac{4(T-1)^2}{(2T-N-3)(2T-N-5)}\left(\mathrm{var}(\delta\boldsymbol{\mu}^T\boldsymbol{\Sigma}^{-1}\delta\boldsymbol{\mu}) + \mathbb{E}_{\delta\boldsymbol{\mu}}\left(\delta\boldsymbol{\mu}^T\boldsymbol{\Sigma}^{-1}\delta\boldsymbol{\mu}\right)^2\right).\end{aligned} \tag{42}$$

The second term in Eq. (41) can be expressed as

$$\mathbb{E}\left(\delta\boldsymbol{\mu}^T\boldsymbol{S}^{-1}\delta\boldsymbol{\mu}\right)^2 = \frac{4(T-1)^2}{(2T-N-3)^2}\mathbb{E}_{\delta\boldsymbol{\mu}}\left(\delta\boldsymbol{\mu}^T\boldsymbol{\Sigma}^{-1}\delta\boldsymbol{\mu}\right)^2. \tag{43}$$

Together, this results in Eq. (41) to be given by

$$\mathrm{var}(\delta\boldsymbol{\mu}^T\boldsymbol{S}^{-1}\delta\boldsymbol{\mu}) = \frac{4(T-1)^2}{(2T-N-3)(2T-N-5)}\left(\mathrm{var}(\delta\boldsymbol{\mu}^T\boldsymbol{\Sigma}^{-1}\delta\boldsymbol{\mu}) + \frac{2}{2T-N-3}\mathbb{E}_{\delta\boldsymbol{\mu}}\left(\delta\boldsymbol{\mu}^T\boldsymbol{\Sigma}^{-1}\delta\boldsymbol{\mu}\right)^2\right). \tag{44}$$

Therefore, var$(I_{g,bc})$ can be simplified to

$$\mathrm{var}\left(\hat{I}_{g,bc}\right) = \frac{2}{2T-N-5}\left(\frac{2T-N-3}{2\delta\theta^4}\mathrm{var}(\delta\boldsymbol{\mu}^T\boldsymbol{\Sigma}^{-1}\delta\boldsymbol{\mu}) + \frac{1}{\delta\theta^4}\mathbb{E}_{\delta\boldsymbol{\mu}}\left(\delta\boldsymbol{\mu}^T\boldsymbol{\Sigma}^{-1}\delta\boldsymbol{\mu}\right)^2\right). \tag{45}$$

To simplify this expression, note that if $\boldsymbol{\epsilon} \sim \mathcal{N}(\boldsymbol{\mu}, \boldsymbol{\Sigma})$, then, for a constant matrix $\boldsymbol{\Lambda}$, we have

$$\mathbb{E}(\boldsymbol{\epsilon}^T\boldsymbol{\Lambda}\boldsymbol{\epsilon}) = \mathrm{Tr}(\boldsymbol{\Lambda}\boldsymbol{\Sigma}) + \boldsymbol{\mu}^T\boldsymbol{\Lambda}\boldsymbol{\mu}. \tag{46}$$

Additionally, for a symmetric matrix $\boldsymbol{\Lambda}$, the variance of the quadratic form is expressed as

$$\mathrm{var}(\boldsymbol{\epsilon}^T\boldsymbol{\Lambda}\boldsymbol{\epsilon}) = 2\,\mathrm{Tr}\left(\boldsymbol{\Lambda}\boldsymbol{\Sigma}\boldsymbol{\Lambda}\boldsymbol{\Sigma}\right) + 4\boldsymbol{\mu}^T\boldsymbol{\Lambda}\boldsymbol{\Sigma}\boldsymbol{\Lambda}\boldsymbol{\mu}. \tag{47}$$

Applying Eqs. (46) and (47) yields

$$\mathrm{var}(\delta\boldsymbol{\mu}^T\boldsymbol{\Sigma}^{-1}\delta\boldsymbol{\mu}) = \frac{8N}{T^2} + \frac{8}{T}\delta\theta^2 I_g, \qquad \mathbb{E}_{\delta\boldsymbol{\mu}}\left(\delta\boldsymbol{\mu}^T\boldsymbol{\Sigma}^{-1}\delta\boldsymbol{\mu}\right)^2 = \frac{4N^2}{T^2} + \frac{4N}{T}\delta\theta^2 I_g + \delta\theta^4 I_g^2. \tag{48}$$

Using these expressions results in the final variance

$$\mathrm{var}\left(\hat{I}_{g,bc}\right) = \frac{2}{2T-N-5}\left(I_g^2 + \frac{4(2T-3)}{T\delta\theta^2}I_g + \frac{4N(2T-3)}{T^2\delta\theta^4}\right) \tag{49}$$

This is the expression provided by Kanitscheider et al. [2015a]. Unfortunately, it is a function of the true information $I_g$, which is unknown, such that the variance cannot be evaluated from data.

To find an unbiased estimate of this variance, note that the true information, $I_g$, shows up as $I_g$ and $I_g^2$. We already have an unbiased estimate of $I_g$, and will now derive such an unbiased estimate for $I_g^2$. Let us denote this estimate by $\left(\hat{I_g^2}\right)_{bc}$ (in contrast to the squared $\hat{I}_{g,bc}$, which is $\hat{I}_{g,bc}^2$). We find it by

$$\begin{aligned}\mathbb{E}\left((\hat{I}_{g,bc})^2\right) &= \mathrm{var}\left(\hat{I}_{g,bc}\right) + \mathbb{E}\left(\hat{I}_{g,bc}\right)^2\\ &= \frac{2}{2T-N-5}\left(I_g^2 + \frac{4(2T-3)}{T\delta\theta^2}I_g + \frac{4N(2T-3)}{T^2\delta\theta^4}\right) + I_g^2\\ &= \frac{1}{2T-N-5}\left((2T-N-3)I_g^2 + \frac{8(2T-3)}{T\delta\theta^2}I_g + \frac{8N(2T-3)}{T^2\delta\theta^4}\right).\end{aligned} \tag{50}$$

Solving for $I_g^2$ and substituting $I_g$ by its bias-corrected estimate $\hat{I}_{g,bc}$ reveals the bias-corrected estimate

$$\left(\hat{I_g^2}\right)_{bc} = \frac{2T-N-5}{2T-N-3}\hat{I}_{g,bc}^2 - \frac{1}{2T-N-3}\frac{8(2T-3)}{T\delta\theta^2}\hat{I}_{g,bc} - \frac{1}{2T-N-3}\frac{8N(2T-3)}{T^2\delta\theta^4}, \tag{51}$$

which satisfies $\mathbb{E}\left(\left(\hat{I_g^2}\right)_{bc}\right) = I_g^2$. Substituting the bias corrected estimates of $I_g$ and $I_g^2$ into Eq. (49) results after some algebra in the unbiased variance estimate

$$\operatorname{var}\left(\hat{I}_{g,bc}\right) = \frac{2}{2T-N-3}\left(\hat{I}_{g,bc}^2 + \frac{4(2T-3)}{T\delta\theta^2}\hat{I}_{g,bc} + \frac{4N(2T-3)}{T^2\delta\theta^4}\right), \tag{52}$$

which can be computed from data.

#### 1.5 Covariance of bias-corrected generalized Fisher information

As we are interested in how information scales with population size, we also need to know how information estimates for different subpopulations relate to each other. Knowing this relationship is essential to our model fits, as fitting the information scaling models to information estimates that are correlated across different population sizes could results in significant mis-estimates if these correlations are ignored. In fact, we will use the results from this section to show in Sec. 3.5 that the increase in information due to adding one more neuron to a population is uncorrelated across different subpopulations. Based on this insight, we thus fitted these information increases rather than absolute informations, as illustrated in Fig. 4 in the main text.

To identify the relation between the information estimates for different subpopulations, we will focus on two subpopulations with $N_x$ and $N_y$ neurons ($N_y \le N_x$) where the latter consists of a subset of neurons of the former. That is, the subpopulation with $N_x$ neurons contains all of the $N_y$ neurons in the (possibly) smaller subpopulation. We are interested in how their information estimates co-vary if we estimate both information measures from the same set of $T$ trials.

To find this covariance, let us decompose the true (i.e., non-empirical) moments of the larger subpopulation into

$$\delta\boldsymbol{f}_x = \begin{pmatrix}\delta\boldsymbol{f}_y \\ \delta\boldsymbol{f}_z\end{pmatrix}, \qquad \boldsymbol{\Sigma}_x = \begin{pmatrix}\boldsymbol{\Sigma}_y & \boldsymbol{\Sigma}_u \\ \boldsymbol{\Sigma}_u^T & \boldsymbol{\Sigma}_z\end{pmatrix}. \tag{53}$$

Here, $\delta\boldsymbol{f}_x$ and $\delta\boldsymbol{f}_y$ are the population tuning differences of the larger and smaller subpopulation, respectively, and we have ordered the neurons in the larger subpopulation such that it contains all shared neurons first, followed by all non-shared neurons. This re-ordering is possible, as the information estimates are independent or how neurons are ordered within a population. Furthermore, $\boldsymbol{\Sigma}_x$ and $\boldsymbol{\Sigma}_y$ are the noise covariance matrices of the larger and smaller subpopulation, and $\boldsymbol{\Sigma}_u$ is the the covariance of shared with non-shared neurons.

Experimentally, we cannot directly observe these moments, but instead estimate them through the empirical moments,

$$\delta\boldsymbol{\mu}_x = \begin{pmatrix}\delta\boldsymbol{\mu}_y \\ \delta\boldsymbol{\mu}_z.\end{pmatrix}, \qquad \boldsymbol{S}_x = \begin{pmatrix}\boldsymbol{S}_y & \boldsymbol{S}_u \\ \boldsymbol{S}_u^T & \boldsymbol{S}_z.\end{pmatrix} \tag{54}$$

Using the same properties as in the previous section, these empirical moments relate to the true moments by

$$\delta\boldsymbol{\mu}_x \sim \mathcal{N}\left(\boldsymbol{\delta f}_x, \frac{2}{T}\boldsymbol{\Sigma}_x\right), \qquad \boldsymbol{S}_x^{-1} \sim \mathcal{W}^{-1}\left(2(T-1)\boldsymbol{\Sigma}_x^{-1}, 2(T-1)\right), \tag{55}$$

$$\delta\boldsymbol{\mu}_y \sim \mathcal{N}\left(\boldsymbol{\delta f}_y, \frac{2}{T}\boldsymbol{\Sigma}_y\right), \qquad \boldsymbol{S}_y^{-1} \sim \mathcal{W}^{-1}\left(2(T-1)\boldsymbol{\Sigma}_y^{-1}, 2(T-1)\right), \tag{56}$$

The empirical covariances additionally have the properties [Muirhead, 2005]

$$(\boldsymbol{S}_z - \boldsymbol{S}_u\boldsymbol{S}_y^{-1}\boldsymbol{S}_u^T)^{-1} \sim \mathcal{W}^{-1}\left(2(T-1)(\boldsymbol{\Sigma}_z - \boldsymbol{\Sigma}_u\boldsymbol{\Sigma}_y^{-1}\boldsymbol{\Sigma}_u^T)^{-1}, 2(T-1)\right), \tag{57}$$

$$\boldsymbol{S}_u\boldsymbol{S}_y^{-1}|\boldsymbol{S}_y^{-1} \sim \mathcal{MN}_{N_y\times(N_x-N_y)}\left(\boldsymbol{\Sigma}_u\boldsymbol{\Sigma}_y^{-1}, \frac{1}{2(T-1)}(\boldsymbol{\Sigma}_x - \boldsymbol{\Sigma}_u\boldsymbol{\Sigma}_y^{-1}\boldsymbol{\Sigma}_u^T), \boldsymbol{S}_y^{-1}\right), \tag{58}$$

where $\mathcal{MN}$ is the matrix-normal distribution.

From Eq. (39), the bias-corrected generalized information for two subpopulations denoted as $I^x_{g,bc}$ and $I^y_{g,bc}$ can be written as

$$\hat{I}^x_{g,bc} = \frac{2T - N_x - 3}{2T - 2} \frac{\delta\boldsymbol{\mu}_x^T \boldsymbol{S}_x^{-1} \delta\boldsymbol{\mu}_x}{\delta\theta^2} - \frac{2N_x}{T\delta\theta^2}, \tag{59}$$

$$\hat{I}^y_{g,bc} = \frac{2T - N_y - 3}{2T - 2} \frac{\delta\boldsymbol{\mu}_y^T \boldsymbol{S}_y^{-1} \delta\boldsymbol{\mu}_y}{\delta\theta^2} - \frac{2N_y}{T\delta\theta^2}. \tag{60}$$

We can decompose $\hat{I}^x_{g,bc}$ into two terms. The first term is the shared information which is common between subpopulations $x$ and $y$ as both of them contains all of neurons in subpopulation $y$. The second term is the information gain that is gained by adding the non-shared neurons. This decomposition can be expressed as

$$\hat{I}^x_{g,bc} = \hat{I}^y_{g,bc} + \delta\hat{I}^{x-y}_{g,bc}, \tag{61}$$

where $\delta\hat{I}^{x-y}_{g,bc}$ is the information gain due to the non-shared components between subpopulations $x$ and $y$. The covariance of $\hat{I}^x_{g,bc}$ and $\hat{I}^y_{g,bc}$ is given by

$$\text{cov}\left(\hat{I}^x_{g,bc}, \hat{I}^y_{g,bc}\right) = \text{var}\left(\hat{I}^y_{g,bc}\right) + \text{cov}\left(\hat{I}^y_{g,bc}, \delta\hat{I}^{x-y}_{g,bc}\right), \tag{62}$$

where we already have expression for the variance on the right-hand side (i.e., Eq. (52)), and only need to find an expression for the covariance.

To calculate $\text{cov}\left(\hat{I}^y_{g,bc}, \delta\hat{I}^{x-y}_{g,bc}\right)$, let us first find an expression for $\delta\hat{I}^{x-y}_{g,bc}$. To find this expression, note that, by the decomposition of $\delta\boldsymbol{\mu}_x$ and $\boldsymbol{S}_x$, and using block matrix inversion,

$$\delta\boldsymbol{\mu}_x^T \boldsymbol{S}_x^{-1} \delta\boldsymbol{\mu}_x = \delta\boldsymbol{\mu}_y^T \boldsymbol{S}_y^{-1} \delta\boldsymbol{\mu}_y + \left(\delta\boldsymbol{\mu}_z - \boldsymbol{S}_u \boldsymbol{S}_y^{-1} \delta\boldsymbol{\mu}_y\right)^T \left(\boldsymbol{S}_z - \boldsymbol{S}_u \boldsymbol{S}_y^{-1} \boldsymbol{S}_u^T\right)^{-1} \left(\delta\boldsymbol{\mu}_z - \boldsymbol{S}_u \boldsymbol{S}_y^{-1} \delta\boldsymbol{\mu}_y\right), \tag{63}$$

Substituting this relationship into Eqs. (59) and (60) results in the bias-corrected information gain

$$\begin{aligned} \delta\hat{I}^{x-y}_{g,bc} &= \hat{I}^x_{g,bc} - \hat{I}^y_{g,bc} \\ &= \frac{N_y - N_x}{2T - N_y - 3} \hat{I}^y_{g,bc} + \frac{2T - N_x - 3}{2T - 2} \frac{\left(\delta\boldsymbol{\mu}_z - \boldsymbol{S}_u \boldsymbol{S}_y^{-1} \delta\boldsymbol{\mu}_y\right)^T \left(\boldsymbol{S}_z - \boldsymbol{S}_u \boldsymbol{S}_y^{-1} \boldsymbol{S}_u^T\right)^{-1} \left(\delta\boldsymbol{\mu}_z - \boldsymbol{S}_u \boldsymbol{S}_y^{-1} \delta\boldsymbol{\mu}_y\right)}{\delta\theta^2} + \text{const}, \end{aligned} \tag{64}$$

where "const" captures all non-stochastic terms that do not contribute to the covariance. Overall, this results in

$$\begin{aligned} \text{cov}\left(\hat{I}^y_{g,bc}, \delta\hat{I}^{x-y}_{g,bc}\right) = \frac{N_y - N_x}{2T - N_y - 3} \text{var}\left(\hat{I}^y_{g,bc}\right) + \frac{(2T - N_x - 3)(2T - N_y - 3)}{(2T-2)^2 \delta\theta^4} \\ \times \text{cov}\left(\delta\boldsymbol{\mu}_y^T \boldsymbol{S}_y^{-1} \delta\boldsymbol{\mu}_y, \left(\delta\boldsymbol{\mu}_z - \boldsymbol{S}_u \boldsymbol{S}_y^{-1} \delta\boldsymbol{\mu}_y\right)^T \left(\boldsymbol{S}_z - \boldsymbol{S}_u \boldsymbol{S}_y^{-1} \boldsymbol{S}_u^T\right)^{-1} \left(\delta\boldsymbol{\mu}_z - \boldsymbol{S}_u \boldsymbol{S}_y^{-1} \delta\boldsymbol{\mu}_y\right)\right), \end{aligned} \tag{65}$$

where we have substituted Eq. (60) for $I^y_{g,bc}$ to find the second term on the right-hand side. The first term of Eq. (65) is known from Eq. (52). The covariance expression in the second term can be expressed as

$$\begin{aligned} &\text{cov}\left(\delta\boldsymbol{\mu}_y^T \boldsymbol{S}_y^{-1} \delta\boldsymbol{\mu}_y, \left(\delta\boldsymbol{\mu}_z - \boldsymbol{S}_u \boldsymbol{S}_y^{-1} \delta\boldsymbol{\mu}_y\right)^T \left(\boldsymbol{S}_z - \boldsymbol{S}_u \boldsymbol{S}_y^{-1} \boldsymbol{S}_u^T\right)^{-1} \left(\delta\boldsymbol{\mu}_z - \boldsymbol{S}_u \boldsymbol{S}_y^{-1} \delta\boldsymbol{\mu}_y\right)\right) \\ &\quad = \mathbb{E}\left(\delta\boldsymbol{\mu}_y^T \boldsymbol{S}_y^{-1} \delta\boldsymbol{\mu}_y \left(\delta\boldsymbol{\mu}_z - \boldsymbol{S}_u \boldsymbol{S}_y^{-1} \delta\boldsymbol{\mu}_y\right)^T \left(\boldsymbol{S}_z - \boldsymbol{S}_u \boldsymbol{S}_y^{-1} \boldsymbol{S}_u^T\right)^{-1} \left(\delta\boldsymbol{\mu}_z - \boldsymbol{S}_u \boldsymbol{S}_y^{-1} \delta\boldsymbol{\mu}_y\right)\right) \\ &\quad - \mathbb{E}\left(\delta\boldsymbol{\mu}_y^T \boldsymbol{S}_y^{-1} \delta\boldsymbol{\mu}_y\right) \mathbb{E}\left(\left(\delta\boldsymbol{\mu}_z - \boldsymbol{S}_u \boldsymbol{S}_y^{-1} \delta\boldsymbol{\mu}_y\right)^T \left(\boldsymbol{S}_z - \boldsymbol{S}_u \boldsymbol{S}_y^{-1} \boldsymbol{S}_u^T\right)^{-1} \left(\delta\boldsymbol{\mu}_z - \boldsymbol{S}_u \boldsymbol{S}_y^{-1} \delta\boldsymbol{\mu}_y\right)\right) \end{aligned} \tag{66}$$

First we evaluate the last expectation in Eq. (66) which is

$$\mathbb{E}\left(\left(\delta\boldsymbol{\mu}_z - \boldsymbol{S}_u \boldsymbol{S}_y^{-1} \delta\boldsymbol{\mu}_y\right)^T \left(\boldsymbol{S}_z - \boldsymbol{S}_u \boldsymbol{S}_y^{-1} \boldsymbol{S}_u^T\right)^{-1} \left(\delta\boldsymbol{\mu}_z - \boldsymbol{S}_u \boldsymbol{S}_y^{-1} \delta\boldsymbol{\mu}_y\right)\right) \tag{67}$$

Conditioned on $\boldsymbol{S}_y^{-1}$, we observe that $(\boldsymbol{S}_z - \boldsymbol{S}_u\boldsymbol{S}_y^{-1}\boldsymbol{S}_u^T)^{-1}$ is independent of $\boldsymbol{S}_u\boldsymbol{S}_y^{-1}$ [Muirhead, 2005]. Thus we can first take the expectation of $(\boldsymbol{S}_z - \boldsymbol{S}_u\boldsymbol{S}_y^{-1}\boldsymbol{S}_u^T)^{-1}$ to get

$$\frac{2T-2}{2T-N_x-3}\left(\delta\boldsymbol{\mu}_z - \boldsymbol{S}_u\boldsymbol{S}_y^{-1}\delta\boldsymbol{\mu}_y\right)^T\left(\boldsymbol{\Sigma}_z - \boldsymbol{\Sigma}_u\boldsymbol{\Sigma}_y^{-1}\boldsymbol{\Sigma}_u^T\right)^{-1}\left(\delta\boldsymbol{\mu}_z - \boldsymbol{S}_u\boldsymbol{S}_y^{-1}\delta\boldsymbol{\mu}_y\right). \tag{68}$$

Next, we observe that $\boldsymbol{S}_u\boldsymbol{S}_y^{-1}|\boldsymbol{S}_y^{-1}$ is matrix normal, which has a simple expression for the expectation of its quadratic form. Using this expression yields

$$\frac{2T-2}{2T-N_x-3}\left(\delta\boldsymbol{\mu}_z - \boldsymbol{\Sigma}_u\boldsymbol{\Sigma}_y^{-1}\delta\boldsymbol{\mu}_y\right)^T\left(\boldsymbol{\Sigma}_z - \boldsymbol{\Sigma}_u\boldsymbol{\Sigma}_y^{-1}\boldsymbol{\Sigma}_u^T\right)^{-1}\left(\delta\boldsymbol{\mu}_z - \boldsymbol{\Sigma}_u\boldsymbol{\Sigma}_y^{-1}\delta\boldsymbol{\mu}_y\right) + \frac{N_x-N_y}{2T-N_x-3}\delta\boldsymbol{\mu}_y^T\boldsymbol{S}_y^{-1}\delta\boldsymbol{\mu}_y \tag{69}$$

We do not need to complete the expectation over the remaining random variables because most involved terms cancel out each other later on.

Utilizing the same strategy we evaluate the expectation of the first term in Eq. (66) which is given by

$$\mathbb{E}\left(\delta\boldsymbol{\mu}_y^T\boldsymbol{S}_y^{-1}\delta\boldsymbol{\mu}_y\left(\delta\boldsymbol{\mu}_z - \boldsymbol{S}_u\boldsymbol{S}_y^{-1}\delta\boldsymbol{\mu}_y\right)^T\left(\boldsymbol{S}_z - \boldsymbol{S}_u\boldsymbol{S}_y^{-1}\boldsymbol{S}_u^T\right)^{-1}\left(\delta\boldsymbol{\mu}_z - \boldsymbol{S}_u\boldsymbol{S}_y^{-1}\delta\boldsymbol{\mu}_y\right)\right). \tag{70}$$

Its expectation with respect to $\left(\boldsymbol{S}_z - \boldsymbol{S}_u\boldsymbol{S}_y^{-1}\boldsymbol{S}_u^T\right)^{-1}$ is

$$\frac{2T-2}{2T-N-3}\delta\boldsymbol{\mu}_y^T\boldsymbol{S}_y^{-1}\delta\boldsymbol{\mu}_y\left(\delta\boldsymbol{\mu}_z - \boldsymbol{S}_u\boldsymbol{S}_y^{-1}\delta\boldsymbol{\mu}_y\right)^T\left(\boldsymbol{\Sigma}_z - \boldsymbol{\Sigma}_u\boldsymbol{\Sigma}_y^{-1}\boldsymbol{\Sigma}_u^T\right)^{-1}\left(\delta\boldsymbol{\mu}_z - \boldsymbol{S}_u\boldsymbol{S}_y^{-1}\delta\boldsymbol{\mu}_y\right). \tag{71}$$

The expectation with respect to $\boldsymbol{S}_u\boldsymbol{S}_y^{-1}|\boldsymbol{S}_y^{-1}$ is given by

$$\frac{2T-2}{2T-N_x-3}\delta\boldsymbol{\mu}_y^T\boldsymbol{S}_y^{-1}\delta\boldsymbol{\mu}_y\left(\delta\boldsymbol{\mu}_z - \boldsymbol{\Sigma}_u\boldsymbol{\Sigma}_y^{-1}\delta\boldsymbol{\mu}_y\right)^T\left(\boldsymbol{\Sigma}_z - \boldsymbol{\Sigma}_u\boldsymbol{\Sigma}_y^{-1}\boldsymbol{\Sigma}_u^T\right)^{-1}\left(\delta\boldsymbol{\mu}_z - \boldsymbol{\Sigma}_u\boldsymbol{\Sigma}_y^{-1}\delta\boldsymbol{\mu}_y\right) + \frac{N_x-N_y}{2T-N_x-3}\left(\delta\boldsymbol{\mu}_y^T\boldsymbol{S}_y^{-1}\delta\boldsymbol{\mu}_y\right)^2 \tag{72}$$

Utilizing the fact that $\delta\boldsymbol{\mu}_y$ and $\delta\boldsymbol{\mu}_z - \boldsymbol{\Sigma}_u\boldsymbol{\Sigma}_y^{-1}\delta\boldsymbol{\mu}_y$ are jointly Gaussian and uncorrelated, which means they are independent, we can combine Eqs. (72) and (69) to simplify the expression in Eq. (66) to

$$\begin{aligned}
&\operatorname{cov}\left(\delta\boldsymbol{\mu}_y^T\boldsymbol{S}_y^{-1}\delta\boldsymbol{\mu}_y, \left(\delta\boldsymbol{\mu}_z - \boldsymbol{S}_u\boldsymbol{S}_y^{-1}\delta\boldsymbol{\mu}_y\right)^T\left(\boldsymbol{S}_z - \boldsymbol{S}_u\boldsymbol{S}_y^{-1}\boldsymbol{S}_u^T\right)^{-1}\left(\delta\boldsymbol{\mu}_z - \boldsymbol{S}_u\boldsymbol{S}_y^{-1}\delta\boldsymbol{\mu}_y\right)\right) \\
&= \frac{N_x-N_y}{2T-N_x-3}\left(\mathbb{E}\left(\left(\delta\boldsymbol{\mu}_y^T\boldsymbol{S}_y^{-1}\delta\boldsymbol{\mu}_y\right)^2\right) - \mathbb{E}\left(\delta\boldsymbol{\mu}_y^T\boldsymbol{S}_y^{-1}\delta\boldsymbol{\mu}_y\right)^2\right) \\
&= \frac{N_x-N_y}{2T-N_x-3}\operatorname{var}\left(\delta\boldsymbol{\mu}_y^T\boldsymbol{S}_y^{-1}\delta\boldsymbol{\mu}_y\right) \\
&= \frac{(N_x-N_y)(2T-2)^2\delta\theta^4}{(2T-N_x-3)(2T-N_y-3)^2}\operatorname{var}\left(\hat{I}_{g,bc}^y\right).
\end{aligned} \tag{73}$$

Substituting Eq. (73) into Eq. (65) results in

$$\operatorname{cov}\left(\hat{I}_{g,bc}^y, \delta\hat{I}_{g,bc}^{x-y}\right) = \frac{N_y-N_x}{2T-N_y-3}\operatorname{var}\left(\hat{I}_{g,bc}^y\right) + \frac{N_x-N_y}{2T-N_y-3}\operatorname{var}\left(\hat{I}_{g,bc}^y\right) = 0, \tag{74}$$

which means that information in the smaller population is uncorrelated to the information gain obtained from non-shared neurons. As a consequence,

$$\operatorname{cov}\left(\hat{I}_{g,bc}^x, \hat{I}_{g,bc}^y\right) = \operatorname{var}\left(\hat{I}_{g,bc}^y\right). \tag{75}$$

Note the this only holds for the bias-corrected information estimates. For the naïve estimates, a similar derivation shows that $\hat{I}_{g,nv}^y$ and $\delta\hat{I}_{g,nv}^{x-y}$ are correlated.

### 2 Information scaling models

We assume that information in the recorded population is limited by the presence of information-limiting correlations [Moreno-Bote et al., 2014]. In this case, the noise covariance matrix $\boldsymbol{\Sigma}_N$ for a population of $N$ neurons decomposes into

$$\boldsymbol{\Sigma}_N = \boldsymbol{\Sigma}_{0,N} + \frac{1}{I_\infty} \boldsymbol{f}'_N \boldsymbol{f}'^T_N, \tag{76}$$

where $\boldsymbol{\Sigma}_{0,N}$ is the non-limiting covariance component, $I_\infty$ is the asymptotic information, and $\boldsymbol{f}'_N$ is the derivative of the mean population activity. All of these quantities depend on the stimulus, $\theta$, but we will keep this dependency implicit for notational convenience. In the $N \to \infty$ limit, only the second component limits information, while the information associated with $\boldsymbol{\Sigma}_{0,N}$ grows without bounds.

To see how information grows in the presence of information-limiting correlations, note that the Sherman-Morrison formula allows us to express $\boldsymbol{\Sigma}_N^{-1}$ by

$$\boldsymbol{\Sigma}_N^{-1} = \boldsymbol{\Sigma}_{0,N}^{-1} - \frac{\boldsymbol{\Sigma}_{0,N}^{-1} \boldsymbol{f}'_N \boldsymbol{f}'^T_N \boldsymbol{\Sigma}_{0,N}^{-1}}{I_\infty + \boldsymbol{f}'^T_N \boldsymbol{\Sigma}_{0,N}^{-1} \boldsymbol{f}'_N} \tag{77}$$

Let us denote the linear Fisher information associated with the non-limiting component by $I_{0,N} = \boldsymbol{f}'^T_N \boldsymbol{\Sigma}_{0,N}^{-1} \boldsymbol{f}'_N$. Then, after some algebra, the total Fisher information is given by

$$I_N = \boldsymbol{f}'^T_N \boldsymbol{\Sigma}_N^{-1} \boldsymbol{f}'_N = \boldsymbol{f}'^T_N \boldsymbol{\Sigma}_{0,N}^{-1} \boldsymbol{f}'_N - \frac{\boldsymbol{f}'^T_N \boldsymbol{\Sigma}_{0,N}^{-1} \boldsymbol{f}'_N \boldsymbol{f}'^T_N \boldsymbol{\Sigma}_{0,N}^{-1} \boldsymbol{f}'_N}{I_\infty + \boldsymbol{f}'^T_N \boldsymbol{\Sigma}_{0,N}^{-1} \boldsymbol{f}'_N} = \frac{1}{\frac{1}{I_{0,N}} + \frac{1}{I_\infty}}, \tag{78}$$

or, equally, $I_N^{-1} = I_{0,N}^{-1} + I_\infty^{-1}$. This result forms the core of our information scaling models. For the remainder of this section we will discuss how we would expect information $\boldsymbol{I}_{0,N}$ in the non-limiting component to scale, and the impact of measurement noise on overall information scaling.

#### 2.1 Linear non-limiting information scaling for large $N$

To characterize the scaling of $I_{0,N}$ with $N$, let us use the spectral decomposition

$$\boldsymbol{\Sigma}_{0,N} = \sum_{n=1}^N \sigma_{N,n}^2 \boldsymbol{z}_{N,n} \boldsymbol{z}_{N,n}^T, \tag{79}$$

with variances $\sigma_{N,1}^2, \dots, \sigma_{N,N}^2$ and principal directions $\boldsymbol{z}_{N,1}, \dots, \boldsymbol{z}_{N,N}$. Then, $I_{0,N}$ is given by

$$I_{0,N} = \sum_{n=1}^N \frac{\left(\boldsymbol{f}'^T_N \boldsymbol{z}_{N,n}\right)^2}{\sigma_{N,n}^2} = \|\boldsymbol{f}'_N\|^2 \sum_{n=1}^N \frac{\cos^2(\alpha_{N,n})}{\sigma_{N,n}^2}, \tag{80}$$

where $\alpha_{N,n}$ is the angle between $\boldsymbol{f}'_N$ and $\boldsymbol{z}_n$.

To see how $I_{0,N}$ scales with $N$, let us assume that the $\alpha_{N,n}$'s are independent of the $\sigma_{N,n}^2$'s. Furthermore, $f'_{N,n}$ (that is, the $n$th component of $\boldsymbol{f}'_N$) is $\mathcal{O}(1)$, such that $\|\boldsymbol{f}'_N\|^2$ will be $\mathcal{O}(N)$. In addition, geometry requires that $\sum_{n=1}^N \cos^2(\alpha_{N,n}) = 1$, such that each $\cos^2(\alpha_{N_n})$ is $\mathcal{O}(1/N)$ [Moreno-Bote et al., 2014]. Together, this yields

$$I_{0,N} \propto N \sum_{n=1}^N \frac{1}{N} \frac{1}{\sigma_{N,n}^2} = \sum_{n=1}^N \frac{1}{\sigma_{N,n}^2}. \tag{81}$$

Therefore, under these assumptions, the scaling of $I_{0,N}$ only depends on the scaling of the eigenvalue spectrum $\{\sigma_{N,1}^2, \dots, \sigma_{N,N}^2\}$ of $\boldsymbol{\Sigma}_{0,N}$.

For the following, we will assume that each neuron in the population features some small amount of "private" noise that is not correlated with the variability of other neurons. This private noise introduces a lower bound,

$\sigma_0^2$, on the variances, that is $\sigma_{N,n}^2 \geq \sigma_0^2$ for all $n$. Together with the above expression, this allows us to derive a lower bound on the scaling of non-limiting information. In particular, by Jensen's inequality

$$I_{0,N} \propto N \left( \frac{1}{N} \sum_{n=1}^{N} \frac{1}{\sigma_{N,n}^2} \right) \geq N \frac{1}{\frac{1}{N} \sum_{n=1}^{N} \sigma_{N,n}^2} \propto N. \tag{82}$$

The second-to-last expression contains the average variance, which is lower-bounded by $\sigma_0^2$ and of order one. Therefore, the scaling of $I_{0,N}$ is at least $\mathcal{O}(N)$.

To gain further insight into the scaling of $I_{0,N}$, assume a sequence of non-limiting covariance matrices $\mathbf{\Sigma}_{0,M}, \mathbf{\Sigma}_{0,M-1}, \mathbf{\Sigma}_{0,M-2}, \dots$, starting with some large population with $M$ neurons. Each consecutive matrix $\mathbf{\Sigma}_{0,N-1}$ is constructed from the next-larger matrix $\mathbf{\Sigma}_{0,N}$ by removing a single neuron, such that they share all entries except for one row and column associated with that neuron. If we order their eigenvalues according to $\sigma_{N,1}^2 \geq \sigma_{N,2} \geq \dots \sigma_{N,N}^2$ and $\sigma_{N-1,1}^2 \geq \sigma_{N-1,2}^2 \geq \dots \geq \sigma_{N-1,N-1}^2$, it is known that these eigenvalues obey the interleaved ordering

$$\sigma_{N,1}^2 \geq \sigma_{N-1,1}^2 \geq \sigma_{N,2}^2 \geq \sigma_{N-1,2}^2 \geq \dots \sigma_{N-1,N-1}^2 \geq \sigma_{N,N}^2. \tag{83}$$

Using $I_{0,N} \propto \sum_{n=1}^{N} \sigma_{N,n}^{-2}$, the information increase when moving from $N-1$ to $N$ neurons becomes

$$I_{0,N} - I_{0,N-1} \propto \sum_{n=1}^{N-1} \left( \frac{1}{\sigma_{N,n}^2} - \frac{1}{\sigma_{N-1,n}^2} \right) + \frac{1}{\sigma_{N,N}^2}. \tag{84}$$

This information increase is $\mathcal{O}(1)$ if both terms on the left-hand side are $\mathcal{O}(1)$.

The second term is $\mathcal{O}(1)$ if there exists some positive constant $C$ such that, for all $N$ above some $N_0$, $\sigma_{N,N}^{-2} \leq C$. As $\sigma_{N,N}^2$ is always the smallest eigenvalue of the covariance matrix, this implies that $\mathcal{O}(1)$ can be guaranteed as long as $\sigma_{N,N}^2$ remains positive with increasing $N$, which is satisfied by our previous assumption that each neuron has some private noise. If it instead would go to zero, we would have $\lim_{N \to M} \sigma_{N,N}^{-2} = 0$, violating the requirement.

For the first term we observe that the hierarchical eigenvalue relationship of nested matrices implies that $\sigma_{N,n}^{-2} \leq \sigma_{N-1,n}^{-2}$ for all $n = 1, \dots, N-1$. This implies that every element in the sum is negative. However, the information increase $I_{0,N} - I_{0,N-1}$ cannot be negative. Therefore, the second term on the left-hand side has to be at least as large as the negative first term (i.e., the sum), that is

$$\frac{1}{\sigma_{N,N}^2} \geq - \sum_{n=1}^{N} \left( \frac{1}{\sigma_{N,n}^2} - \frac{1}{\sigma_{N-1,n}^2} \right). \tag{85}$$

As $\sigma_{N,N}^{-2}$ is $\mathcal{O}(1)$, the sum cannot be larger than $\mathcal{O}(1)$. Overall, as long as none of the variances become zero with increasing $N$, the increase in $I_{0,N}$ will be $\mathcal{O}(1)$, which implies that $I_{0,N}$ scales with $\mathcal{O}(N)$.

#### 2.2 Models for $I_{0,N}$

The above argument shows that, under rather general conditions, $I_{0,N}$ can be expected to scale with $\mathcal{O}(N)$. However, it does not tell us about how $I_{0,N}$ behaves for small $N$, which depends on the details of the structure of $\mathbf{\Sigma}_{N,0}$.

To describe the details of this structure, we compared two models for $I_{0,N}$. The first, called the *lim* model, directly follows the scaling results and assumes that $I_{0,N} = cN$ with some parameter $c$ that is independent of $N$. The second model, called the *lim-exp* model, allows the non-limiting information to initially grow supralinearly before converging to a linear growth. We derived this model by integrating $c\left(1 - e^{-N/\tau}\right)$ from zero to $N$, resulting in

$$I_{0,N} = c\left(N + \tau\left(e^{-\frac{N}{\tau}} - 1\right)\right), \tag{86}$$

with the additional parameter $\tau$ that controls the extent of the initial supralinearity (in units of $N$). We have chosen this particular model, as it turns out easier to fit that alternative models (such as, for example, integrating a re-scaled logistic sigmoid over the positive half-line) that provide qualitatively similar qualitative $I_{0,N}$ scaling. This model approaches $I_{0,N} = cN$ in the $\tau \to 0$ limit. Model comparison revealed the *lim* model to significantly outperform the *lim-exp* model (Fig. S1), such that we focused on the *lim* model in the main text.

##### 2.3 Impact of measurement noise

Our recordings of neural activity might be noisy, introducing additional variability into our estimates of $\mathbf{\Sigma}_N$ and $\boldsymbol{f}'_N$. To estimate the effect of such measurement noise, we assume it to be of equal magnitude and independent across neurons, such that it adds an additional diagonal term to the covariance decomposition,

$$\mathbf{\Sigma}_N = \mathbf{\Sigma}_{0,N} + \frac{1}{I_\infty}\boldsymbol{f}'_N\boldsymbol{f}'^T_N + \sigma^2_{rec}\boldsymbol{I}, \tag{87}$$

where $\sigma^2_{rec}$ denotes the variance of the measurement noise. We don't assume it to impact differential correlations, as those limit information *in the brain*, rather than our measurement thereof.

Following the same derivation as in the beginning of this section, the information in a population of $N$ neurons becomes

$$I_N = \frac{1}{\frac{1}{I_{0,rec,N}} + \frac{1}{I_\infty}}, \tag{88}$$

where

$$I_{0,rec,N} = \boldsymbol{f}'^T_N \left(\mathbf{\Sigma}_{0,N} + \sigma^2_{rec}\boldsymbol{I}\right)^{-1} \boldsymbol{f}'_N, \tag{89}$$

is the non-limiting information, including measurement noise. We can, as before, use the spectral decomposition $\mathbf{\Sigma}_{0,N} = \sum_{n=1}^N \sigma^2_{N,n}\boldsymbol{z}_n\boldsymbol{z}_n^T$ and observe that $\boldsymbol{I} = \sum_{n=1}^N \boldsymbol{z}_n\boldsymbol{z}_n^T$ , resulting in

$$\mathbf{\Sigma}_{0,N} + \sigma^2_{rec}\boldsymbol{I} = \sum_{n=1}^N \left(\sigma^2_{N,n} + \sigma^2_{rec}\right)\boldsymbol{z}_n\boldsymbol{z}_n^T. \tag{90}$$

This shows that measurement noise increases all eigenvalues of $\mathbf{\Sigma}_{0,N}$ by the same magnitude.

This has several consequences. First, the added variance baseline results in $I_{0,rec,N}$ to grow more slowly with $N$ than $I_{0,N}$. Second, this baseline causes in the eigenvalues of $\mathbf{\Sigma}_{0,N} + \sigma^2_{rec}\boldsymbol{I}$ to be more similar to each other than those of $\mathbf{\Sigma}_{0,N}$ alone. As a consequence, the growth of $I_{0,rec,N} \propto \sum_{n=1}^N \left(\sigma^2_{N,n} + \sigma^2_{rec}\right)^{-1}$ with $N$ is more linear than that of $I_{0,N} \propto \sum_{n=1}^N \sigma^{-2}_{N,n}$. This might make $I_{0,rec,N} = c_{rec}N$ a good model of non-limiting information growth, even if $I_{0,N} = cN$ is not. Third, as the measurement noise impacts only $I_{0,rec,N}$ but not $I_\infty$, measurement noise only impacts our estimates of $c$ but not of $I_\infty$. Fourth, measurement noise will lower our estimates of $c$, and therefore increase our estimates of $N_a = a/(1-a)I_\infty/c$, which is the population size at which a fraction $a$ of the asymptotic information $I_\infty$ is reached.

#### 3 Estimating the information scaling moments from neural data

Here, we fix the discrimination (i.e., the pair of drift directions, $\theta_1$ and $\theta_2$) and discuss how we estimate the moments of Fisher information for different population sizes. To do so, we assume a large population with $M$ neurons of which we subsample $N$ neurons, and where $N \ll M$. Rather than focusing on the moments of the Fisher information $I_n$ for population size $n \leq N$, we will instead focus on the moments of the Fisher information increase, $\Delta I_n = I_n - I_{n-1}$ (with $I_0 = 0$), when increasing the population size from $n-1$ to $n$ neurons, for reasons that become apparent later. Our aim is to estimate the mean, $\mathbb{E}\left(\Delta I_n\right)$, the variance, var $\left(\Delta I_n\right)$, and the covariance, cov $\left(\Delta I_n, \Delta I_m\right)$, for different population sizes $n$ and $m$.

##### 3.1 Generative model and desired moments

To describe the stochasticity of $\Delta I_n$, we assume the following generative process. Assume that neurons in the large population have indices 1 to $M$, and that we uniformly draw a subset of $N$ different neurons with indices $i_1, i_2, \ldots, i_N$, denoted $i_{1:N}$. This subpopulation has moments $\boldsymbol{f}'_{i_{1:N}}$ and $\boldsymbol{\Sigma}_{i_{1:N}}$, that in turn can be used to compute its associated Fisher information. However, we do not directly observe these moments, but instead record the population activity across $T$ trials for each stimulus, $\theta_1$ and $\theta_2$, from which we compute the empirical moments $\boldsymbol{\gamma}_{i_{1:N}}$ and $\boldsymbol{\Omega}_{i_{1:N}}$. These empirical moments are in turn used to compute the Fisher information increases $\Delta\hat{I}_{1:N}$, using the bias-corrected estimates discussed further above. In summary, the generative process follows the Markov chain

$$i_{1:N} \rightarrow \boldsymbol{f}'_{i_{1:N}}, \boldsymbol{\Sigma}_{i_{1:N}} \rightarrow \boldsymbol{\gamma}_{i_{1:N}}, \boldsymbol{\Omega}_{i_{1:N}} \rightarrow \Delta\hat{I}_{1:N}. \tag{91}$$

In this Markov chain, the first and last transition are deterministic, and the center transition is stochastic. Therefore, we can write the generative model as

$$p\left(\Delta\hat{I}_{1:N}\right) = \sum_{i_{1:N}} p\left(\Delta\hat{I}_{1:N}\left(\boldsymbol{\gamma}_{i_{1:N}}, \boldsymbol{\Omega}_{i_{1:N}}\right) | i_{1:N}\right) p\left(i_{1:N}\right), \tag{92}$$

where the Fisher information increases are a deterministic function of the empirical moments, and the sum is over different subpopulations drawn from the larger population. We assume these draws to be uniform, that is $p\left(i_{1:N}\right) \propto 1$.

To find the moments of $\Delta\hat{I}_n$, we use iterated expectation, variance, and covariance, which, for a Markov chain $Z \rightarrow X_1, X_2$ is given by

$$\mathbb{E}_{X_1}\left(X_1\right) = \mathbb{E}_Z\left(\mathbb{E}_{X_1|Z}\left(X_1\right)\right), \tag{93}$$

$$\mathrm{var}_{X_1}\left(X_1\right) = \mathbb{E}_Z\left(\mathrm{var}_{X_1|Z}\left(X_1\right)\right) + \mathrm{var}_Z\left(\mathbb{E}_{X_1|Z}\left(X_1\right)\right), \tag{94}$$

$$\mathrm{cov}_{X_1,X_2}\left(X_1, X_2\right) = \mathbb{E}_Z\left(\mathrm{cov}_{X_1,X_2|Z}\left(X_1, X_2\right)\right) + \mathrm{cov}_Z\left(\mathbb{E}_{X_1|Z}\left(X_1\right), \mathbb{E}_{X_2|Z}\left(X_2\right)\right). \tag{95}$$

Applied to our generative model, that yields the decompositions

$$\mathbb{E}_{\Delta\hat{I}_n}\left(\Delta\hat{I}_n\right) = \mathbb{E}_{i_{1:N}}\left(\mathbb{E}_{\Delta\hat{I}_n|i_{1:N}}\left(\Delta\hat{I}_n\right)\right), \tag{96}$$

$$\mathrm{var}_{\Delta\hat{I}_n}\left(\Delta\hat{I}_n\right) = \mathbb{E}_{i_{1:N}}\left(\mathrm{var}_{\Delta\hat{I}_n|i_{1:N}}\left(\Delta\hat{I}_n\right)\right) + \mathrm{var}_{i_{1:N}}\left(\mathbb{E}_{\Delta\hat{I}_n|i_{1:N}}\left(\Delta\hat{I}_n\right)\right), \tag{97}$$

$$\mathrm{cov}_{\Delta\hat{I}_n,\Delta\hat{I}_m}\left(\Delta\hat{I}_n, \Delta\hat{I}_m\right) = \mathbb{E}_{i_{1:N}}\left(\mathrm{cov}_{\Delta\hat{I}_n,\Delta\hat{I}_m|i_{1:N}}\left(\Delta\hat{I}_n, \Delta\hat{I}_m\right)\right) + \mathrm{cov}_{i_{1:n}}\left(\mathbb{E}_{\Delta\hat{I}_n|i_{1:N}}\left(\Delta\hat{I}_n\right), \mathbb{E}_{\Delta\hat{I}_m|i_{1:N}}\left(\Delta\hat{I}_m\right)\right), \tag{98}$$

where both variance and covariance are decomposed into (i) the (co)variance of the information increase for a fixed subpopulation $i_{1:N}$, averaged across different subpopulations, and (ii) how the average information increase for a fixed subpopulation (co)varies across different subpopulations.

Our data does not allow us to directly estimate these moments for two reasons. First, we don't observe the larger population, and so can't use it to draw different subpopulations from this larger population. We will address how we handle this limitation in the next subsection. Second, we only observe a single set of empirical moments, $\boldsymbol{\mu}$ and $\boldsymbol{S}$, for the subpopulation that we record from. We will address how we handle this limitation in the remaining subsections.

##### 3.2 Simulating samples from a large, unobserved population

Our generative model assumes that we are subsampling $N$ neurons from a large neural populations of $M$ neurons. Our data, in contrast, are population recordings from a single neural population with $N$ neurons. To use these recordings to simulate sampling from various subpopulations of the larger population, we assume these subpopulations to be statistically similar to the recorded population. That is, the different sampled subpopulations will contain neurons with similar activity statistics as the recorded population. Thus, each sampled subpopulation will contain all neurons from the recorded population, but in a different order for each sampled subpopulation. We will simulate this by introducing a new index set $j_{1:N} = j_1, j_2, \ldots, j_N$ that, for each sampled subpopulation $i_{1:N}$, contains a random order of the indices $1, \ldots, N$ of neurons in the recorded population. With this, all of the above moments across $i_{1:N}$ will become moments across $j_{1:N}$, while taking into account that the recorded subpopulation is used as a proxy for sampling different subpopulations from a larger populations. We will describe the consequences of this for each of the moments separately.

#### 3.3 Estimating the mean

The desired mean of the information increase $\Delta\hat{I}_n$ is, by Eq. (96) the average information increase for a particular set of empirical moments, $\gamma_{i_{1:N}}$ and $\boldsymbol{\Omega}_{i_{1:N}}$, for a particular subpopulation $i_{1:N}$, averaged across different subpopulations. We deal with not being able to sample different subpopulations by replacing $i_{1:N}$ by a randomly ordered recorded population $j_{1:N}$. Furthermore, we cannot draw different empirical moments, $\gamma_{i_{1:N}}$ and $\boldsymbol{\Omega}_{i_{1:N}}$ for a given subpopulation, as would be required to compute $\mathbb{E}_{\Delta\hat{I}|i_{1:N}}\left(\Delta\hat{I}_n\right)$. We will replace this expectation with our best estimate thereof, which is the Fisher information increase estimate based on the bias-correctet Fisher information, estimated from the empirical moments of the recorded population, $\boldsymbol{\mu}$ and $\boldsymbol{S}$. Overall, this leads to the approximate estimate,

$$\mathbb{E}_{i_{1:N}}\left(\mathbb{E}_{\Delta\hat{I}_n|i_{1:N}}\left(\Delta\hat{I}_n\right)\right) \approx \mathbb{E}_{j_{1:N}}\left(\Delta\hat{I}_n\left(\boldsymbol{\mu}_{j_{1:N}}, \boldsymbol{S}_{j_{1:N}}\right)\right), \tag{99}$$

where $\boldsymbol{\mu}_{j_{1:N}}$ and $\boldsymbol{S}_{j_{1:N}}$ denote the empirical moments with neurons ordered according to $j_{1:N}$. As our Fisher information estimate is unbiased, the above estimate will be unbiased as well. In practice, we approximate the expectation over $j_{1:N}$ by 10000 random ordering.

#### 3.4 Estimating the variance

The variance, Eq. (97), is decomposed into two terms. The first, $\mathbb{E}_{i_{1:N}}\left(\mathrm{var}_{\Delta\hat{I}_n|i_{1:N}}\left(\Delta\hat{I}_n\right)\right)$, is the variance of the Fisher information increase for a fixed subpopulation, averaged across many subpopulations. This term captures the uncertainty in $\Delta\hat{I}_n$ due to using the empirical moments to estimate it. The second term, given $\mathrm{var}_{i_{1:N}}\left(\mathbb{E}_{\Delta\hat{I}_n|i_{1:N}}\left(\hat{I}_n\right)\right)$, captures how the average Fisher information increase for a given subpopulation varies across different subpopulations. Our data doesn't allow us to compute either of these terms directly. However, it turns out that they are both well-approximated by how the Fisher information increase estimated from the empirical moments, $\boldsymbol{\mu}$ and $\boldsymbol{S}$, varies across different population orders, $j_{1:N}$, that is

$$\mathbb{E}_{i_{1:N}}\left(\mathrm{var}_{\Delta\hat{I}_n|i_{1:N}}\left(\Delta\hat{I}_n\right)\right) + \mathrm{var}_{i_{1:N}}\left(\mathbb{E}_{\Delta\hat{I}_n|i_{1:N}}\left(\Delta\hat{I}_n\right)\right) \approx \mathrm{var}_{j_{1:N}}\left(\Delta\hat{I}_n\left(\boldsymbol{\mu}_{j_{1:N}}, \boldsymbol{S}_{j_{1:N}}\right)\right). \tag{100}$$

To understand why this approximation works, we need to consider two components that contribute to the empirical moments of the recorded neurons. The first is that, for each neuron and each neuron pair, these empirical moments are noisy, as they are estimated from a limited number of trials. Thus, we can approximate the effect of using empirical rather than true moments, as captured by the first term in Eq. (97), by computing the variance across different neurons in the population, as achieved by the variance across different orderings, $j_{1:N}$. The second factor is that different neurons contribute different amounts of information to the population. This comes into play in the second term in Eq. (97), and is again well-approximated by the variance across different orderings, $j_{1:N}$. As it seems paradoxical that the same variance can capture both kinds of effects at the same time, we have demonstrated it in simulations of neural populations, shown in Fig. S7a.

#### 3.5 Estimating the covariance

As the variance, the covariance, Eq. (98) can be decomposed into two terms that capture different sources of uncertainty. The first term, $\mathbb{E}_{i_{1:N}}\left(\mathrm{cov}_{\Delta\hat{I}_n,\Delta\hat{I}_m|i_{1:N}}\left(\Delta\hat{I}_n, \Delta\hat{I}_m\right)\right)$, captures the uncertainty associated with estimating empirical moments from a limited number of trials. To find this covariance, assume $n \neq m$ and note that,

$$\begin{aligned}\text{cov}\left(\Delta\hat{I}_n, \Delta\hat{I}_m\right) &= \text{cov}\left(\hat{I}_n - \hat{I}_{n-1}, \hat{I}_m - \hat{I}_{m-1}\right) \\ &= \text{cov}\left(\hat{I}_n, \hat{I}_m\right) - \text{cov}\left(\hat{I}_n, \hat{I}_{m-1}\right) - \text{cov}\left(\hat{I}_{n-1}, \hat{I}_m\right) + \text{cov}\left(\hat{I}_{n-1}, \hat{I}_{m-1}\right)\end{aligned} \tag{101}$$

where all covariances are conditional on $i_{1:N}$. Without loss of generality we can assume that $n > m$, and use Eq. (75) from Sec. 1.4 to find

$$\text{cov}\left(\Delta\hat{I}_n, \Delta\hat{I}_m\right) = \text{var}\left(\hat{I}_m\right) - \text{var}\left(\hat{I}_{m-1}\right) - \text{var}\left(\hat{I}_m\right) + \text{var}\left(\hat{I}_{m-1}\right) = 0. \tag{102}$$

This shows, that, conditional on $i_{1:N}$, the information increase estimates are uncorrelated.

The second term, $\text{cov}_{i_{1:N}}\left(\mathbb{E}_{\Delta\hat{I}_n|i_{1:N}}\left(\Delta\hat{I}_n\right), \mathbb{E}_{\Delta\hat{I}_m|i_{1:N}}\left(\Delta\hat{I}_m\right)\right)$, captures how the average Fisher information increase associated with adding the $n$th neuron correlates with that when adding the $m$th neuron across different subpopulation samples. On average, these increases will be negatively correlated, for the following reason. The variance of the information estimate $\hat{I}_n = \sum_{k=1}^{n} \Delta\hat{I}_k$ can be decomposed into

$$\text{var}\left(\hat{I}_n\right) = \sum_{k=1}^{n}\left(\text{var}\left(\Delta\hat{I}_k\right) + 2\sum_{l=1}^{k-1}\text{cov}\left(\Delta\hat{I}_k, \Delta\hat{I}_l\right)\right), \tag{103}$$

which shows the impact of the individual variances, as well as the covariance between estimates associated with different population sizes. For a population of $M$ neurons, the estimate of total information, $\hat{I}_M$, will be the same, irrespective of how the neurons are ordered within that subset. Therefore, var $\left(\hat{I}_M\right) = 0$. However, as, by definition, var $\left(\Delta\hat{I}_n\right) \geq 0$, the above decomposition implies that the covariances need to be on average negative, to ensure that the sum of variances and covariances becomes zero.

The same principle applies if we estimate the variance of $\Delta\hat{I}_n$ by shuffling the order, $j_{1:N}$, of neurons in a smaller, recorded population. If this population has $N$ neurons, then var $\left(\mathbb{E}_{\hat{I}_N|j_{1:N}}\left(\hat{I}_N\right)\right) = 0$, irrespective of $j_{1:N}$, such that the information increase estimates will be negatively correlated.

Recall that we use population order shuffling as a proxy for repeatedly subsampling $N$ neurons from a larger population of $M$ neurons. The shuffling-induced negative correlations arise from using the same $N$ recorded neurons across all estimates. If we instead subsample a larger population, the different sampled subpopulations are bound to share a smaller number of neurons. For two subpopulations that share no neurons, these estimates would be completely uncorrelated. However, even for $N \ll M$, two random subpopulations of size $N$ are likely to share neurons of the larger population. Indeed, the same intuition underlying the birthday paradox [Suzuki et al., 2006] tells us that we are almost guaranteed to find such shared neurons. However, the correlations don't only depend on the presence of shared neurons, but also on how many of them are shared, and the latter will decrease significantly for larger $M$. To show that this significantly lowers the impact of negative correlations on the total variance, we compare this variance computed with and without accounting for these correlations for different $M$'s. As Fig. S7b shows, their impact drops significantly with growing $M$. Therefore, we will approximate them to be zero, that is

$$\text{cov}_{\Delta\hat{I}_n,\Delta\hat{I}_m}\left(\Delta\hat{I}_n, \Delta\hat{I}_m\right) \approx 0. \tag{104}$$

This results in an overestimate of the variance of the Fisher information estimate, and make our fits less certain, and, as a consequence, more conservative.

### 4 Population activity models

We used two different models to simulate population activity, as described below.

#### 4.1 A Gaussian population activity model with limited information

We used a simple Gaussian activity model to satisfies the assumptions of Gaussianity underlying the generalized linear Fisher information, and to test some of the properties of our estimates. This model violates some properties of neural activity, like non-negativity, but is convenient for our purposes, as it supports fine control over the eigenvalues of $\mathbf{\Sigma}_0$, the alignment of $\boldsymbol{f}'$ to $\mathbf{\Sigma}_0$, and the asymptotic information, $I_\infty$. For a population size of $N$ neurons, we generated $\mathbf{\Sigma}_0$ by drawing a random orthonormal matrix $\boldsymbol{Z}_0$ of size $N \times N$ that forms the eigenvectors of $\mathbf{\Sigma}_0$. We parameterized the eigenvalues by $\sigma_{n,0}^2 = \sigma_0^2 + \sigma_b m^{-\beta}$, which together form the diagonal matrix $\boldsymbol{D}_0 = \mathrm{diag}\left(\sigma_{0,1}^2, \dots, \sigma_{0,N}^2\right)$. $\boldsymbol{Z}_0$ and $\boldsymbol{D}_0$ together specify $\mathbf{\Sigma}_0$ by $\mathbf{\Sigma}_0 = \boldsymbol{Z}_0\boldsymbol{D}_0\boldsymbol{Z}_0^T$. For a given $\boldsymbol{f}'$, the full noise covariance is then given by $\mathbf{\Sigma} = \mathbf{\Sigma}_0 + I_\infty^{-1}\boldsymbol{f}'\boldsymbol{f}'^T$.

For Fig. S2, we drew a random $\boldsymbol{f}' \sim \mathcal{N}(\mathbf{0}, \boldsymbol{I})$, and subsequently rescaled the vector such that $\|\boldsymbol{f}'\| = g$. This makes the alignment of $\boldsymbol{f}'$ to the eigenvectors of $\mathbf{\Sigma}_0$ roughly uniform on average. For this figure, we use parameters $I_\infty = 20$ (or $I_\infty = \infty$ for the unlimited-information case), $g = 20$, $\sigma_0^2 = 10^{-3}$, $\sigma_b^2 = 1$, and $\beta = 0.1$.

For Fig. S7, we specified the alignment of $\boldsymbol{f}'$ to the eigenvectors of $\mathbf{\Sigma}_0$ by $\alpha_n = \sigma_\alpha^2 + \propto e^{-n/\tau_\alpha}$, normalized such that $\sum_n \alpha_n = 1$. This yields $\tilde{\boldsymbol{f}}' = \sum_n \alpha_n \boldsymbol{z}_n$ ($\boldsymbol{z}_n$ is the $n$th eigenvector of $\mathbf{\Sigma}_0$), and $\boldsymbol{f}' = \sqrt{g_f N}\tilde{\boldsymbol{f}}'/\|\tilde{\boldsymbol{f}}'\|$. The magnitude of $\boldsymbol{f}'$ here scales with $\sqrt{N}$ to ensure roughly similar information across different $N$'s. The used parameters were $I_\infty = 100$, $g_f = 0.008$, $\sigma_0^2 = 5 \times 10^{-5}$, $\sigma_g^2 = 3$, $\beta = 0.5$, $\sigma_\alpha^2 = 10^{-3}$, and $\tau_\alpha = 30$, which results in population statistics comparable to those shown in Figs. 3 and 8 in the main text.

#### 4.2 A visual hierarchy population activity model

We relied on [Kanitscheider et al., 2015b] for a more realistic model of V1 population activity that is driven by pixel-level inputs. Details of this model can be found in Kanitscheider et al. [2015b]. Briefly, a population of $N$ neurons responded to a $P \times P$ pixelated images $\boldsymbol{J}$ of an oriented Garbor. The $n$th neuron's linear filter $\boldsymbol{F}_n$ was for each $(x, y)$ pixel determined by

$$ce^{-\frac{(x^2+y^2)}{2\sigma^2}} \cos\left(\frac{2\pi x}{\lambda}\cos(\theta_n) + \frac{2\pi y}{\lambda}\sin(\theta_n) + \phi\right), \tag{105}$$

where $c$ is the Michelson contrast, $\theta_n$ determines the neuron's tuning, $\sigma^2$ determines the size of the exponential envelope, and $\lambda$ and $\phi$ are the Garbor's frequency and phase, respectively. The filter was computed by the above function for each $(x, y)$ and then standardized to have mean zero and unit variance across all $(x, y)$. Image templates, $\boldsymbol{J}(\theta)$, in response to stimulus $\theta$ were generated equally, with $\theta_n$ replaced by the template's orientation, $\theta$. Each neuron's gain, $a_n$, was drawn from a log-normal distribution with unit mean and variance $\sigma_a^2$, and then multiplied by the overall gain, $g$.

Neural population activity is assumed to arise from the image template with Gaussian pixel noise (zero mean, variance $\sigma_0^2$), followed by application of the per-neuron linear filters, $\boldsymbol{F}_n$, multiplied by their gain $a_n$, and a Poisson step. For Fig. S2, we estimated information from a set of trials, in each of which neural activity was generated from a different pixel noise instantiation. For Fig. S5, we skipped the Poisson step, as it introduced additional noise and was not required for the point we were trying to make. Instead, we estimated Fisher information from approximations to the neural mean responses and their covariance matrix, following Kanitscheider et al. [2015b]. We computed the mean response of neuron $n$ to image $\boldsymbol{J}$ by $f_n(\theta) = \left[a_n \sum_{xy} F_{n,xy} J_{xy}(\theta)\right]_+$, where $[\cdot]_+$ is the threshold-linear function that sets negative values to zero. The population noise covariance was computed by

$$\mathbf{\Sigma}(\theta) = \sigma_0^2\left(\boldsymbol{a}\boldsymbol{a}^T\right) \otimes \left[\boldsymbol{F}^T\boldsymbol{F}\right]_+ + \mathrm{diag}\left(\boldsymbol{a} \otimes \boldsymbol{f}(\theta)\right), \tag{106}$$

where $\otimes$ denotes the (element-wise) Hadamard product, $\boldsymbol{a} = (a_1, \dots, a_N)^T$ is the column vector of per-neuron gains, $\boldsymbol{F}$ is the $P^2 \times N$ filter matrix with per-neuron filters unrolled as vectors along its columns, and $\boldsymbol{f}(\theta)$ is the mean population activity in response to stimulus $\theta$. The information was computed from $\mathbf{\Sigma}(\theta)$ and $\boldsymbol{f}(\theta)$.

For Fig. S2 we used the parameters $\sigma = P/5$, $\lambda = P/1.5$, $\phi = 0$, $c = 1$, $g = 20$, and $\sigma_a = \sqrt{2}$, as in Kanitscheider et al. [2015b], and additionally $N = 2500$, $P = 32$ and $\sigma_0 = 0.25$. To simulate infinite information, we removed pixel noise by setting $\sigma_0 = 0$. For Fig. S5, we used the same parameters except $N = 1000$, $g = 10$, and $\sigma_0 =$

$0.11$, to achieve the desired level of information, and approximate information saturation within the simulated population size. In all simulations, neural tuning, $\theta_n$, was uniformly distributed over $[-\pi, \pi]$, and pixels $(x, y)$ were uniformly distributed over locations $[-(P-1)/2, (P-1)/2]$ in both dimensions.

### References

Rubén Moreno-Bote, Jeffrey Beck, Ingmar Kanitscheider, Xaq Pitkow, Peter Latham, and Alexandre Pouget. Information-limiting correlations. *Nature Neurosience*, 17(10):1410–1417, 2014. doi: 10.1038/nn.3807.

Deep Ganguli and Eero P. Simoncelli. Efficient sensory encoding and bayesian inference with heterogeneous neural populations. *Neural Computation*, 26:2103–2134, 2014. doi: 10.1162/NECO_a_00638.

Thomas M. Cover and Joy A. Thomas. *Elements of Information Theory*. Wiley Series in Telecommunications and Signal Processing. Whiley-Interscience, 2nd edition, 2006.

Wei Ji Ma, Jeffrey M Beck, Peter E Latham, and Alexandre Pouget. Bayesian inference with probabilistic population codes. *Nature Neuroscience*, 9:1432–1438, 2006. doi: 10.1038/nn1790.

James O. Berger. *Statistical decision theory and Bayesian analysis*. Springer Series in Statistics. Springer, 2nd edition, 1993.

David Marvin Green and John A. Swets. *Signal detection theory and psychophysics*. Peninsula Pub, 1989.

Bruno B. Averbeck and Daeyeol Lee. Effects of noise correlations on informatino encoding and decoding. *Journal of Neurophysiology*, 95:3633–3644, 2006. doi: 10.1152/jn.00919.2005.

Yuzhi Chen, Wilson S. Geisler, and Eyal Seidemann. Optimal decoding of correlated neural population responses in the primate visual cortex. *Nature Neuroscience*, 9(11):1412–1420, 2006. doi: 10.1038/nn1792.

Ramon Nogueira, Nicole E. Peltier, Akiyuki Anzai, Gregory C. DeAngelis, Julio Martínez-Trujillo, and Rubén Moreno-Bote. The effects of population tuning and trial-by-trial variability on information encoding and behavior. *The Journal of Neuroscience*, 0859-19, 2019. doi: 10.1523/JNEUROSCI.0859-19.2019.

James O. Berger. *Pattern recognition and machine learning*. Information Science and Statistics. Springer, 2011.

Ingmar Kanitscheider, Ruben Coen-Cagli, Adam Kohn, and Alexandre Pouget. Measuring fisher information accurately in correlated neural populations. *PLoS Computational Biology*, 11(6):e1004218, 2015a. doi: 10.1371/journal.pcbi.1004218.

Richard A. Johnson and Dean W. Wichern. *Applied multivariate statistical analysis*. Pearson, 5th edition, 2007.

Robb J. Muirhead. *Aspects of multivariate statistical theory*. Wiley, 2nd edition, 2005.

Kazuhiro Suzuki, Dongvu Tonien, Kaoru Kurosawa, and Koji Toyota. Birthday paradox for multi-collisions. In Min Surp Rhee and Byoungcheon Lee, editors, *Information Security and Cryptology — ICISC 2006*, pages 29–40. Springer-Verlag Berlin Heidelberg, 2006. doi: 10.1007/11927587_5.

Ingmar Kanitscheider, Ruben Coen-Cagli, and Alexandre Pouget. Origin of information-limiting noise correlations. *Proceedings of the National Academy of Sciences*, 112(50):E6973–E6982, 2015b. doi: 10.1073/pnas.1508738112.

### 5 Supplementary Tables

| | | | Session | | | | | | |
|---|---|---|---|---|---|---|---|---|---|
| Mouse | Contrast | Avg. | A | B | C | D | E | F | G |
| 1 | 10% | $0.12 \pm 0.02$ | 0.14 | 0.11 | | | | | |
| 2 | 10% | $0.07 \pm 0.03$ | 0.11 | 0.04 | | | | | |
| 3 | 10% | $0.16 \pm 0.02$ | 0.22 | 0.20 | 0.09 | 0.13 | 0.14 | | |
| 4 | 10% | $0.12 \pm 0.01$ | 0.10 | 0.11 | 0.12 | 0.10 | 0.13 | 0.13 | 0.16 |
| 5 | 10% | $0.22 \pm 0.06$ | 0.12 | 0.16 | 0.20 | 0.40 | | | |
| | 25% | $0.23 \pm 0.03$ | 0.19 | 0.20 | 0.22 | 0.32 | | | |
| 6 | 10% | $0.20 \pm 0.02$ | 0.16 | 0.23 | 0.22 | | | | |
| | 25% | $0.22 \pm 0.01$ | 0.20 | 0.23 | 0.24 | | | | |

Table S1: Average Fisher information per neuron in $rad^{-2}/neuron$, across all sessions/mice, averaged across all $\delta\theta = 45^\circ$ discriminations. The average Fisher information was computed from the Fisher information scaling for trial-shuffled data that removed across-neuron correlations. For individual neurons, it can be computed by $2\left(\langle r|\theta_1\rangle - \langle r|\theta_2\rangle\right)^2 / \left(\delta\theta^2\left(\text{var}\left(r|\theta_1\right) + \text{var}\left(r|\theta_2\right)\right)\right)$, where $r|\theta_j$ is the neural response to stimulus $\theta_j$. The *Avg.* column provides the average across sessions (mean $\pm$ 1 SEM).

### 6 Supplementary Figures

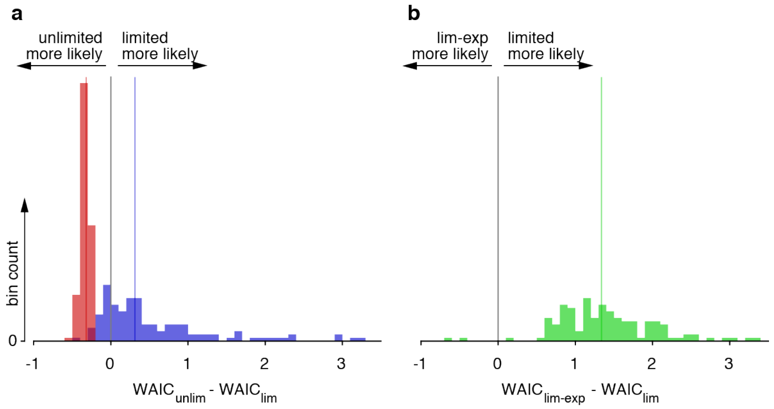

Figure S1: Model comparison of different information scaling models. Both panels show histograms of differences in the Watanabe-Akaike Information Criterion (WAIC) for two different models fitted to the measured information scaling curves across all eight discriminations with $\delta\theta = 45°$, sessions, and mice. (a) shows the WAIC difference for fitting a model that assumes no information limitation (*unlim*) to one that does (*lim*), for regular (blue) and shuffled (red) data. For regular data this difference is in most cases positive, indicating that the information-limiting model fits the data better. In fact, even for individual negative WAIC differences, the average across all eight WAIC difference within a session remains positive. For shuffled data, a model assuming no information limitation fits the data better in all instances. This confirms that our model comparison is not biased towards the model assuming limited information. (b) shows the WAIC difference for fitting two models that assume limited information (see Sec. 2.2), one with linear scaling of the non-limiting component (*lim*), and one assuming initial supralinear scaling of that component (*lim-exp*). The latter only fits the data better in few instances. In those, the average WAIC difference across all discriminations within that session is nonetheless positive. The colored lines in (a) and (b) show the median WAIC difference across all comparisons.

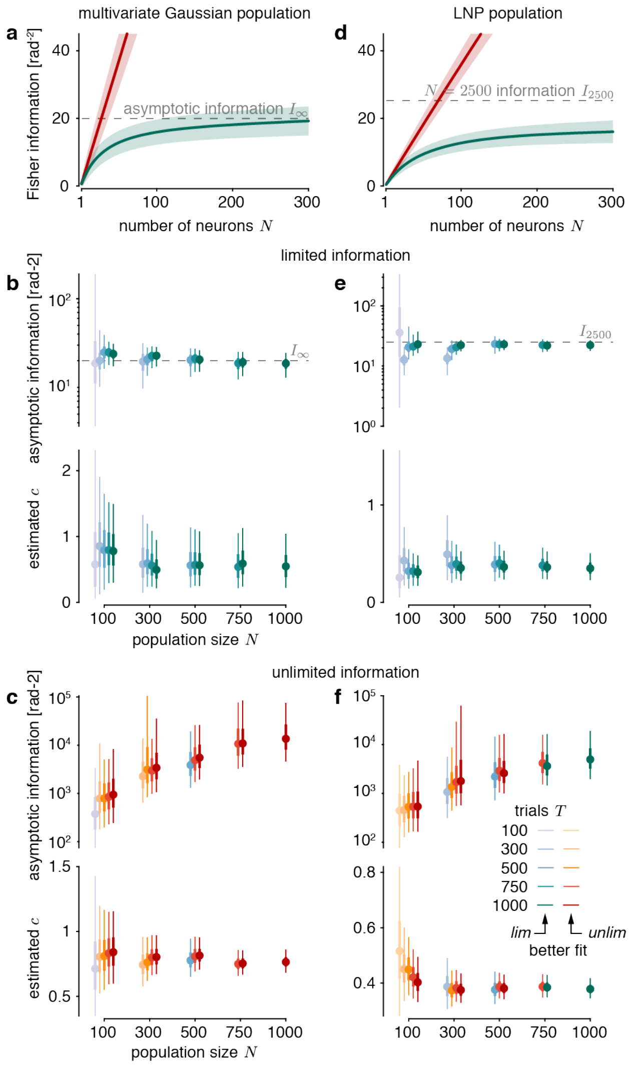

Figure S2: Recovering asymptotic information from simulated population activity. We simulated neural population activity, using either a multivariate Gaussian population model (a-c; see Sec. 4.1 for details) or a linear-nonlinear Poisson model (d-f; see Sec. 4.2 for details) and fitted a linear scaling (*unlim*) and a limited information scaling model (*lim*). For each model type, we generated two large datasets (limited information and unlimited information; $\delta\theta = \theta_2 - \theta_1 = 45°$ in both cases) and then subsampled neurons and trials to perform the fits. (a,d) Example information scaling for $N = 300$ and $T = 500$ (mean $\pm$ 1SD information estimation; green/red = limited/unlimited information). For the Gaussian model we could specify the asymptotic information $I_\infty$ (dashed grey line). For the LNP model we estimated it from the information $I_{2500}$ at $N = 2500$ neurons. (b-c,e-f) Estimated asymptotic information and non-limiting information scaling for the *lim* model from data with different population sizes $N$ and numbers of trials $T$ per stimulus. The posterior estimates are shown as in Fig. 4c in the main text. Blue/green and orange/red colors indicate a better fit by the *lim* and *unlim* model (WAIC for model comparison), respectively. Asymptotic information is well-estimated by the *lim* model (b,e), and more certain for larger $N$ and $T$. Model comparison in most cases (28 out of 30 for Gaussian model, 26 out of 30 for LNP model) correctly identifies if information was limited or unlimited (colors).

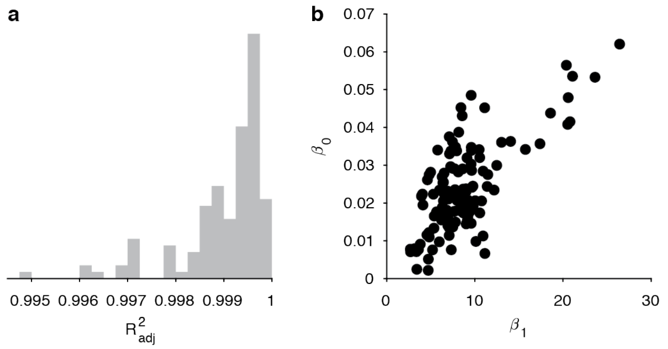

Figure S3: Statistics of a linear fit $I_N^{-1} = \beta_0 + \beta_1 N^{-1}$ across all eight discriminations with $\delta\theta = 45°$, sessions, and mice. (a) The adjusted $R^2$ is close to one for all fits. (b) Both intercept, $\beta_0$, and slope, $\beta_1$, are significantly above zero for all discriminations. The plot shows these intercepts with 95% CIs, which are obscured by the dots.

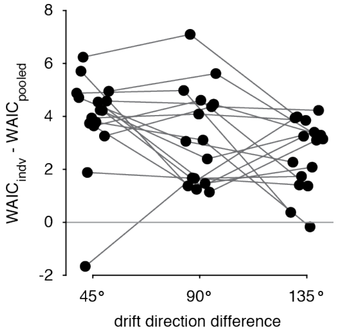

Figure S4: Model comparison of per-discrimination fits vs. pooled fits across multiple discriminations. The figure shows for each session (individual sessions connected by grey lines; horizontally jittered for clarity) the WAIC difference of fitting the information scaling of individual discriminations (*indv*) vs. fitting all of these discriminations simultaneously (*pooled*). The mostly positive WAIC differences, preferring pooled fits, confirm that the information scaling across different discriminations with the same drift direction difference $\delta\theta$ were exceedingly similar. The tested discriminations were $45°$ vs. $90°$, $135°$ vs. $180°$, $225°$ vs. $270°$, and $315°$ vs. $0°$ ($\delta\theta = 45°$); $45°$ vs. $135°$, $90°$ vs. $180°$, $225°$ vs. $315°$, and $270°$ vs. $0°$ ($\delta\theta = 90°$); and $45°$ vs. $180°$, $90°$ vs. $315°$, and $225°$ vs. $0°$ ($\delta\theta = 135°$). The WAIC differences for $\delta\theta = 315°$ had overall smaller magnitudes, as they pooled across three rather than four discriminations.

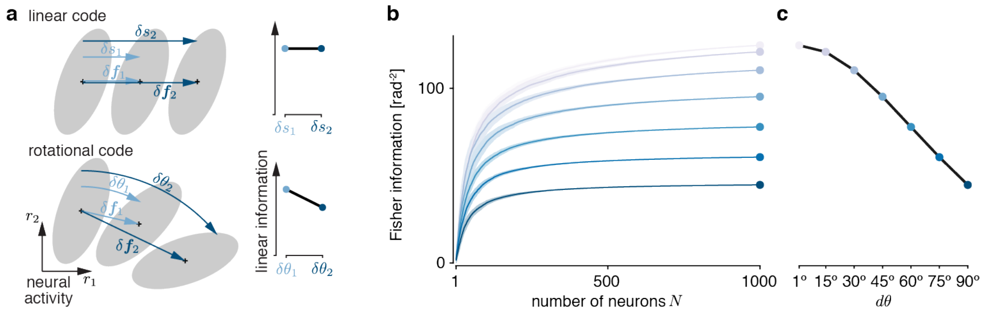

Figure S5: Linear Fisher information is expected to drop with increasing $\delta\theta$. (a) Generalized linear Fisher information measures how easy it is to discriminate two stimuli from the population responses they evoke. This discriminabiliy is measured by the performance of a linear discriminator, normalized by the stimulus difference ($\delta s$ or $\delta\theta$). For population responses (dots = mean population activity for one stimulus, shaded areas = 1SD of the noise covariance; $\delta \boldsymbol{f}_i$ = difference in mean population activity for different $\delta s_i$ / $\delta\theta_i$) whose mean response changes linearly with the stimulus $s$, this information remains unchanged when $\delta s$ changes (top; $\delta s_1$ vs. $\delta s_2$). Population activity that encodes a circular stimulus $\theta$ is bound to violate this linearity, and its associated linearly decodable information drops with an increase in $\delta\theta$ (bottom; $\delta\theta_1$ vs. $\delta\theta_2$). This occurs also if a non-linear decoder that accounts for the circularity of $\theta$ would recover the same information, irrespective of $\delta\theta$, and is not a bug of the linear decoder, which nonetheless correctly identifies all linearly decodable information (that drops with $\delta\theta$). (b) We demonstrate this effect by simulating V1 population in response to oriented Garbor pattern, and estimate the information encoded about their orientation. We show how information grows with population sizes for stimulus pairs with different $\delta\theta$ (colors; mean $\pm$ 1SD across different orders with which neurons are added to the population). (c) The information at $N = 1000$, which we use as a proxy for $I_\infty$, drops with $\delta\theta$, for the reason illustrated for the rotational code in (a). Details of the simulations to generate (b) and (c) are described in Sec. 4.2. The simulations quantify information about oriented Garbor pattern rather than the drift direction of drifting gratings, and so should only be qualitatively compared to the data in the main text.

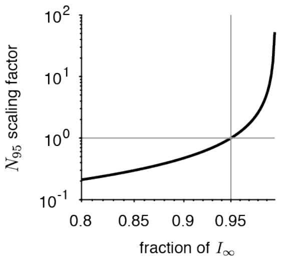

Figure S6: The scaling of the estimated population size with the fraction of asymptotic information. Let $N_a$ denote the population size required to encode $a$% of the total asymptotic information, $I_\infty$. Changing $a$ results in a simple re-scaling of $N_a$. This figures illustrates this re-scaling for different $a$, using $N_{95}$ as a base measure. For example, if we would be interested in $N_{90}$ instead of $N_{95}$, we would read off the scaling factor for 90%, and would re-scale the reported $N_{95}$ to get estimates for $N_{90}$.

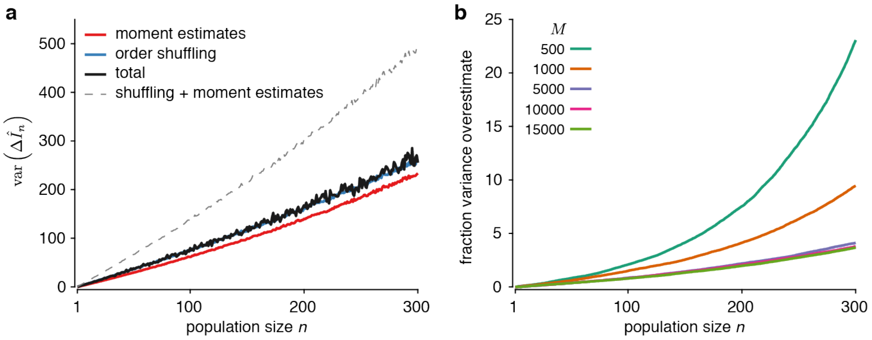

Figure S7: The variance and covariance of Fisher information scaling. We simulated virtual populations of different sizes $M$ as described in Sec. 4.1, yielding $\boldsymbol{f}'$ and $\boldsymbol{\Sigma}$ for each $M$. (a) To demonstrate that the variance in the Fisher information increase estimate due to shuffling well-approximates the combined variance due to population subsampling and due to estimating the moments from a finite number of trials, we generated one population with $M = 10,000$ neurons. We in turn drew 100 empirical moments, $\gamma \sim \mathcal{N}\left(\boldsymbol{f}', 2\boldsymbol{\Sigma}/(T\delta\theta)^2\right)$ and $\boldsymbol{\Omega} \sim \mathcal{W}\left(\boldsymbol{\Sigma}/(2T-2), 2T-1\right)$, corresponding to estimating these moments from $T = 1,000$ trials each for two drift directions separated by $\delta\theta = 45°$. We additionally subsampled $N = 300$ neurons of the full population ten times, resulting in ten $i_{1:N}$ neuron indices, and, for each $i_{1:N}$, containing a fixed set of neurons, shuffled their order 100 times, resulting in 100 $j_{1:N}$ per $i_{1:N}$. For the empirical moments, we computed the Fisher information increase for each subsampled, shuffled population $j_{1:N}$, resulting in $10^6$ estimates for each population size $n \in 1, \ldots N$. The figure shows the variance due to shuffling only (blue, averaged over different subsamples and empirical moments), and due to empirical moments only (red, averaged over different subsamples and shuffles). As comparison, we computed the total variance of the same estimate across 100 subsampled populations with $N = 300$ neurons for each set of empirical moment (black; variance across $10^4$ estimates), which is the variance we aim to estimate. As the plot shows, the variance due to shuffling well-approximates this total variance. A naïve sum of the variance due to empirical moments and shuffling (grey dashed) would over-estimate the total variance. (b) To estimate the degree by which the variance of the Fisher information increase, $\text{var}\left(\Delta\hat{I}_n\right)$, is overestimated when ignoring the negative correlations across different $\Delta\hat{I}_n$'s, we generate populations of different sizes, $M$, and their associated moments. For each population, we then estimated the covariance $\text{cov}\left(\Delta\hat{I}_n, \Delta\hat{I}_m\right)$ across 1,000 different subsamples $i_{1:N}$ of populations of $N = 300$ neurons. In turn, we estimated the Fisher information variance once when taking into account this covariance, $\text{var}\left(\hat{I}_n\right) = \sum_{j=1}^{n}\left(\text{var}\left(\Delta\hat{I}_j\right) + 2\sum_{k=1}^{j-1}\text{cov}\left(\Delta\hat{I}_k, \Delta\hat{I}_j\right)\right)$, and once when not doing so, $\tilde{\text{var}}\left(\hat{I}_n\right) = \sum_{j=1}^{n}\text{var}\left(\Delta\hat{I}_j\right)$. The plot shows the resulting fraction $\left(\tilde{\text{var}}\left(\Delta\hat{I}_n\right) - \text{var}\left(\Delta\hat{I}_n\right)\right)/\text{var}\left(\Delta\hat{I}_n\right)$ for different $n$ and $M$ as an average across ten different generated populations, and shows that the variance overestimate becomes smaller for larger populations.
